# Listening to the bats of Carajás: Applied bioacoustics for species inventory and environment use in a mosaic of forests, savannas, and industrial mining in the Brazilian Amazonia

**DOI:** 10.1101/2024.08.23.609443

**Authors:** Lidiane Gomes, Enrico Bernard

## Abstract

Bats emit echolocation calls for orientation, foraging, and social interactions. These calls are mostly species-specific, reliable for inventories and to assess habitat use, characteristics useful for large, species-rich but poorly sampled areas. This is the case of Carajás, in Brazilian Amazonia, a mosaic of cave-rich dense forests and unique metalophilous savannas (known as *canga*), harboring a rich bat fauna but also industrial iron ore mining, stressing the need to preserve biodiversity. We used bioacoustics (142,000 minutes of recording) to inventory bats at 61 points in Carajás and identified 43 sonotypes of seven bat families, including species rarely recorded with capture nets. Eleven species were recorded for the first time in Carajás. Species richness varied among environments – forests being the richest – but *cangas* had greater richness stability and a more distinct species composition. All areas with imminent mining had high bat richness. Richness in a post-mined area increased, possibly indicating resilience of some species. By providing a reference sound library for bats in Carajás, we proved the usefulness of biacoustics to improve the environmental licensing processes involving mining in biodiversity-rich areas, useful not only for Amazonia but also for other tropical environments with high bat species richness.

## INTRODUCTION

Bats participate in several ecological interactions and play important roles in plant pollination, seed dispersal, and insect predation, providing valuable ecosystem services (Fleming and Muchhala 2008; Fleming et al. 2009; Kunz et al. 2011; Ramírez-Fráncel et al. 2022). Because their species and ecological richness, and different levels of tolerance to changes in the landscape, bats are also considered indicators of environmental quality and impacts of human activities (Medellín et al. 2000; Schulze et al. 2000; Jones et al. 2009; Struebig et al. 2013; Russo et al. 2021; Di Ponzio et al. 2023). However, to use bats as environmental indicators, they must first be correctly recorded and identified (Arias-Aguilar et al. 2018; Sugai et al. 2019). In this sense, bats stand out for their remarkable ability to produce high-frequency sounds for spatial orientation, which is called echolocation (Fenton et al. 2016). Echolocation allows bats to detect, locate, and classify objects, helping with spatial orientation and foraging; however, it also makes these animals conspicuous and their calls recordable (Schnitzler et al. 2001; Blumstein et al. 2011; Sugai et al. 2019). Furthermore, most echolocation signals are species-specific and reliable for the correct identification of the emitting species (e.g. O’Farrell and Gannon 1999; Arias-Aguilar et al. 2018). These characteristics provide interesting possibilities for the use of echolocation calls for inventorying, identifying, and recording the use of the environment by bats (Schnitzler et al. 2003; MacSwiney et al., 2008; Denzinger and Schnitzler 2013; Marques et al. 2016; Silva and Bernard 2017; Appel et al. 2021). Such an approach can be particularly useful in large, species-rich but poorly sampled areas (Blumstein et al. 2011; Sugai et al. 2019).

Brazil is home to high bat species richness, with 186 species recorded (Garbino et al. 2024). In Brazil, the Amazon region accounts for around 76% of these species (Delgado-Jaramillo et al. 2020). However, most information on bats in the Brazilian Amazonia comes from a small number of in-depth inventories conducted in a few specific locations (Bernard et al. 2011; Delgado-Jaramillo et al. 2020). Moreover, a fraction of those bat inventories used bioacoustics sampling (Barnett et al. 2006; Appel et al. 2021; Di Ponzio et al. 2023). Thus, large areas with very favorable habitats and expected high biodiversity were not sampled for this group, pointing that the actual bat species richness in Amazonia is likely greater than what is already known and estimated (Tavares et al. 2024). However, despite its enormous biodiversity, several parts of the Brazilian Amazonia are under strong human pressure, with high rates of habitat loss and degradation (Aguiar et al. 2020; Lapola et al. 2023). This alarming scenario indicates that parts of the rich Amazonian biodiversity may be lost without scientific documentation (Carvalho et al. 2023).

The region of Carajás, in the southeast of Pará state, is a mosaic from dense forest areas to a more open special vegetation, called *canga*, on the mountain tops of ferruginous substrates, with several endemic species (Viana et al. 2016; Mota et al. 2018). In addition to being home to high biodiversity, Carajás is one of the largest mineral deposits on the planet, with very relevant iron ore mines, which means that this region simultaneously experiences pressure for mineral exploration and the need to preserve its unique biodiversity. Due to its geological formation, Carajás also houses a high concentration of caves (Piló et al. 2015; ICMBio 2016) and such a combination of high richness and diversity of habitats proved positive for bat species as well. Until recently, 75 species of bats belonging to 46 genera and eight families were known to Carajás (Tavares et al. 2012), and species distribution modeling indicated a high potential for another 20-25 species in the region (do Amaral et al. 2023). However, the available bat sampling for Carajás relied on mist nets and/or captures in caves, which generated a biased sampling towards species of the Phyllostomidae family (Tavares et al. 2012), whose capture is more frequent, but is poorly detected acoustically (Silva and Bernard 2017; Appel et al. 2021; Carvalho et al. 2022).

Mining is a high-impact activity, that can be destructive to the environment in which it occurs (Maus et al. 2022; Aska et al. 2024). In regions of high biodiversity, such as Amazonia, the licensing of industrial mining requires special attention (e.g. Sonter et al. 2018). In fact, bats play a key-role in the environmental licensing of mineral activities in Brazil: caves in mining areas must be classified as having maximum, high, medium, or low relevance, and at least ten of the variables used in such classification directly deals with bats (Brasil 2008; Brasil 2022). Such licensing can be improved by using applied bat bioacoustics. In this context, here we describe the use of passive acoustic monitoring to inventory the bat fauna in the Carajás region, in the Brazilian Amazonia. Our objectives were: i) carry out an acoustic inventory to access a portion of subsampled bat species and complement the information available for the region; ii) evaluate the use of a mosaic of different environments in Carajás, including areas under large-scale industrial mining; iii) describe the echolocation calls of the detected species, and iv) make a reference library of these echolocation calls publicly available, aiming to improve the environmental licensing of mineral activity in the region.

## MATERIALS AND METHODS

### Study area

This study was conducted in the Carajás National Forest (FNC – Fig. 1). Created in 1998, FNC covers 411,948.87 hectares, and is in the southeast of the State of Pará (6°4’14.972” S, 50°4’6.886” W), in the Brazilian Amazonia (ICMBio 2016). The FNC is mainly formed by dense rainforests on the plains and in the gentler reliefs of mountainous areas, and open rainforests on steep slopes (ICMBio 2016). On the flat tops of the mountains, there is ferruginous rock vegetation, popularly known as *canga*, a vegetation that varies from herbaceous and shrubby forms to forest (ICMBio 2016). These *canga* areas represent one of the rarest types of rock fields in the Amazon region and are characterized by a high degree of endemism (Viana et al. 2016). Carajás is one of the most important speleological regions in Brazil, with more than 1,500 caves (Piló et al. 2015). This entire region comprises one of the largest mineral provinces on the planet and is the object of industrial exploitation of iron ore, copper, and other minerals by the mining company Vale S.A., the second largest mining company on the planet.

**Figure 1:**
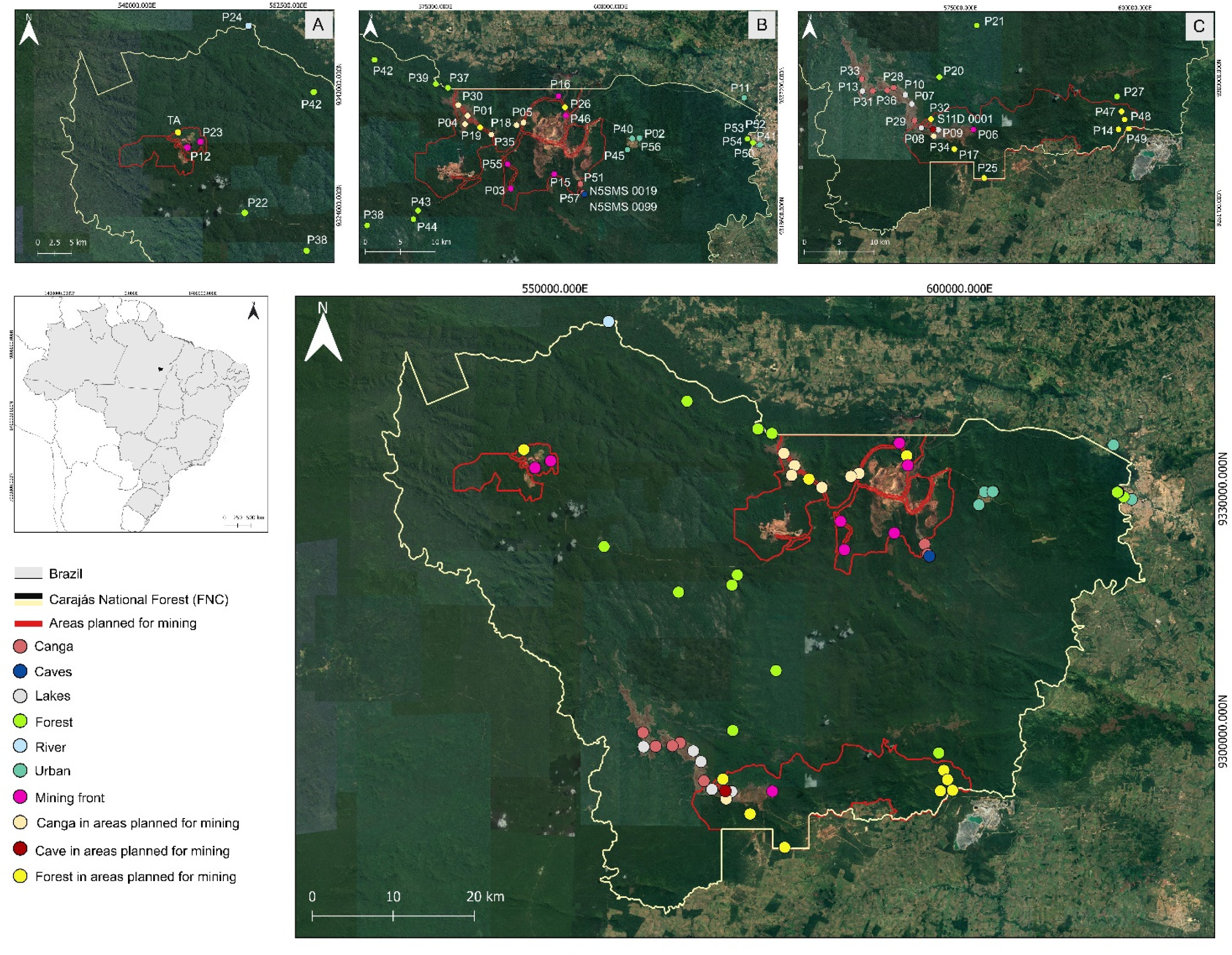
Satellite image of Carajás National Forest, in Pará state, Brazilian Amazonia, with 61 points in different environments sampled for bats between 2021 and 2022, based on the recording of their echolocation calls. Different colors represent different environments sampled, and the red lines mark areas under mining concessions. A) detail of Mina Igarapé Bahia; B) detail of Mina Norte e C) detail of Mina Sul.

We conducted out our study between 2021 and 2023, with three field expeditions, from October 21st to November 17th, 2021, from June 26th to July 22nd, 2022 and May 2nd to 28th, 2023. Sampling at the FNC was carried out in 10 environments: *canga*, caves, lakes, forest, river, urban, mining front (areas where mineral extraction was already underway), *canga* in areas planned for mining, cave in area planned for mining, and forest in areas planned for mining. The areas planned for mining were those where there was already a request for mineral activity that should begin soon. In total, 61 points were sampled (Fig. 1, Supplementary Table S1). For analysis purposes, the lake and river sampling sites were grouped into environments corresponding to their surrounding matrix. For example, if a river was in a forested area, it was placed in the forest category.

### Recordings and analysis of echolocation calls

We used Audiomoths to record bat echolocation signals (www.openacousticdevices.info/audiomoth). To compose the reference bank of echolocation signals and expand the record of species in the region, these recorders were placed at predefined points (see description in the previous section) over a minimum of three consecutive nights, in the three different campaigns to expand the possibility of recording the largest possible number of bat species when active. The recorders were set to record for 15 seconds, followed by a 15-second pause, from 6:00 pm to 6:00 am. The sampling rate of the recorders was 384 kHz, which allowed the recording of frequencies of up to 192 kHz. However, we set 8 kHz as the minimum frequency to reduce the possibility of recording insects and amphibians with signals below this range. The sensitivity of the recorders (gain) was set to average. The recorders were wrapped in a zip-type plastic bag and attached with a wire to the vegetation, approximately 1.5 meters from the ground, or placed on rocks, always with the microphone unobstructed.

All audio files were manually analyzed to detect and identify various echolocation calls emitted by bats in the recordings. Initially, we classified the different calls into sonotypes that corresponds to acoustic morphospecies (Aide et al. 2017). Visual and auditory analyses were performed using Kaleidoscope (Wildlife Acoustics Inc., USA). After analyzing the sonotypes, we sought to identify echolocation signals at the lowest possible taxonomic level (e.g. Jung et al. 2014; Arias-Aguilar et al. 2018), measuring their acoustic parameters: Maximum energy frequency (FME, in kHz); Minimum Frequency (Fmin, in kHz); Maximum Frequency (Fmax, in kHz); Initial frequency (Fin, in kHz); Final frequency (Ffinal, in kHz); Bandwidth (BW, in kHz); Duration (Dur, in seconds); Interval between pulses/signals (IPI, in seconds); Harmonics; and Frequency Modulation (Supplementary Table S2). All acoustic parameters were measured using the Raven Pro 1.6.5 software (www.birds.cornell.edu/ccb/raven-pro/), and the spectrograms were produced as follows:

FFT window = 1024, “Hamming” type, hop size = 102, overlap = 90%. The oscylograms were created using the “seewave” package (Sueur et al. 2008) in the R program, version 4.3.1 (R Core Team 2023). We also used Audacity (https://www.audacityteam.org/download/) to filter background noise from the recordings for which we generated images for the results. Subsequently, to assist in species identification, we compared our data with calls available in recording libraries (https://www.sbeq.net/bioacustica) and specialized literature (Jung et al. 2014; Arias-Aguilar et al. 2018).

### Discriminant function analysis

We used discriminant function analysis to distinguish calls from bat families whose signals are difficult to identify at the species level, ensuring that they were, in fact, of different sonotypes. We used the same acoustic parameters as those described previously for the analysis. For Vespertilionidae, we performed a discriminant analysis for the genera *Eptesicus* and *Lasiurus*, based on 135 sound files and 1193 echolocation calls. We also conducted discriminant function analysis for the genus *Myotis* using 79 audio files and 612 signals. For Molossidae, we performed a discriminant function analysis for *Cynomops* (75 audio files and 398 signals) and *Eumops* (13 audio files and 81 signals). All discriminant function analyses were conducted using the “MASS” package (Venables and Ripley 2002), and corresponding figures were created using the “ggplot2” package (Wickham 2016), both in the R program environment (R Core Team 2023).

### Species richness and environment use

We used interpolation and extrapolation of Hill numbers (Chao et al. 2014), carried out in the “iNEXT” package (Hsieh and Chao 2022) in the R program environment (R Core Team 2023), to verify the sampling effort required to reach the asymptote of species richness for the different environments sampled. We investigated the completeness and variability in species richness estimates using different estimators, such as Chao, Jackknife, and Bootstrap, using the “vegan” package (Oksanen et al. 2022) in the R program environment (R Core Team 2023). It is worth mentioning that for the rarefaction analysis and richness estimators, we chose to exclude the urban environment, as we observed that the rarefaction analysis results related to this environment exhibited a saturation curve suggesting abnormalities.

We employed Generalized Linear Mixed Models (GLMM) to verify whether there were differences in the richness found in the different sampled environments. We generated a model using species richness as a response variable and environment as explanatory variables, while different sampling years and sampling time were entered into the model as random variables; we used the “Poisson” family in this model. In our sampling, there was a difference between the number of sites sampled in forest (15 sites), *canga* (10), forest in an area planned for mining (10), *canga* in an area planned for mining (9), mining front (8), and urban areas (1) when compared to caves (2) and caves in areas planned for mining (one site), which had a relatively smaller number of sampling points. Thus, while carrying out GLMM analyses with cave data, we encountered a convergence problem, and chose to exclude data related to caves and caves in areas planned for mining from the GLMM analysis. For the GLMM analysis, we used the “glmmTMB” package (Brooks et al. 2017), and evaluated the test assumptions using the “DHARMa” package (Hartig 2020). Figures were created using the “ggplot2” package (Wickham 2016), both in the R environment (R Core Team 2023).

## RESULTS

### Richness of sonotypes and species

132 samples were taken at 61 points, which resulted in a total of 142,401.40 minutes of audio. We detected 43 bat sonotypes in Flona Carajás (Table 1), 21 of them classified at the species level and 22 at the genus level. These records were distributed in seven families: Emballonuridae and Molossidae, with five genera each; Phyllostomidae, Mormoopidae, Noctilionidae, and Thyropteridae, with one genus each. For Vespertilionidae, three genera were recorded: *Myotis*, and *Eptesicus* and *Lasiurus* forming a group of species difficult to separate. Families with highest species richness were Emballonuridae and Molossidae, and 11 species were recorded for the first time in the Carajás region: *Cormura brevirostris*, *Diclidurus ingens*, *Peropteryx trinitatis*, *Saccopteryx bilineata*, *Pteronotus alitonus*, *Noctilio albiventris*, *Noctilio leporinus*, *Molossops neglectus*, *Molossus molossus*, *Molossus rufus* and *Promops centralis* (Table 1).

**Table 1.**
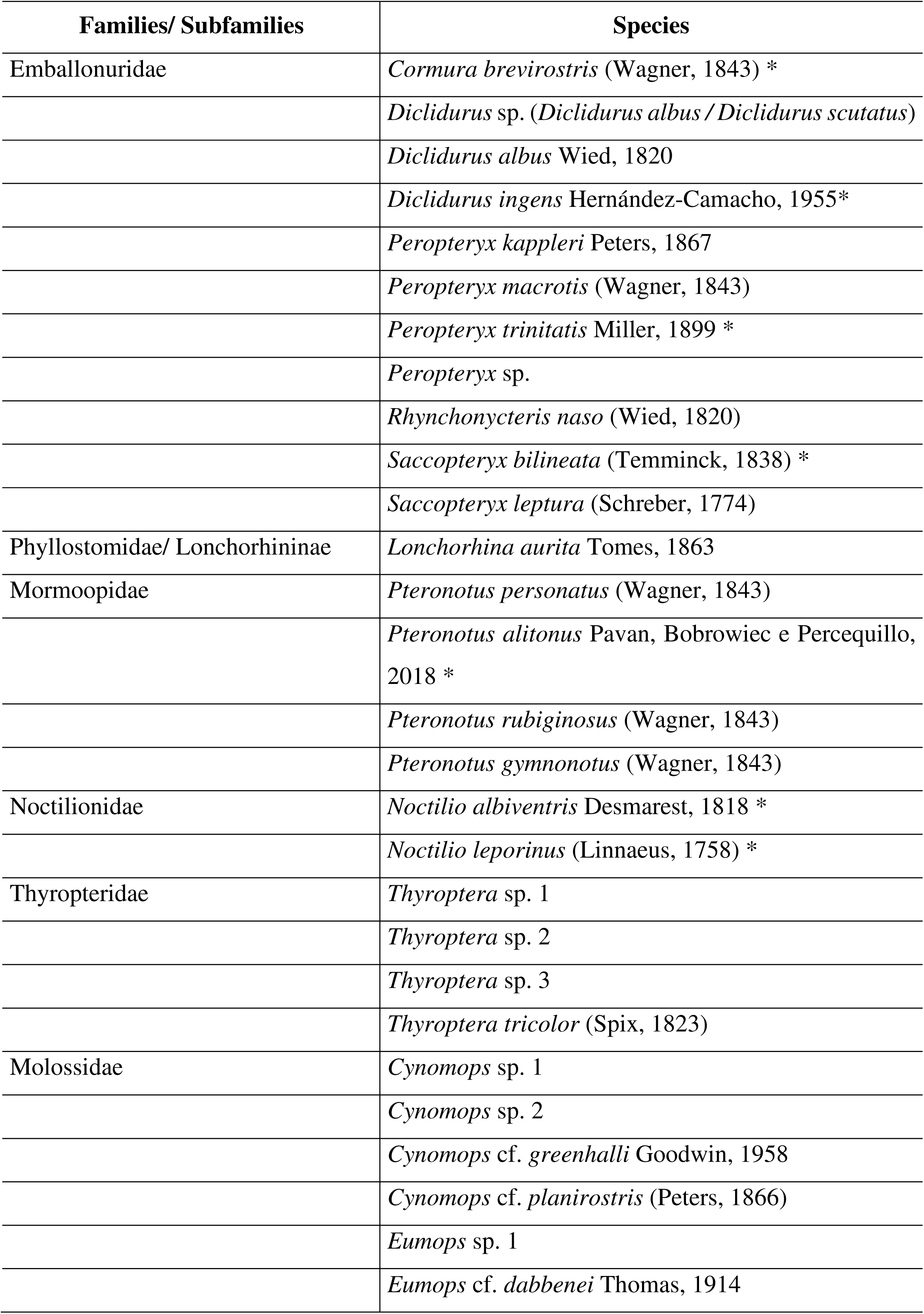
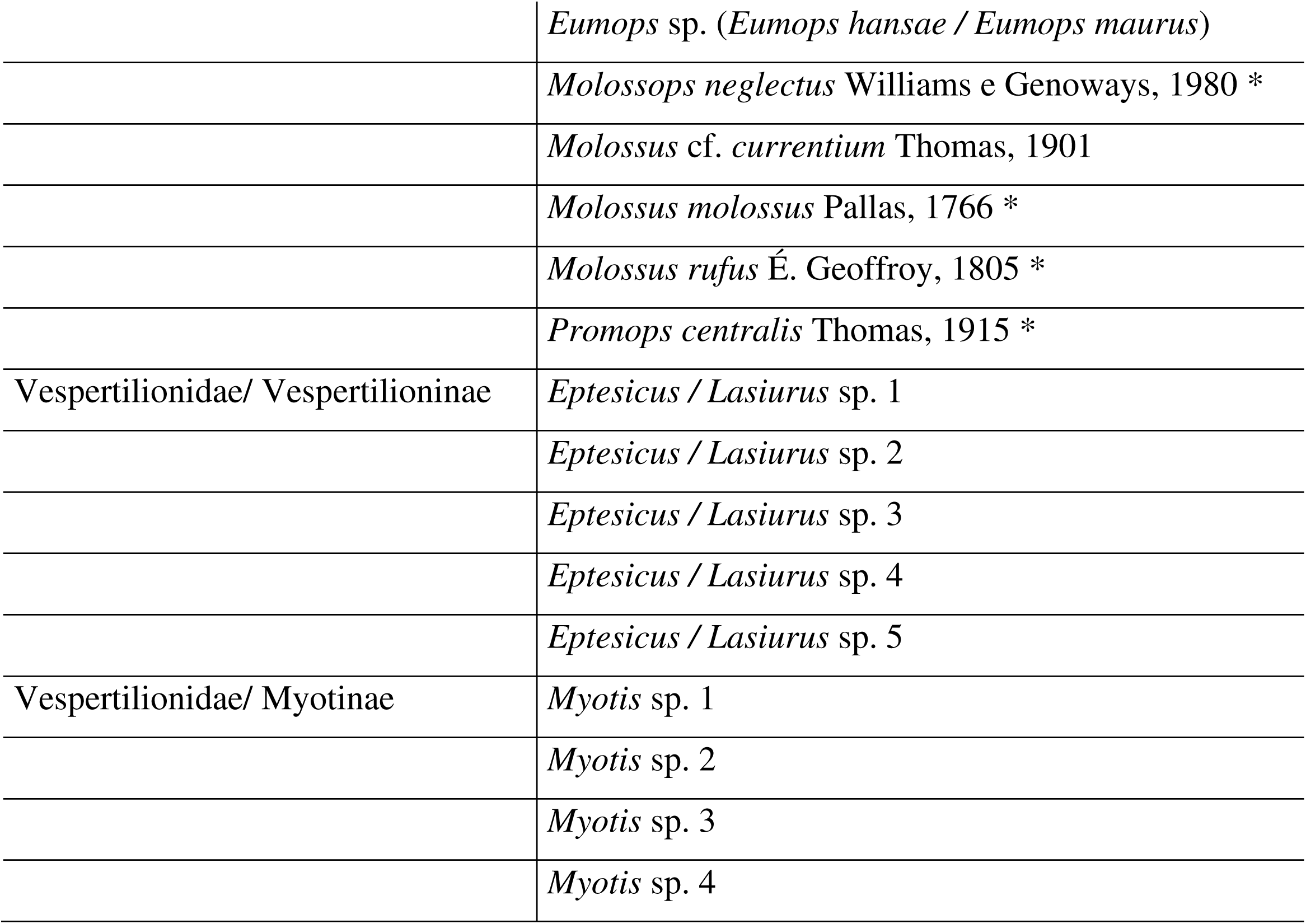
Bat species and sonotypes recorded based on their echolocation calls between 2021 and 2023 at the Floresta Nacional de Carajás, Brazilian Amazonia. *New records for the region of Carajás.

We obtained records of bats at all sampled sites in the three-year inventory. In 2021, we sampled 35 sites, with species richness ranging from 3 to 17 species per site; in 2022, 51 sites were sampled and richness varied from 3 to 12 species per point; in 2023, we sampled 46 sites, with species richness ranging between 2 and 18 per site (Supplementary Tables 1, 2 and 3). When samples taken in different years were pooled together, richness varied from 7 to 35 species (Table 2).

**Table 2.** Observed and estimated bat species richness in different environments in the Carajás National Forest, Brazilian Amazonia, based on the recording of echolocation calls. Chao (chao), Jackknife (jack) and Bootstrap (boot) richness estimators were used, and the values are presented along with their respective standard errors (SE). Recording in minutes.

| Environment | Observed<br>species<br>richness | Richness estimators |  |  | Sampling<br>points | Recordings<br>(min.) |
| --- | --- | --- | --- | --- | --- | --- |
| | | chao ±<br>SE | jack ± SE | boot ±<br>SE | | |
| Canga | 34 | 40 ± 5 | 41 ± 4 | 37 ± 2 | 10 | 22208.50 |
| Forest | 35 | 42 ± 7 | 42 ± 4 | 38 ± 2 | 15 | 36339.75 |
| Mining front | 34 | 46 ± 9 | 45 ± 5 | 39 ± 3 | 8 | 19459.50 |
| Canga in areas<br>planned for<br>mining | 30 | 41 ± 12 | 36 ± 4 | 33 ± 2 | 9 | 25035.25 |
| Forest in areas<br>planned for<br>mining | 34 | 48 ± 10 | 48 ± 8 | 40 ± 4 | 10 | 14217.25 |
| Cave | 7 | 8 ± 8 | 9 ± 2 | 8 ± 1 | 2 | 5590.00 |
| Cave in areas<br>planned for<br>mining | 7 | 7 ± 0 | 7 ± 0 | 7 ± 0 | 1 | 3000.00 |

### Identification of echolocation calls

We described the calls of the 43 recorded sonotypes, with details of the signal structure for each (Appendix S1). Acoustic parameter analysis was performed for 2448 echolocation signals covering different identified genera and families (Supplementary Table S2). Discriminant function analysis was useful to distinguish the distinct sonotypes of some genera. For Molossidae, we analyzed 398 calls from the genus *Cynomops*, indicating four different species. For *Eumops*, we analyzed 81 calls, and sonotypes were separated in three distinct species (Appendix S1). For Vespetilionidae, discriminant function analyses were conducted for the genera *Eptesicus* and *Lasiurus* together, with 1193 calls evaluated, and sonotypes were separated in five distinct species. For *Myotis*, we analyzed 612 calls and sonotypes were separated into four distinct species (Appendix S1).

### Species richness and composition in different environments

The sampling effort in each environment varied from 3,000 minutes for caves in areas planned for mining, to 36,339.75 minutes in the forest environment (Fig. 2). Species richness varied from 4 to 21 species in the sampled locations. The cave environment had the lowest species richness, while *canga* in areas planned for mining and *canga* in mining front the highest (Fig. 3; Table 2). Overall, *canga* and *canga* in areas planned for mining had the highest species richness, followed by river environments in the forest matrix, forest, and forest in areas planned for mining. Furthermore, high species richness was observed at one point in the urban environment and at two mining front points, the latter being a deactivated gold mine in Igarapé Bahia (Fig. 3).

**Figure 2:**
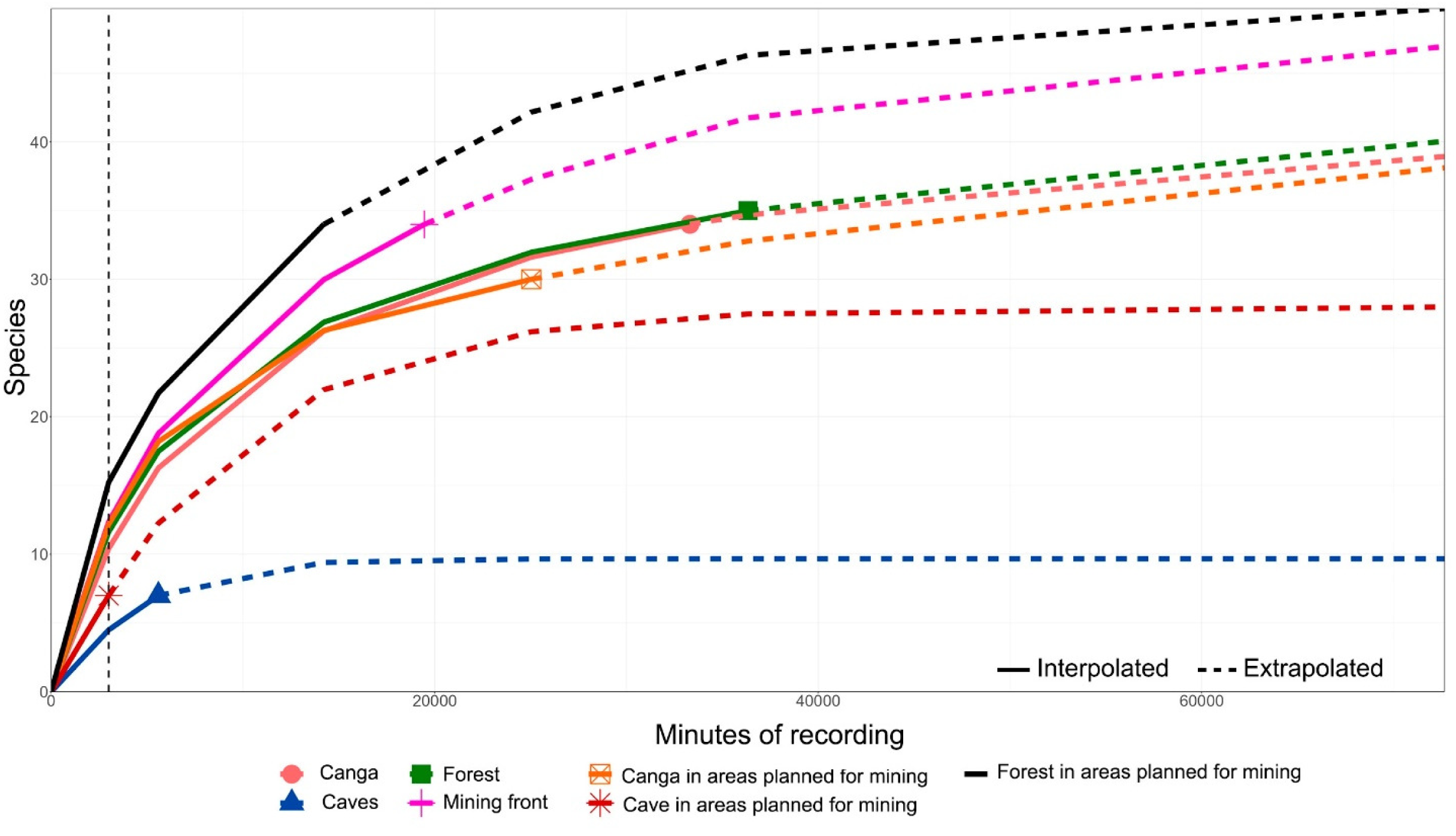
Rarefactions curves for bat species richness recorded in seven different environments at the Carajás National Forest, in Pará state, Brazilian Amazonia, based on the recording of their echolocation calls. We used interpolation and extrapolation of Hill numbers (see Methods) to verify the sampling effort required to reach the asymptote for each environment sampled. Sampling effort expressed in minutes of recording. The dashed vertical line presents the smallest sampling effort used.

**Figure 3:**
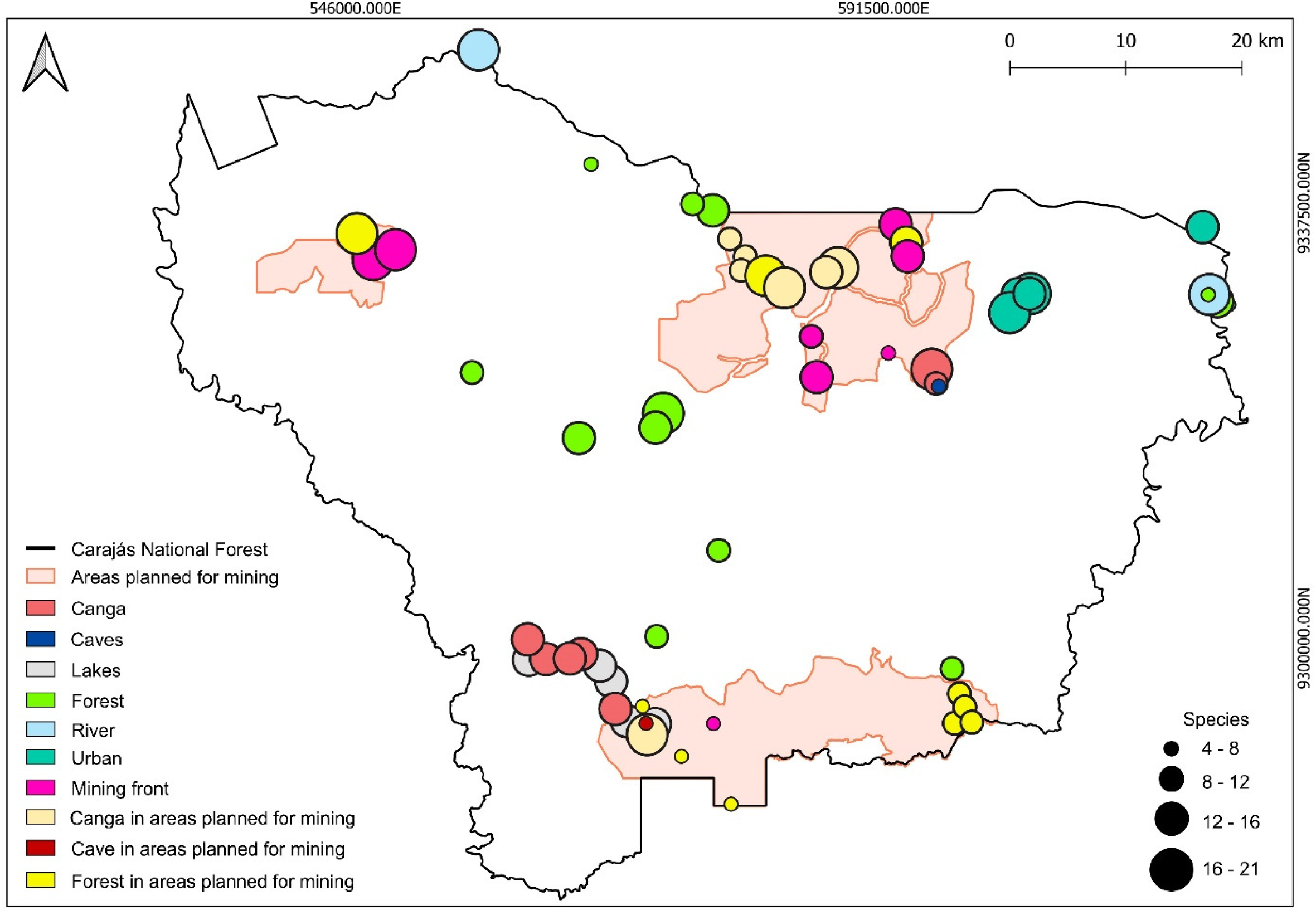
Bat species richness map for Carajás National Forest, in Pará state, Brazilian Amazonia. Richness was obtained based on the recording of echolocation calls at 61 sampling points, between 2021 and 2023. Circles are proportional to the number of species recorded, and colors represent different environment categories sampled.

There was high variation in species richness within the same environment (Fig. 4). In the *canga* environment, we observed the smallest disparity between the minimum (12 spp.) and maximum (18 spp.) richness. On the other hand, the mining front environments (5 - 21 spp.), forest in areas planned for mining (6 - 19 spp.) and urban environment (5 - 19 spp.) showed the greatest variation between the minimum and maximum richness (Fig. 4).

**Figure 4:**
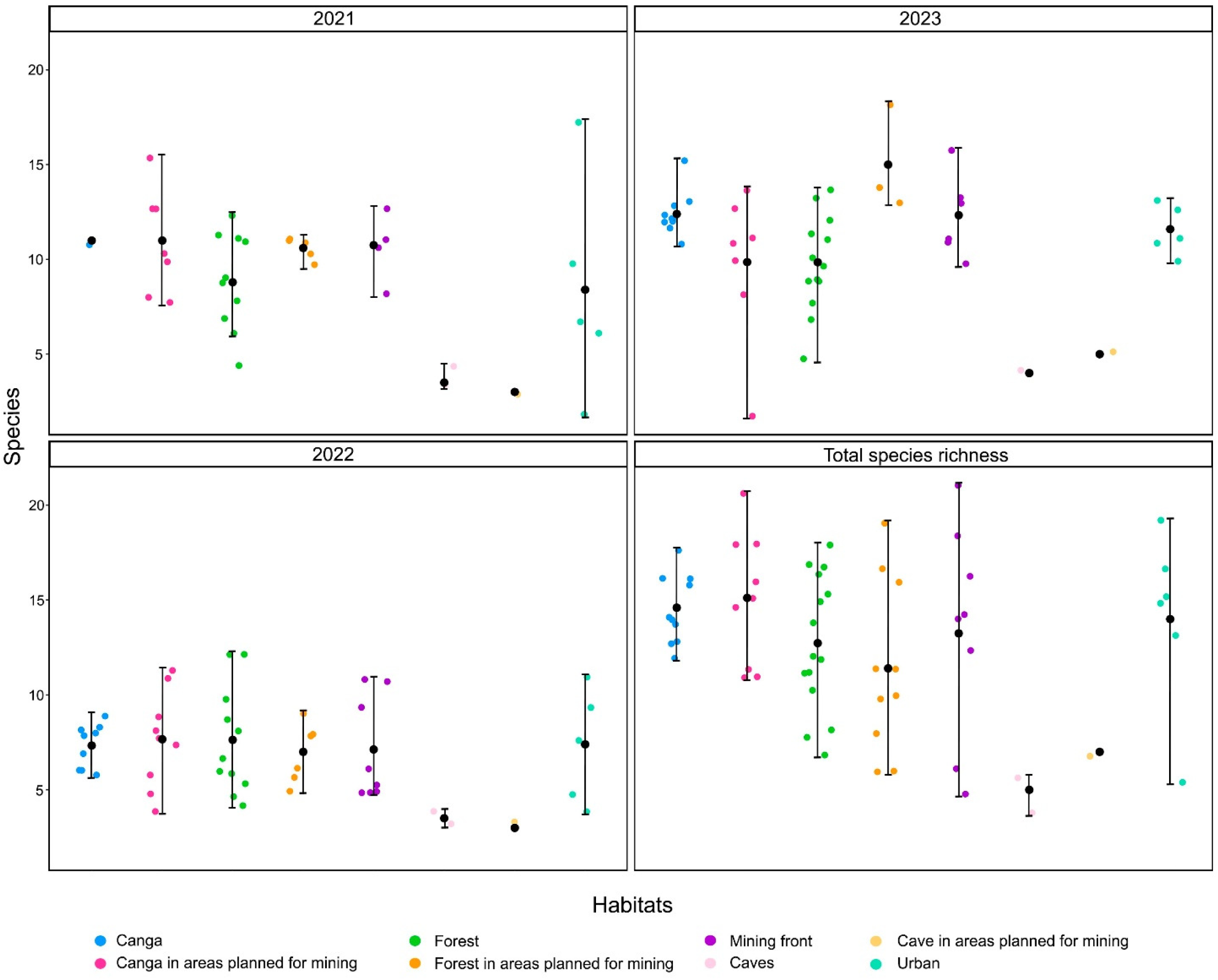
Mean, minimum and maximum values of richness (plus year total) for bat species accessed by bioacoustics in eight categories of environments in the Carajás National Forest, Pará state, Brazilian Amazonia, sampled between 2021 and 2023. Colored dots represent the species richness in each sampled site, and were not aligned for better visualization. The acronyms mean: CA (Canga), CPM (Canga in an area planned for mining), FL (Forest), FPM (Forest in an area planned for mining), FLR (Mining front), CAV (Cave), CAVPM (Cave in an area planned for mining) and UR (Urban).

The species saturation curves were different for each environment but, except for caves in areas planned for mining, all other sampled categories were close to reaching the asymptote. This trend was confirmed when we compared both observed and projected species richness with different estimators: for *canga*, the observed richness was 34 species, with 37 to 40 estimated species; for forest, the observed richness was 35 species and the estimated richness was 38 to 42 species (Table 2).

The GLMM analysis indicated that, species richness was lower in the three years of sampling in forest areas compared to *cangas*, while there were no significant differences between the other environments (Fig. 4, Table 3). Species composition varied depending on sampling location. *Pteronotus personatus*, *Pteronotus rubiginosus*, *Pteronotus gymnonotus*, and *Lonchorhina aurita* were found in all environments sampled. *Pteronotus personatus* and *P. rubiginosus* were found in 60 and 59 of the 61 sites sampled, respectively. In contrast, *Promops centralis*, *Thyroptera* sp. 3, *Eumops* sp. 1, *Diclidurus albus*/*Diclidurus scutatus* only occurred at a single sampling point, whereas *Noctilio albiventris*, *Molossops neglectus*, *Peropteryx trinitatis*, *Rhynchonycteris naso* and *Pteronotus alitonus* occurred at two to five sampling points. *Diclidurus albus*, *Peropteryx macrotis*, *Peropteryx trinitatis*, and *Peropteryx* sp. occurred mainly in *canga* and *canga* in areas planned for mining (Fig. 5).

**Figure 5:**
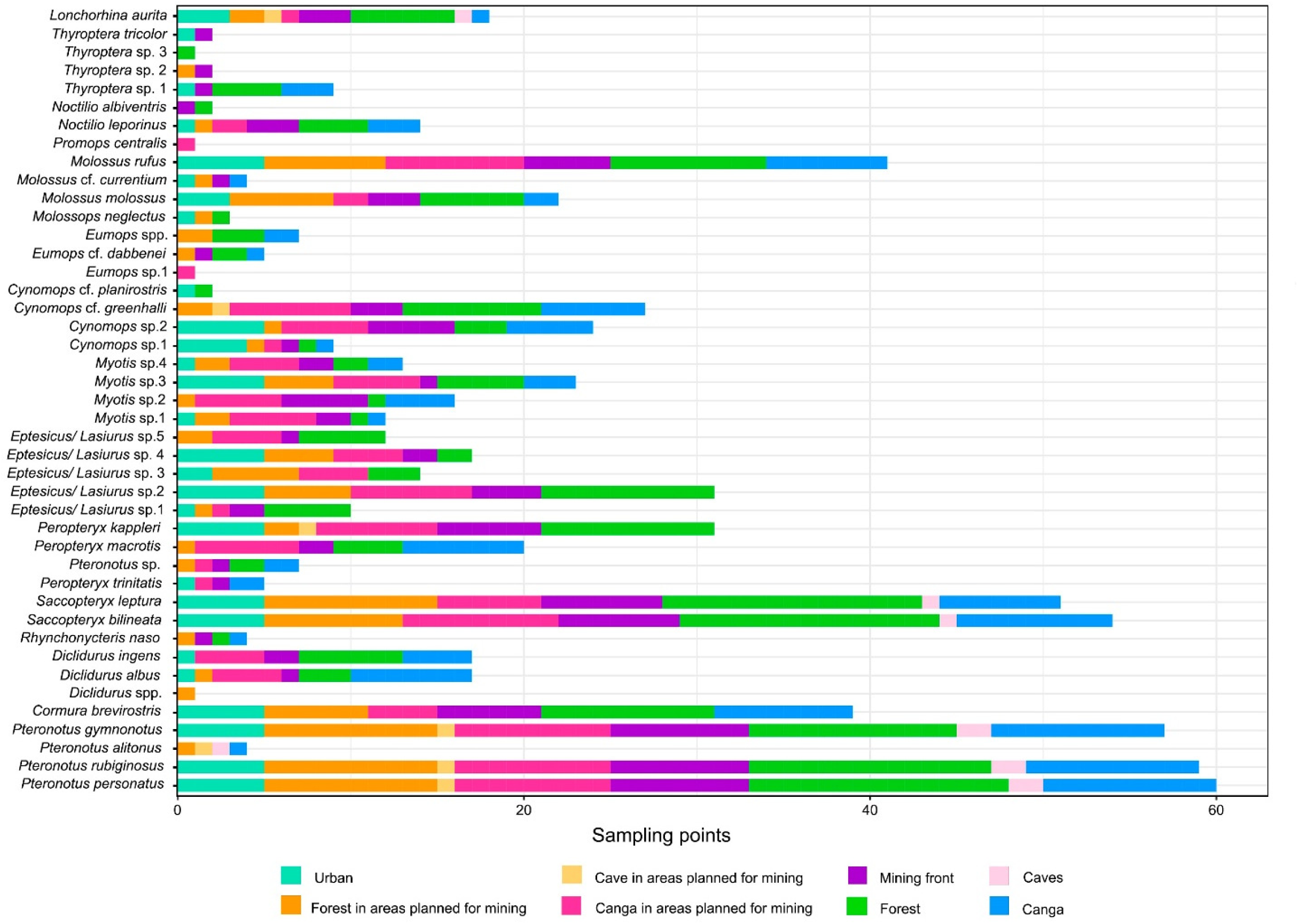
Bat species recorded by bioacoustics at 61 sampling points in different environments at Carajás National Forest, Pará state, Brazilian Amazonia, between 2021 and 2023.

**Table 3.** Values obtained using a mixed generalized linear model (GLMM) comparing the richness of bat species accessed by bioacoustics in different environments in the Carajás National Forest, Brazilian Amazonia, between 2021 and 2023. There was statistically significant difference only between forest and *canga* environments.

|  | Estimate | Standard error | z- value | p- value |
| --- | --- | --- | --- | --- |
| (intercept) | 2.33572 | 0.12510 | 18.671 | <2e-16 |
| Forest<sup>1</sup> | -0.21001 | 0.09381 | -2.239 | **0.0252** |
| Mining front<sup>1</sup> | -0.05132 | 0.10377 | -0.495 | 0.5209 |
| Canga in area planned for mining<sup>1</sup> | -0.11102 | 0.10032 | -1.107 | 0.2684 |
| Forest in area planned for mining<sup>1</sup> | -0.05231 | 0.11354 | -0.461 | 0.6450 |
| Urban<sup>1</sup> | -0.05313 | 0.11307 | -0.470 | 0.6384 |

<sup>1</sup> Different to *canga*

## DISCUSSION

In this study, carried out in the Carajás National Forest region in the State of Pará, Brazilian Amazonia, we used bioacoustics as a tool to inventory bat species in the region, as well as to infer species richness and the use of a mosaic of different environments, including forests, areas of metallophilic savannas (*canga*), and areas under the influence of industrial iron ore mining. After approximately 142,000 minutes of recording at 61 sampling points, we identified at least 43 sonotypes of species belonging to seven of the nine bat families occurring in Brazil. Our results demonstrated that the use of bioacoustics resulted in an increase in the known bat species richness in the Carajás region. Species richness varied between caves and other environments, with forests being richer. However, the differences in species richness between non-cave environments were subtle, whereas *canga* environments demonstrated greater stability in species richness, and a more distinct species composition.

All 43 recorded sonotypes could be identified at the genus level, 21 of them at the specific level, resulting in 11 species being recorded for the first time in the Carajás region (Tavares et al. 2012). This confirms that the use of bioacoustics can increase the number of bat species recorded in areas of high biodiversity, such as the Brazilian Amazonia, helping access species that had previously not been detected by conventional inventories. The addition of new species produces a more accurate characterization of local and regional biodiversity, which is particularly important in large, species-rich, but poorly sampled areas such as the Brazilian Amazonia (e.g., Aguiar et al. 2020; Carvalho et al. 2023).

Our bat acoustic inventory in the Carajás region had similarities with acoustic inventories from other areas in Amazonia. Studies conducted by Appel et al. (2021) at the Ducke Reserve, near Manaus, in Central Amazonia, acoustically identified 17 species and five non-identified sonotypes; while Di Ponzio et al. (2023), sampling on islands in the lake of the Balbina Hydroelectric Dam, also in Central Amazonia, identified 22 sonotypes, 17 of which were identified at the specific level, four at the genus level, and one as a phonic complex formed by two acoustically indistinguishable species (*Eptesicus brasiliensis*/*E. chiriquinus*). In the first study (Appel et al. 2021), sampling covered 101 nights, totaling 72,720 minutes of recording, which resulted in approximately 0.018 species/hour of recording, and in the second (Di Ponzio et al. 2023) there were 141 nights and 109,980 minutes of recording, resulting in 0.012 species/hour of recording. Although we recorded more species in Carajás than in these two inventories, we employed a greater sampling effort (132 nights, 142,401 minutes of recording), which, when standardized, resulted in values similar to that of Appel et al. (2021). Except for caves – a non-focal environment in our study – all our species saturation curves were close to reaching the asymptote. The few bats acoustic inventories available for the Brazilian Amazonia points now that 22 up to 43 sonotypes, and 0.012 to 0.018 species/hour of recording, seems a reasonable baseline output for future comparisons.

### New records for Carajás

The 11 species previously unrecorded for the Carajás region (*Cormura brevirostris, Diclidurus ingens, Peropteryx trinitatis, Saccopteryx bilineata, Pteronotus alitonus, Noctilio albiventris, Noctilio leporinus, Molossops neglectus, Molossus molossus, Molossus rufus and Promops centralis*) belong to four families (Emballonuridae, Mormoopidae, Noctilionidae, Molossidae). Some of these species were expected for Carajás. *Noctilio*, for example, is a genus widely distributed throughout the Amazon and Brazil (https://salve.icmbio.gov.br/); however, there were no formal records for the Noctilionidae family in the Carajás region. These two species (*N. leporinus* and *N. albiventris*) are normally associated with areas close to water bodies (Schnitzler et al., 1994). In fact, we detected that *N. albiventris*’ distribuition was limited in our samplings, and was found at only two locations. In contrast, *N. leporinus* had a wider distribution, and was recorded at 14 sampling sites. The same is true for *C. brevirostris* and *S. bilineata*, species with a wide distribution in the Neotropical region but with no previous formal records for Carajás. These species are considered occasional cave users, that is, species that have been recorded using caves but prefer other types of roosts (Barros and Bernard 2023b). Although dozens of bat inventories have been conducted in Carajás by consultancies hired by Vale S.A. (Tavares et al. 2012), our records exemplify how the bat fauna previously known for the region was biased towards a specific type of environment of greatest interest for environmental licensing purposes, notably caves and their immediate surrounding areas. Our records also emphasize that an expanded sample covering different existing environments proved very important to assess the actual species richness in Carajás, and environment-specific inventories, such as those adopted by environmental licensing in mining activities, which frequently focus on the type of environment that will be directly suppressed and/or on habitats with legal protection, may miss some components of the local biodiversity (Sonter et al. 2018).

The largest number of new occurrences in Carajás were in Molossidae, a family with several species with a wide distribution in the Neotropical region (do Amaral et al. 2023). However, molossids are considered difficult to capture in mist nets, as they fly above the forest canopy or in open landscapes (Kalko et al. 2008). Some species recorded by us have a wide occurrence in the Amazon region and Brazil, such as *Molossus rufus* and *Molossus molossus* (do Amaral et al. 2023); however, with no previous no records for Carajás. This also reinforces the efficiency of using bioacoustics to record species with a profile of using environments inaccessible to mist nets, causing them to be frequently under-sampled in inventories with the sole use of this approach. In fact, in Carajás, *M. rufus* and *M. molossus* were identified at several sampling points, suggesting a wide regional distribution. Other species from the same family, such as *Promops centralis* and *Molossus neglectus*, have few records in the Amazon region (do Amaral et al. 2023). The records in Carajás contribute to filling gaps in the occurrence of these species but confirm their rarity, as they were limited to just one to three recording points.

### Bats and environments

Species richness differed mainly between caves and other environments, such as forest, *canga*, areas planned for mining in forest, areas planned for mining in *canga*, mining fronts and urban locations. In non-cave environments, the richness varied between 30 and 35 species. In caves and caves in areas planned for mining, species richness was lower, with seven species recorded. In the Neotropics, some cave environments may have high bat richness, sometimes with more than 20 species recorded in a single cave (Barros et al. 2021). Although phyllostomid species often dominates these inventories, species in this family are frequently poorly detectable based solely on acoustic recordings (e.g. O’Farrell and Gannon 1999; Arias-Aguilar et al. 2018), as in our approach.

Previous studies (Barros et al. 2020; Barros and Bernard 2023a) identified a positive relationship between cave size and species richness. However, species richness also differs between lithologies: while carbonate caves tend to be larger and have greater richness, caves in iron formations, as in Carajás, are often smaller and have a lower bat species richness (Barros and Bernard 2023b). This recent review, based on data for 416 caves in Brazil, pointed out that species richness was 5.3 ± 4.5 (mean ± standard deviation) in carbonate caves, 6.3 ± 3.0 in magmatic caves, 3.4 ± 3.2 in siliciclastic caves, and 1.7 ± 1.8 in iron caves. Therefore, caves in Carajás are expected to have a lower species richness than caves in other lithologies. However, even with the lower richness observed, our data for Carajás clearly demonstrate how effective bioacoustics can expand access to species richness even in cave environments.

When analyzing the number of species per sampling location, the points with the highest species richness were in *canga*, *canga* in areas planned for mining, forest and forest in areas planned for mining, with 16 to 21 species in these areas. Although not under current mining activity, some of these areas already have authorization for mineral exploration. Considering that our data on species richness in these environments were obtained in pre-mining conditions, it is possible to monitor the impacts of the proximity of mining fronts and the installation of mining infrastructure on bats there (e.g. Cristescu et al. 2012; Bunkley et al. 2015;). By adopting sampling protocols similar to those we used, the monitoring of bats in Carajás could be a case study on the use of applied bioacoustics for environmental licensing processes involving mining in biodiversity-rich areas, which is useful not only for Amazonia but also for other tropical environments with high bat species richness.

## ACKNOWLEDGEMENTS

This study was possible thanks to the financial support and logistics provided by the Núcleo Getsor Integrado de Carajás, Instituto Chico Mendes de Conservação da Biodiversidade. We thank André Luis Macedo Vieira, Paulo Jardel Braz Faiad, Wendelo Silva Costa and the entire NGI Carajás team (Silvana Araújo, Amanda Figueiredo, José Sebastião ’Pezão’ and Neuzivaldo ’Theo’), who spared no effort to provide all necessary conditions for access to recording areas. We also thank the Vale S.A. Speleology Sector in Carajás, especially Bruno Scherer. The fieldwork would not have been possible without the invaluable help of Narjara Tércia Pimentel, Eder Barbier and Frederico Hintze. L. Gomes received a research grant from ICMBio, as part of the Project ’Bioacoustics of bats in Carajás: Species inventory and assessment of environmental impacts of mineral exploration’. E. Bernard has a CNPq fellowship.

