## Supplementary Material for "Listening to the bats of Carajás: Applied bioacoustics for species inventory and environment use in a mosaic of forests, savannas, and industrial mining in the Brazilian Amazonia"

1 – Laboratório de Ciência Aplicada à Conservação da Biodiversidade, Departamento de Zoologia, Centro de Biociências, Universidade Federal de Pernambuco, Brazil.

2- Instituto de Desenvolvimento Sustentável Mamirauá (IDSM), Projeto Providence, Tefé, Amazonas, Brazil.

L.G.: https://orcid.org/0000-0001-9526-2158 E.B.: https://orcid.org/0000-0002-2304-1978

**Table S1.** Points acoustically sampled for bat echolocation calls in the Floresta Naciona de Carajás, Brazilian Amazonia, between 2021 a 2023. Habitat: FLM (mining front), CPM (canga in areas planned for mining), CAVPM (caves in areas planned for mining), FPM (forest in areas planned for mining). Samplings are expressed in minutes of recording.

| Points | Habitat | Longitude | Latitude | Year | | | Total sampling (min) |
|---|---|---|---|---|---|---|---|
| | | | | 2021 (min) | 2022 (min) | 2023 (min) | |
| P01 | CPM | -50.2807 | -6.03494 | 1125 | 1109.5 | 1125 | 3359.5 |
| P02 | Urban | -50.0587 | -6.06353 | 830 | 1125 | 1125 | 3080 |
| P03 | FLM | -50.2249 | -6.1289 | 0 | 1125 | 1125 | 2250 |
| P04 | CPM | -50.2839 | -6.04573 | 0 | 1100 | 1125 | 2225 |
| P05 | CPM | -50.2087 | -6.04359 | 1125 | 1125 | 1125 | 3375 |
| P06 | FLM | -50.305 | -6.39901 | 0 | 1125 | 0 | 1125 |
| P07 | Lake/ Canga | -50.3848 | -6.36604 | 0 | 1125 | 1125 | 2250 |
| P08 | Lake/ CPM | -50.3726 | -6.39701 | 0 | 1125 | 1125 | 2250 |
| P09 | Lake/ CPM | -50.3505 | -6.39937 | 750 | 1125 | 0 | 1875 |
| P10 | Lake/ Canga | -50.3933 | -6.35404 | 0 | 1125 | 1125 | 2250 |
| P11 | Urban | -49.9246 | -6.01123 | 0 | 0 | 1125 | 1125 |
| P12 | FLM | -50.5705 | -6.03768 | 937.5 | 1125 | 1125 | 3187.5 |
| P13 | Lake/ Canga | -50.4489 | -6.3494 | 0 | 1125 | 1125 | 2250 |
| P14 | FPM | -50.1172 | -6.39842 | 750 | 0 | 0 | 750 |
| P15 | FLM | -50.1691 | -6.11008 | 0 | 1125 | 0 | 1125 |
| P16 | FLM | -50.1636 | -6.00952 | 884.5 | 1125 | 1125 | 3134.5 |

| | | | | | | | |
|---|---|---|---|---|---|---|---|
| P17 | FPM | -50.3298 | -6.42444 | 0 | 1125 | 0 | 1125 |
| P18 | CPM | -50.2177 | -6.04697 | 1125 | 1125 | 1125 | 3375 |
| P19 | FPM | -50.2647 | -6.04989 | 1125 | 1092.3 | 1125 | 3342.3 |
| P20 | Forest | -50.3493 | -6.33112 | 750 | 1125 | 1125 | 3000,00 |
| P21 | Forest | -50.3011 | -6.26404 | 750 | 1125 | 1228.3 | 3103.3 |
| P22 | Forest | -50.4933 | -6.12562 | 937 | 1125 | 1125 | 3187.5 |
| P23 | FLM | -50.553 | -6.02993 | 1125 | 1137.5 | 1125 | 3387.5 |
| P24 | River/ Forest | -50.4885 | -5.87406 | 0 | 1365.5 | 1125 | 2490.5 |
| P25 | FPM | -50.2912 | -6.4617 | 0 | 1125 | 0 | 1125 |
| P26 | FPM | -50.1554 | -6.02376 | 0 | 1125 | 1125 | 2250 |
| P27 | Forest | -50.1191 | -6.3559 | 750 | 0 | 0 | 750 |
| P28 | Canga | -50.4082 | -6.34487 | 0 | 1125 | 1125 | 2250 |
| P29 | Canga | -50.3814 | -6.38752 | 0 | 1125 | 1125 | 2250 |
| P30 | COM | -50.2927 | -6.02117 | 1125 | 1114.3 | 0 | 2239.3 |
| P31 | Canga | -50.4353 | -6.34891 | 0 | 1125 | 1125 | 2250 |
| P32 | FPM | -50.3602 | -6.38555 | 0 | 1125 | 0 | 1125 |
| P33 | Canga | -50.4497 | -6.33352 | 0 | 1125 | 1125 | 2250 |
| P34 | CPM | -50.3566 | -6.40759 | 750 | 1125 | 1125 | 3000 |
| P35 | CPM | -50.2501 | -6.05921 | 1125 | 1086.5 | 1125 | 3336.5 |
| P36 | Canga | -50.4168 | -6.34834 | 0 | 633.5 | 1125 | 1758.5 |
| P37 | Forest | -50.306 | -5.99901 | 1125 | 1120 | 1125 | 3370 |
| P38 | Forest | -50.4101 | -6.17658 | 937.5 | 1125 | 1125 | 3187.5 |
| P39 | Forest | -50.3217 | -5.99395 | 0 | 1125 | 1125 | 2250 |
| P40 | Urban | -50.0685 | -6.06393 | 750.5 | 1125 | 1125 | 3000.5 |
| P41 | Urban | -49.9042 | -6.07188 | 1678.5 | 1125 | 0 | 2803.5 |
| P42 | Forest | -50.4008 | -5.96299 | 0 | 1125 | 0 | 1125 |
| P43 | Forest | -50.3444 | -6.15724 | 937.5 | 0 | 1125 | 2062.5 |
| P44 | Forest | -50.3504 | -6.16847 | 937.5 | 1125 | 1125 | 3187.5 |
| P45 | Urban | -50.0748 | -6.07841 | 750 | 1125 | 1120.5 | 2995.5 |
| P46 | FLM | -50.1543 | -6.03444 | 750 | 1125 | 1125 | 3000 |
| P47 | FPM | -50.1134 | -6.37533 | 750 | 0 | 0 | 750 |
| P48 | FPM | -50.1093 | -6.38571 | 750 | 0 | 0 | 750 |
| P49 | FPM | -50.1037 | -6.39764 | 750 | 0 | 0 | 750 |
| P50 | Forest | -49.913 | -6.06918 | 1125 | 1125 | 1125 | 3375 |
| P51 | Canga | -50.1353 | -6.1226 | 1125 | 1125 | 1229.5 | 3479.5 |
| P52 | River/ Forest | -49.9193 | -6.06388 | 750 | 1125 | 1125 | 3000 |
| P53 | Forest | -49.9199 | -6.06397 | 0 | 0 | 1125 | 1125 |
| P54 | Forest | -49.9202 | -6.06415 | 0 | 0 | 1125 | 1125 |
| P55 | FLM | -50.2291 | -6.0972 | 0 | 1125 | 1125 | 2250 |
| P56 | Urban | -50.0594 | -6.06362 | 1297.5 | 1125 | 1125 | 3547.5 |
| P57 | Canga | -50.132 | -6.1337 | 0 | 0 | 1220.5 | 1220.5 |
| S11D_0001 | CAVPM | -50.3574 | -6.39884 | 750 | 1125 | 1125 | 3000 |
| TA | FPM | -50.583 | -6.01732 | 0 | 1125 | 1125 | 2250 |
| N5SMS_0019 | Cave | -50.1299 | -6.13617 | 1125 | 1074.5 | 1166 | 3365.5 |
| N5SMS_0099 | Cave | -50.13 | -6.1359 | 1125 | 1099.5 | 0 | 2224.5 |
| | Total: | | | 33378.5 | 56933.1 | 52090.2 | 142401.4 |

#### Appendix

#### Description of echolocation calls

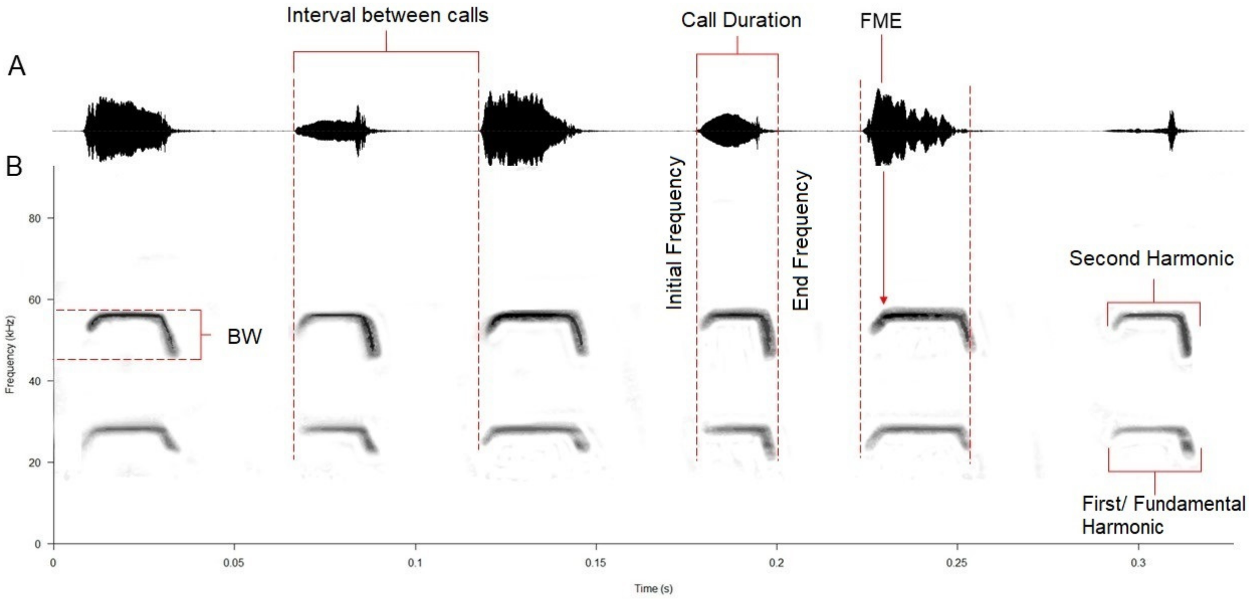


**Figure S1**. Acoustic parameters extracted from bat echolocation calls for possible species identification.

**Tabela S2.** Acoustic parameters extracted from bat echolocation calls used for sonotype or species identification. Raven software was used measurements. Frequency modulation was visually classified based on sound structure. Frequencies were converted from hertz (Hz) to kilohertz (kHz), and duration is expressed in seconds (s).

| **Acoustic Parameters** | **Description** | **Measurements on Raven** |
|---|---|---|
| Frequency of Maximum Energy **- FME** | The frequency at which the sound signal has its maximum peak intensity | Peak Freq (Hz) |
| Minimum Frequency - **FMIN** | Lowest frequency of the sound signal | Freq 5% (Hz) |
| Maximum Frequency - **FMAX** | Highest frequency of the sound signal | Freq 95% (Hz) |
| Initial Frequency/Start frequency - **FINITIAL** | Frequency that initiates the sound signal | Low Freq (Hz) |
| End Frequency - **FEND** | Frequency that ends the sound signal | High Freq (Hz) |
| Bandwidth - **BW** | Total frequency range occupied by the sound signal | Bandwidth 90% (Hz) |
| Call duration - **Dur** | The duration of a single sound signal, measured from the start to the end of the signal | Delta Time (s) |
| Inter-pulse Interval - **IPI** | The interval between two consecutive sound signals, measured from the start of one pulse or call to the start of the next pulse or call | Delta Time (s) |
| Harmonic | Some sound signals have their energy distributed across various distinct and evenly spaced frequencies, known as harmonics. These frequencies are integer multiples of the lowest frequency (first/ fundamental harmonic) | Peak Freq (Hz) de cada harmônico |
| Modulated Frequency - **FM** | Changes in the frequency of the sound signal over time, can be ascending or descending | |

**Appendix S1** – Description of bat echolocation calls recorded in the Floresta Nacional de Carajás, southeastern Pará state, in Brazilian Amazonia, between 2021 and 2023. Each species/sonotype has a textual description of the recorded calls, a table with the respective acoustic parameters (average ± standard deviation, and minimum and maximum values in parenthesis), and a sonogram with oscilogram. Parameter acronyms as in Table S2. Calls are presented in families, genus, species or sonotypes. Frequencies in kHz, and duration in seconds.

#### Family Emballonuridae

Echolocation calls for 11 sonotypes, five genus: *Cormura brevirostris* (Cbre), *Diclidurus albus*/*Diclidurus scutatus* (Dalb/scu), *Diclidurus albus* (Dalb), *Diclidurus ingens* (Ding), *Peropteryx kappleri* (Pkap), *Peropteryx macrotis* (Pmac), *Peropteryx trinitatis* (Ptri), *Peropteryx* sp. (Psp), *Rhynchonycteris naso* (Rnas), *Saccopteryx bilineata* (Sbil), and *Saccopteryx leptura* (Slep).

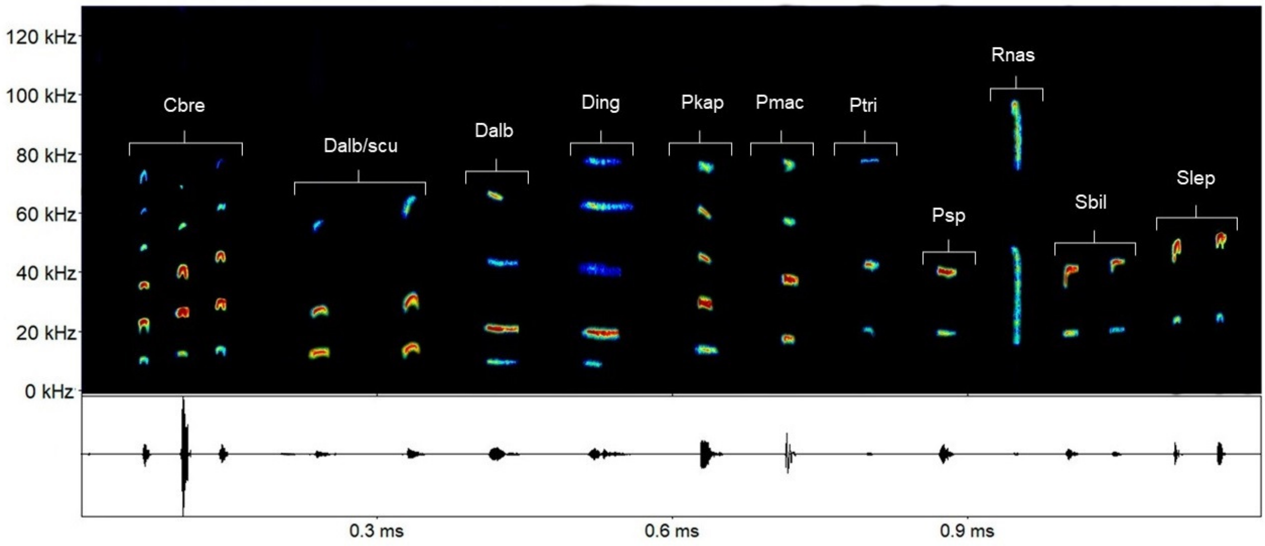

***Cormura brevirostris*** (Wagner, 1843): Composed by three quasi-constant (qCF) calls (types a, b, and c) with gradual frequency variation. FME for call a is around 24.1 kHz, for call b is 27.2 kHz, and 30.8 kHz for call c.

| Parameter | Call a | Call b | Call c |
|---|---|---|---|
| Finitial | 20.25 ±1.08 (18.91-21.51) | 23.97 ±0.36 (23.49-24.33) | 27.39 ±0.53 (26.71-28.14) |
| Ffinal | 25.82 ±0.50 (25.24-26.52) | 29.12 ±0.37 (28.66-29.52) | 32.48 ±0.39 (32.07-32.93) |
| Fmin | 23.25 ±0.19 (22.93-23.43) | 25.89 ±0.55 (25.37-26.73) | 29.72 ±0.42 (29.25-30.25) |
| Fmax | 24.77 ±0.25 (24.46-25.15) | 28.12 ±0.18 (27.93-28.40) | 31.53 ±0.24 (31.21-31.87) |
| FME | 24.13 ±0.10 (24.00-24.25) | 27.28 ±0.23 (27.00-27.50) | 30.86 ±0.38 (30.18-31.18) |
| BW | 1.62 ±0.16 (1.41-1.81) | 2.16 ±0.50 (1.44-2.62) | 1.79 ±0.48 (1.18-2.37) |
| Dur | 0.017 ±0.003 (0.01-0.02) | 0.017 ±0.005 (0.01-0.02) | 0.016 ±0.006 (0.01-0.02) |
| IPI | 0.065 ±0.011 (0.04-0.07) | - | - |
| IC | 0.072 ±0.005 (0.06-0.07) | - | - |

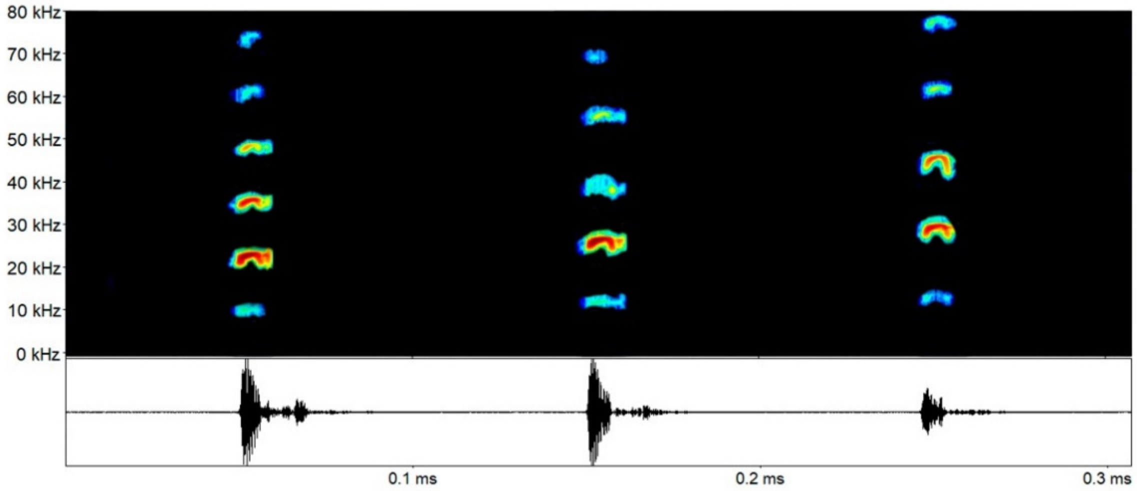

***Diclidurus* spp.**: Composed by two alternating calls (a, lower; b, higher frequency) with quasi-constant (qCF) shape, and FME around 28.0 kHz for call a, and 31.4 kHz for call b.

| Parameter | Call a | Call b |
|---|---|---|
| Finitial | 25.52 ±0.54 (24.54-26.46) | 27.69 ±0.53 (26.70-28.65) |
| Ffinal | 29.80 ±0.48 (28.95-28.12) | 34.45 ±0.74 (33.10-35.61) |
| Fmin | 27.44 ±0.40 (26.62-28.12) | 30.20 ±0.39 (29.62-30.75) |
| Fmax | 28.91 ±0.45 (28.12-29.62) | 33.00 ±0.51 (31.87-33.75) |
| FME | 28.03 ±0.40 (27.00-28.50) | 31.41 ±0.65 (30.75-32.62) |
| BW | 1.46 ±0.28 (1.12-2.25) | 2.79 ±0.26 (2.25-3.00) |
| Dur | 0.015 ±0.001 (0.01-0.01) | 0.015 ±0.001 (0.01-0.01) |
| IPI | 0.096 ±0.003 (0.08-0.10) | - |
| IC | 0.262 ±0.090 (0.20-0.57) | - |

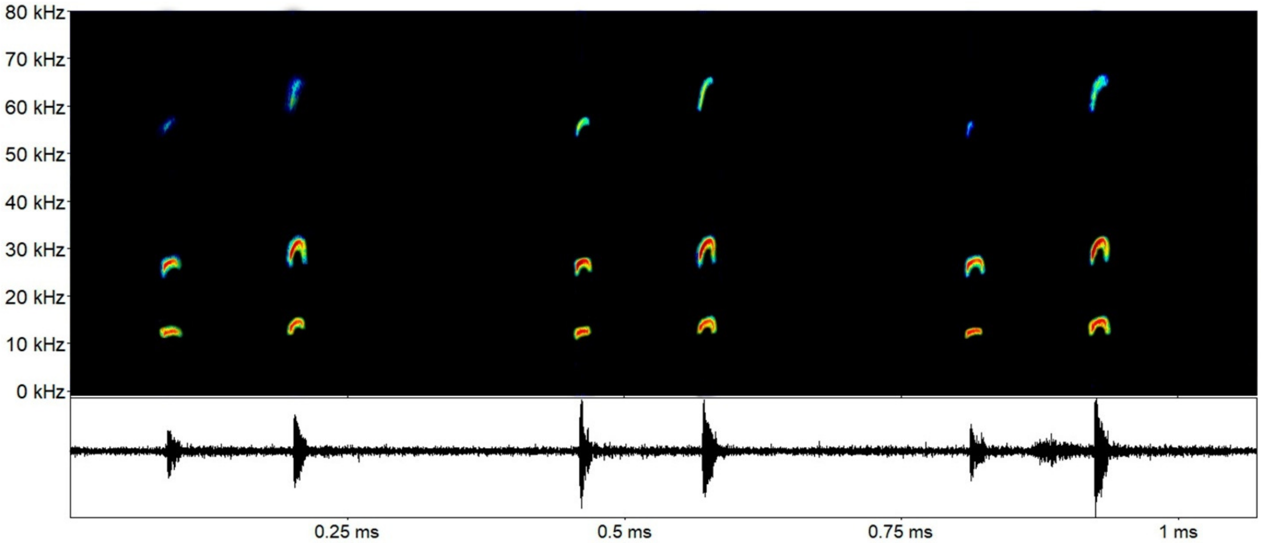

***Diclidurus albus*** Wied, 1820: Composed by a single call, although reported in the literature that the species may emit sequencies with two calls (Arias-Aguilar et al., 2018). The calls we identified had a light frequency modulation, characterized as a quasi-constant (qCF) shape, with FME around 22.3 kHz.

| Parameter | Call |
|---|---|
| Finitial | 20.63 ±0.11 (20.51-20.74) |
| Ffinal | 24.13 ±0.18 (24.01-24.35) |
| Fmin | 21.93 ±0.12 (21.80-22.05) |
| Fmax | 22.86 ±0.22 (22.66-23.10) |
| FME | 22.39 ±0.18 (22.17-22.50) |
| BW | 0.92 ±0.10 (0.85-1.05) |
| Dur | 0.016 ±0.001 (0.015-0.017) |
| IC | 0.547 ±0.147 (0.381-0.663) |

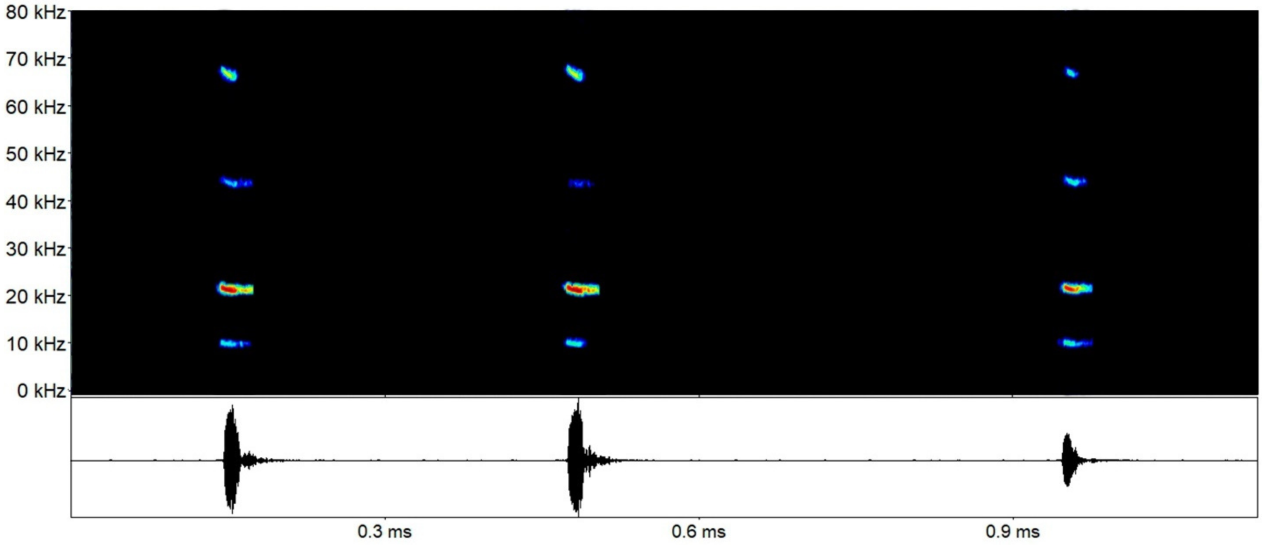

***Diclidurus ingens*** Hernández-Camacho, 1955: Composed by sequencies with a single call, although reported in the literature sequencies with two distinct calls (Arias-Aguilar et al., 2018). Calls with a quasi-constant (qCF) shape, and FME around 18.5 kHz.

| **Parameter** | **Call** |
|---|---|
| Finitial | 16.37 ±0.61 (15.73-17.10) |
| Ffinal | 20.09 ±1.00 (18.87-21.64) |
| Fmin | 17.88 ±0.84 (16.87-19.12) |
| Fmax | 19.00 ±0.96 (18.05-20.56) |
| FME | 18.50 ±0.98 (17.57-20.06) |
| BW | 1.11 ±0.20 (0.93-1.44) |
| Dur | 0.028 ±0.004 (0.02-0.03) |
| IC | 0.59 ±0.308 (0.17-0.94) |

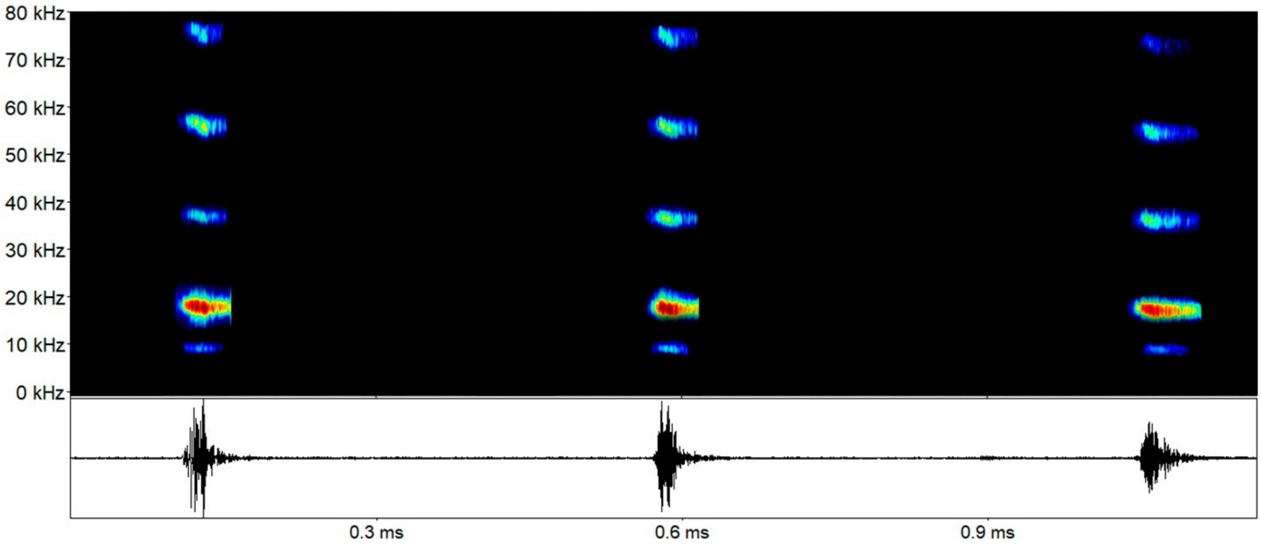

***Peropteryx kappleri*** Peters, 1867: Sequencies composed by a single call, with prominent harmonics. We extracted parameters for the first two harmonics, being the second that with most energy. A quasi-constant (qCF) shape, with FME around 30.8 kHz (30.8-31.6 kHz) in the second harmonic.

| **Parameter** | **1st harmonic** | **2nd harmonic** |
|---|---|---|
| Finitial | 13.10 ±1.49 (11.18-14.83) | 27.29 ±1.60 (27.29-28.97) |
| Ffinal | 17.33 ±1.16 (16.05-18.86) | 32.71 ±1.173 (32.71-33.69) |
| Fmin | 14.99 ±1.08 (14.17-16.56) | 29.93 ±0.65 (29.93-30.67) |
| Fmax | 16.16 ±1.11(15.1579-17.76) | 31.20 ±0.83 (31.20-32.10) |
| FME | 15.70 ±1.00 (14.84-17.14) | 30.80 ±0.80 (30.80-31.68) |
| BW | 1.17 ±0.32 (0.76-1.57) | 1.27 ±0.24 (1.27-1.50) |
| Dur | 0.014 ±0.007 (0.008-0.023) | 0.011 ±0.003 (0.01-0.017) |
| IC | - | 0.189 ±0.046 (0.171-0.248) |

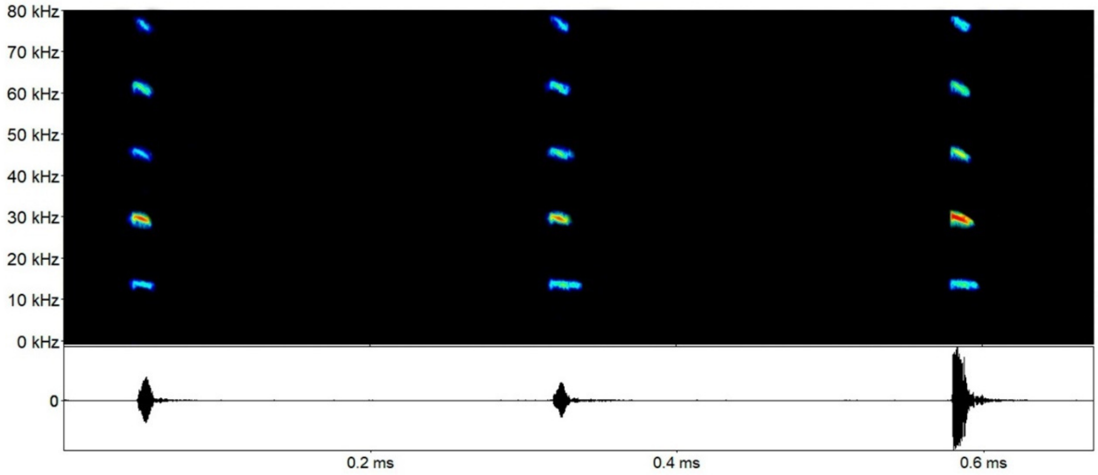

***Peropteryx macrotis*** (Wagner, 1843): Sequencies composed by a single quasi-constant (qCF) call, with FME around 40.2 kHz in the second harmonic (38.6-40.9 kHz).

| Parameter | 1st harmonic | 2nd harmonic |
|---|---|---|
| Finitial | 15.89 ±1.11 (13.99-18.02) | 36.84 ±1.46 (34.66-3.77) |
| Ffinal | 20.47 ±1.12 (18.02-21.86) | 42.05 ±0.79 (40.98-42.84) |
| Fmin | 17.08 ±0.56 (16.12-18.00) | 39.58 ±1.23 (37.78-40.53) |
| Fmax | 19.37 ±0.53 (18.00-19.87) | 40.64 ±1.05 (39.06-41.28) |
| FME | 18.83 ±0.55 (18.00-19.50) | 40.23 ±1.07 (38.64-40.91) |
| BW | 2.57 ±0.54 (1.87-3.37) | 1.05 ±0.22(0.75-1.27) |
| Dur | 0.009 ±0.002 (0.001-0.01) | 0.010 ±0.0004 (0.01-0.01) |
| IC | - | 0.180 ±0.036 (0.13-0.21) |

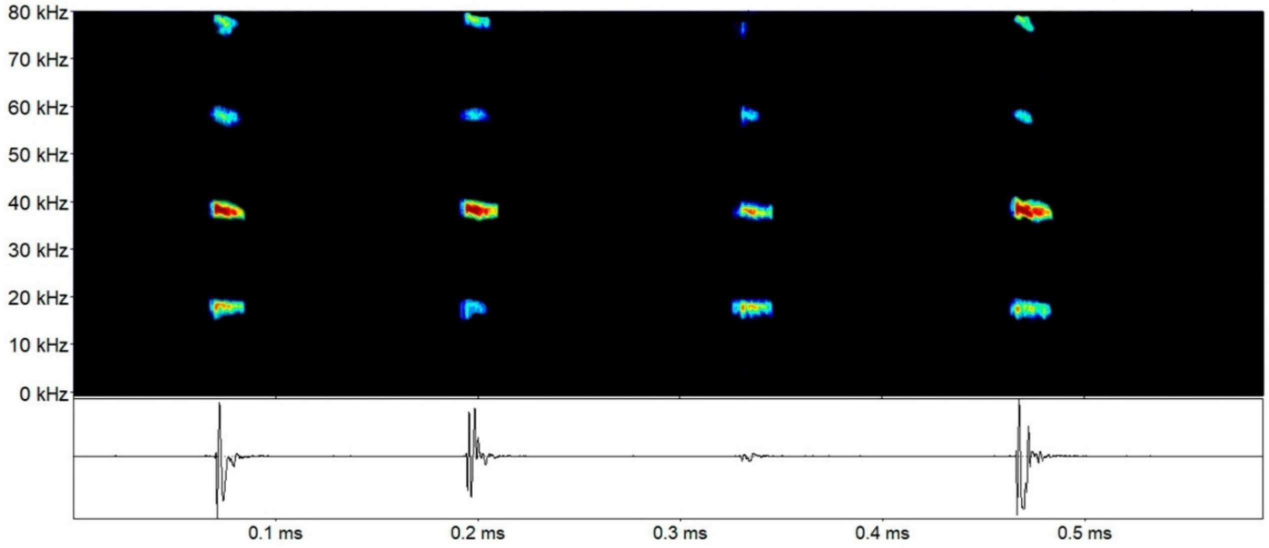

***Peropteryx trinitatis*** Miller, 1899: Sequencies composed by a single quasi-constant (qCF) call, with FME around 44.0 kHz in the second harmonic (40.8-44.6 kHz).

| Parameter | 1st harmonic | 2nd harmonic |
|---|---|---|
| Finitial | 19.47 ±1.58 (15.08-20.67) | 40.81 ±1.99 (34.32-42.95) |
| Ffinal | 24.13 ±0.68 (23.18-25.42) | 4.13 ±0.62 (45.06-47.36) |
| Fmin | 20.36 ±1.22 (17.25-21.37) | 43.25 ±1.13 (39.37-44.25) |
| Fmax | 22.68 ±0.36 (22.12-23.25) | 44.59 ±0.37 (48.87-45.00) |
| FME | 21.75 ±1.33 (18.00-22.50) | 44.07 ±0.76 (40.87-44.62) |
| BW | 2.32 ±1.12 (1.12-4.87) | 1.33 ±0.95 (0.75-4.87) |
| Dur | 0.008 ±0.001 (0.007-0.011) | 0.01 ±0.001 (0.007-0.014) |
| IC | - | 0.10 ±0.03 (0.09-0.21) |

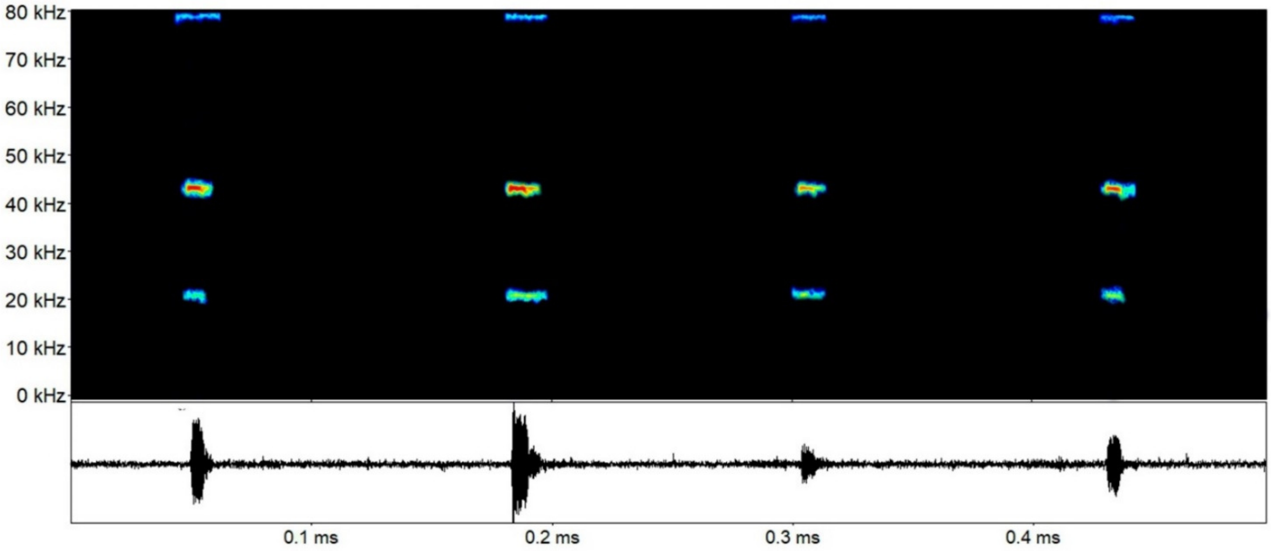

***Peropteryx* sp.**: Sequencies composed by a single quasi-constant (qCF) call, with FME around 41.1 kHz in the second harmonic (40.6-41.5 kHz).

| Parameter | 1st harmonic | 2nd harmonic |
|---|---|---|
| Finitial | 17.15 ±1.48 (15.44-18.10) | 37.54 ±0.92 (36.80-38.58) |
| Ffinal | 22.45 ±0.72 (21.73-23.17) | 42.90 ±0.92 (41.83-48.44) |
| Fmin | 19.00 ±0.83 (18.14-19.81) | 39.89 ±0.67 (39.11-40.28) |
| Fmax | 21.01 ±0.27 (20.81-21.33) | 41.38 ±1.00 (40.31-42.31) |
| FME | 20.40 ±0.34 (20.06-20.75) | 41.11 ±0.44 (40.68-41.58) |
| BW | 2.00 ±0.65 (1.52-2.75) | 1.49 ±0.47 (1.20-2.04) |
| Dur | 0.013 ±0.002 (0.01-0.016) | 0.013 ±0.002 (0.001-0.016) |
| IC | - | 0.098 ±0.006 (0.091-0.102) |

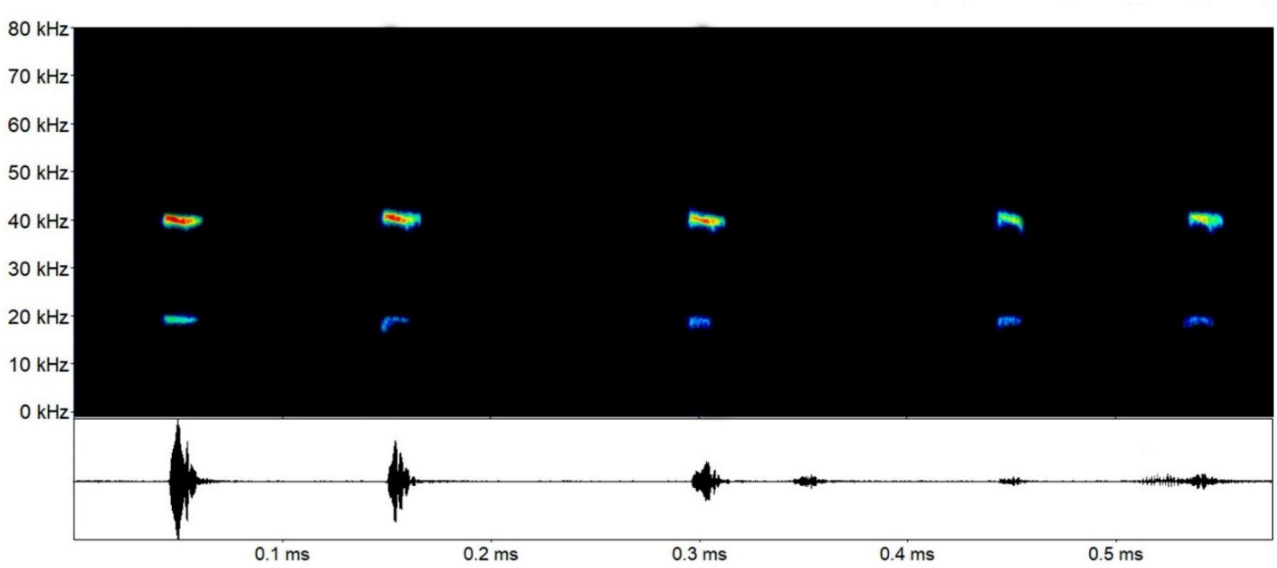

***Rhynchonycteris naso*** (Wied, 1820): Sequencies with calls with two harmonics, sometimes with one of them absent; usually the second harmonic is the most evident. Calls have a quasi-constant (qCF) part followed by a FM part, sometimes not as evident or even absent. FME around 43.9 kHz (39.7-46.3 kHz) in the first harmonic, and 93.5 kHz (88.3-97.0 kHz) in the second.

| Parameter | 1st harmonic | 2nd harmonic |
|---|---|---|
| Finitial | 33.25 ±2.18 (30.97-36.16) | 72.00 ±3.71 (67.95-77.13) |
| Ffinal | 49.88 ±0.24 (49.50-50.13) | 98.02 ±0.51 (97.39-98.51) |
| Fmin | 35.15 ±1.99 (32.96-37.75) | 79.32 ±4.24 (72.27-83.11) |
| Fmax | 48.27 ±0.23 (48.06-48.64) | 96.83 ±0.56 (96.15-97.45) |
| FME | 43.97 ±2.65 (39.71-46.39) | 93.57 ±3.36 (88.39-97.01) |
| BW | 13.11 ±2.00 (10.37-15.10) | 17.51 ±4.51 (13.22-24.64) |
| Dur | 0.006 ±0.001 (0.006-0.009) | 0.007 ±0.001 (0.006-0.01) |
| IC | 0.037 ±0.006 (0.03-0.04) | - |

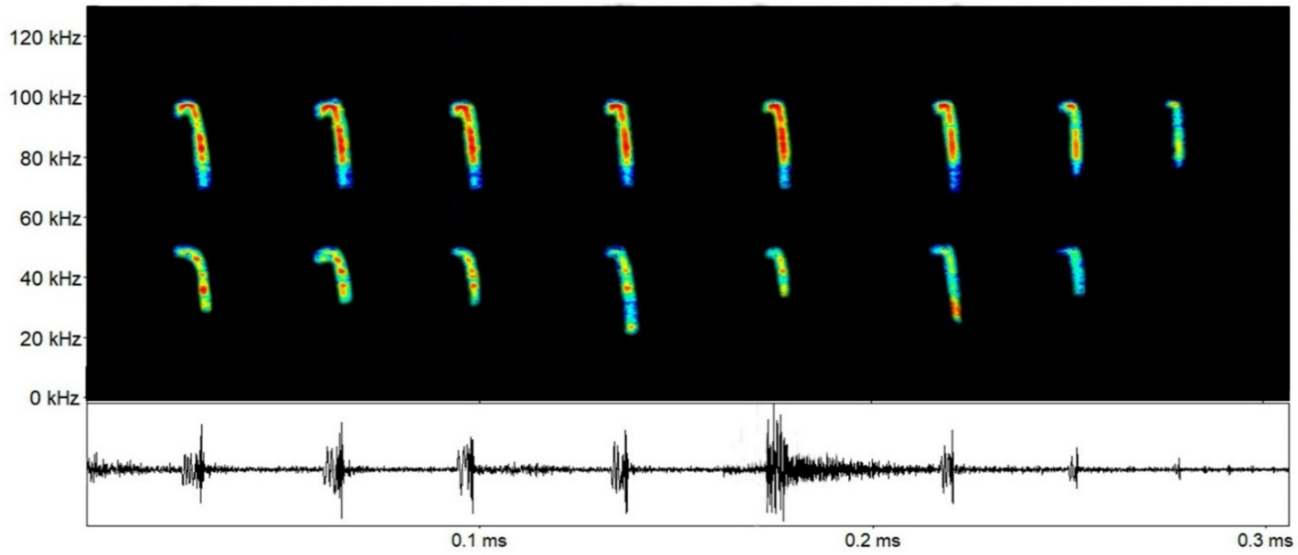

***Saccopteryx bilineata*** (Temminck, 1838): Sequencies have usually two alternating quasi-constant (qCF) ascendent calls (types a and b), but sometimes one of them can be absent. Harmonics sometimes are not distinguishable, but the second one is more evident, with FME around 41.4 kHz (40.3-42.7 kHz) for call type a, and 44.1 kHz (43.3-45.3 kHz) for call type b. However, in other parts of Brazil, reported FME for *S. bilineata* are around 44 kHz for call type a, and 48 kHz for call type b.

| Parameter | Call a | Call b |
|---|---|---|
| Finitial | 36.78 ±1.53 (34.84-38.54) | 39.74 ±1.170 (38.15-40.70) |
| Ffinal | 43.36 ±0.88 (42.27-44.66) | 45.92 ±1.08 (42.33-44.22) |
| Fmin | 40.20 ±0.83 (39.03-41.39) | 43.05 ±0.77 (42.33-44.22) |
| Fmax | 42.14 ±0.98 (40.83-43.57) | 44.69 ±0.83 (43.87-45.86) |
| FME | 41.49 ±0.85 (40.38-42.72) | 44.17 ±0.89 (43.37-45.37) |
| BW | 1.93 ±0.26 (1.54-2.17) | 1.63 ±0.11 (1.45-1.79) |
| Dur | 0.013 ±0.002 (0.009-0.015) | 0.012 ±0.002 (0.008-0.014) |
| IPI | 0.058 ±0.010 (0.048-0.070) | - |
| IC | 0.096 ±0.020 (0.071-0.128) | - |

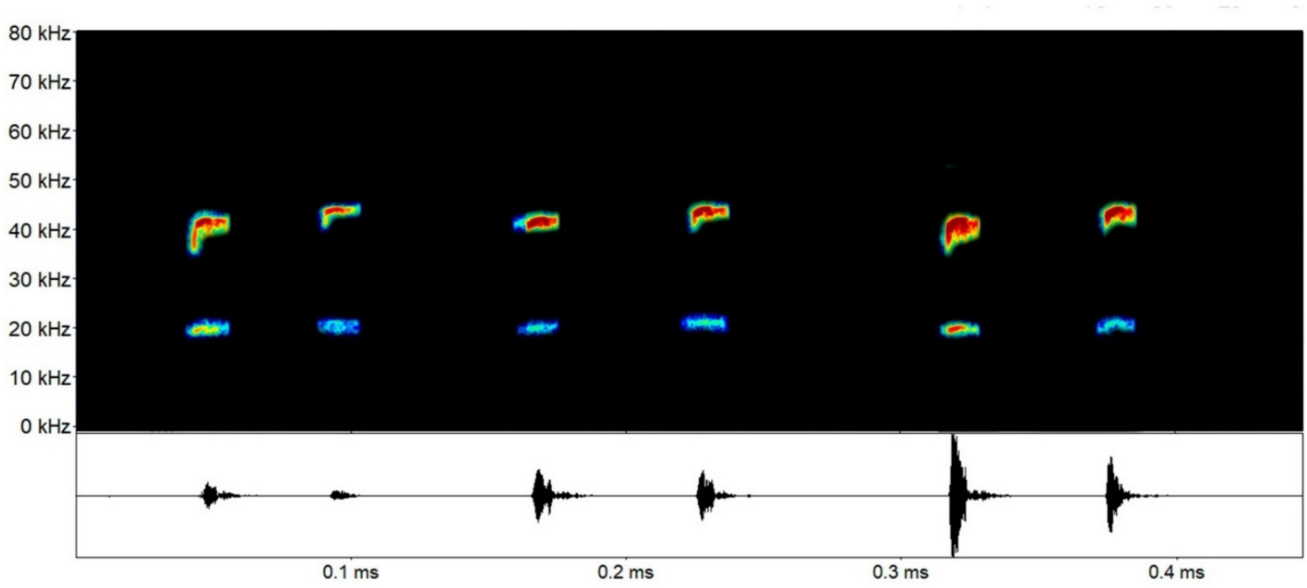

***Saccopteryx leptura*** (Schreber, 1774): Sequencies have usually two alternating quasi-constant (qCF) ascendent calls (a and b), but sometimes one of them can be absent. Harmonics sometimes are not distinguishable, but the second one is more evident, with FME around 49.9 kHz (48.8-51.7 kHz) for call type a, and 52.6 kHz (51.8-54.1 kHz) for call type b.

| Parameter | Call a | Call b |
|---|---|---|
| Finitial | 44.75 ±1.28 (43.30-46.34) | 48.08 ±1.18 (46.81-49.62) |
| Ffinal | 51.91 ±1.19 (51.22-53.69) | 54.34 ±0.94 (53.57-55.70) |
| Fmin | 48.13 ±0.94 (47.14-49.35) | 51.39 ±1.11 (50.40-52.91) |
| Fmax | 50.72 ±1.16 (49.98-52.46) | 53.22 ±1.02 (52.44-54.71) |
| FME | 49.90 ±1.29 (48.85-51.78) | 52.69 ±0.98 (51.88-54.11) |
| BW | 2.58 ±0.59 (1.91-3.11) | 1.82 ±0.33 (1.50-2.28) |
| Dur | 0.008 ±0.003 (0.005-0.012) | 0.007 ±0.001 (0.005-0.009) |
| IPI | 0.036 ±0.004 (0.033-0.043) | |
| IC | 0.061 ±0.009 (0.048-0.069) | |

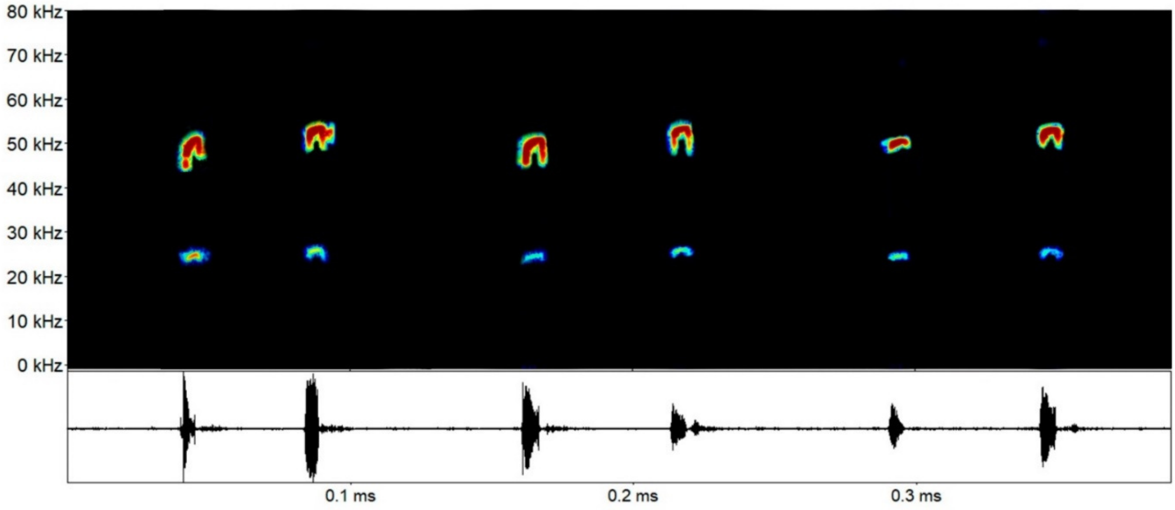

#### Family Phyllostomidae

***Lonchorhina aurita*** Tomes, 1863: Calls with harmonics, but sometimes hard to visualize. In our analysis, the third harmonic was the most evident, as reported in the literature. Calls have a short quasi-constat part (qCF), followed by a FM part. FME around 47.7 kHz for the qCF part (46.87-48.62 kHz).

| Parameter | Call |
|---|---|
| Finitial | 37.92 ±0.54 (37.12-38.32) |
| Ffinal | 49.83 ±0.60 (49.00-50.44) |
| Fmin | 41.29 ±0.62 (40.71-41.91) |
| Fmax | 48.64 ±0.31 (48.30-49.00) |
| FME | 47.73 ±0.81 (46.87-48.62) |
| BW | 7.34 ±0.37 (6.87-7.76) |
| Dur | 0.01 ±0.001 (0.008-0.011) |
| IC | 0.095 ±0.031 (0.069-0.139) |

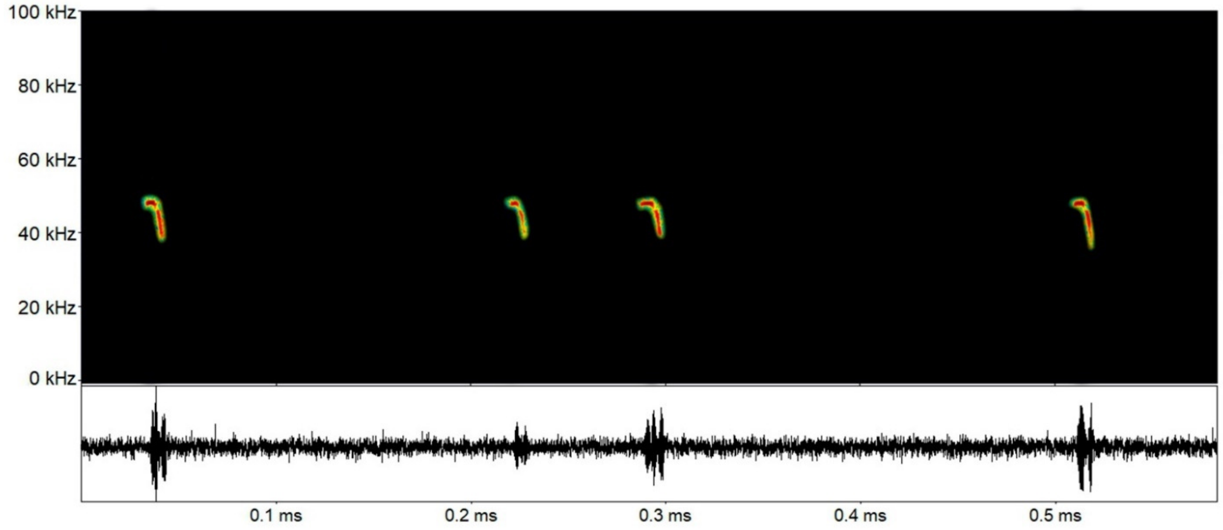

#### Family Mormoopidae

Echolocation calls for *Pteronotus gymnonotus* (Pgym), *Pteronotus personatus* (Pper), *Pteronotus rubiginosus* (Prub), and *Pteronotus alitonus* (Pali) from Carajás region, Brazilian Amazonia.

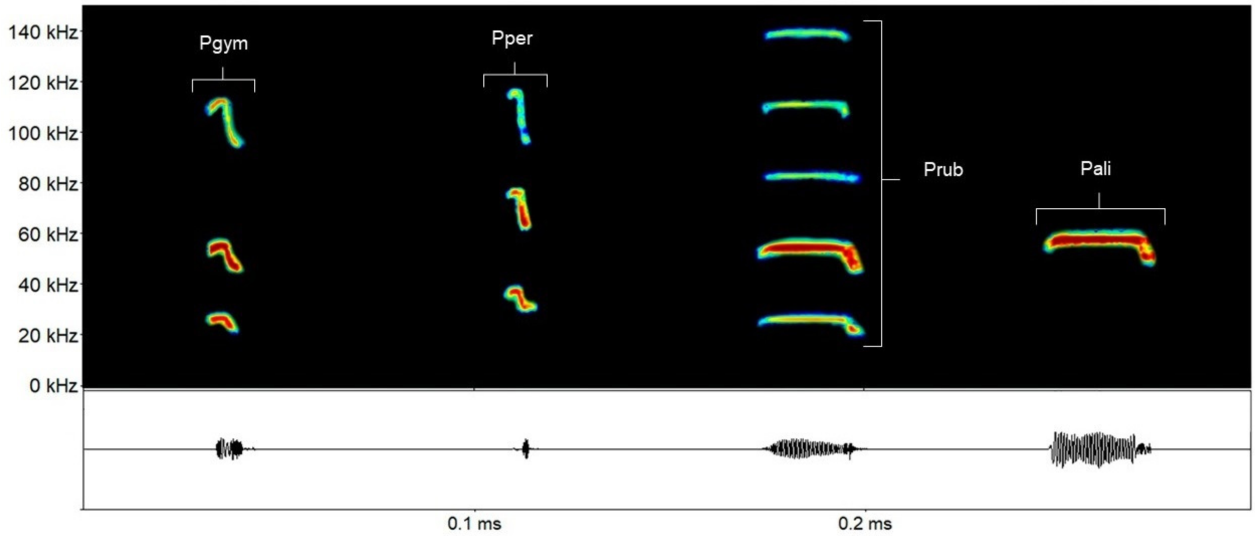

***Pteronotus personatus*** (Wagner, 1843): This is a unique call with a typical evident lazy Z shape, with many harmonics, being the second the most evident, with FME around 64.3 kHz (62.6-67.5 kHz).

| Parameter | 1st harmonic | 2nd harmonic |
|---|---|---|
| Finitial | 30.22 ±0.63 (29.67-30.91) | 61.34 ±0.59 (60.55-61.93) |
| Ffinal | 38.07 ±0.36 (37.76-38.60) | 74.42 ±0.70 (73.49-75.11) |
| Fmin | 31.75 ±0.73 (31.00-32.62) | 62.95 ±0.69 (62.20-63.63) |
| Fmax | 37.11 ±0.47 (36.62-37.75) | 70.26 ±2.34 (68.02-73.42) |
| FME | 35.02 ±2.67 (31.25-37.50) | 64.31 ±2.03 (62.62-67.57) |
| BW | 5.36 ±0.30 (5.07-5.62) | 7.31 ±1.88 (5.51-9.97) |
| Dur | 0.007 ±0.001 (0.006-0.009) | 0.009 ±0.001 (0.007-0.011) |
| IC | - | 0.062 ±0.012 (0.048-0.073) |

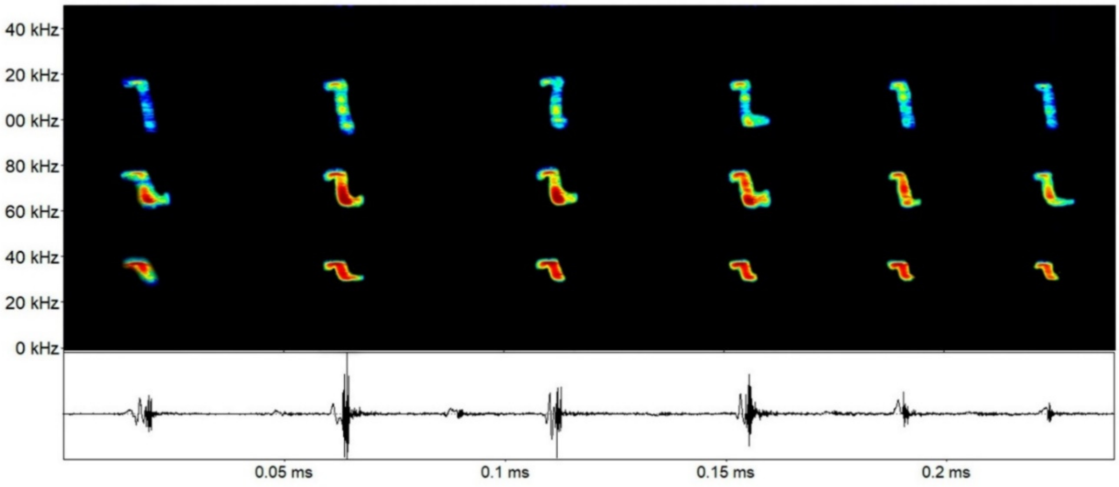


.

***Pteronotus gymnonotus*** (Wagner, 1843): This is a unique call with a typical evident lazy Z shape, with many harmonics, being the second the most evident, with FME around 53.8 kHz (51.6-55.4 kHz).

| Parameter | 1st harmonic | 2nd harmonic |
|---|---|---|
| Finitial | 22.71 ±0.32 (22.34-22.92) | 45.55 ±1.19 (44.49-46.80) |
| Ffinal | 29.30 ±0.62 (28.66-29.90) | 57.54 ±0.75 (56.49-58.19) |
| Fmin | 23.78 ±0.54 (23.43-24.41) | 48.48 ±0.95 (47.25-49.57) |
| Fmax | 28.25 ±0.28 (27.93-28.50) | 56.34 ±0.72 (55.32-57.00) |
| FME | 27.60 ±0.72 (26.81-28.23) | 53.87 ±1.96 (51.66-55.42) |
| BW | 4.46 ±0.39 (4.08-4.8) | 7.85 ±0.29 (7.42-8.07) |
| Dur | 0.008 ±0.009 (0.008-0.01) | 0.01 ±0.0008 (0.009-0.011) |
| IC | - | 0.098 ±0.047 (0.073-0.169) |

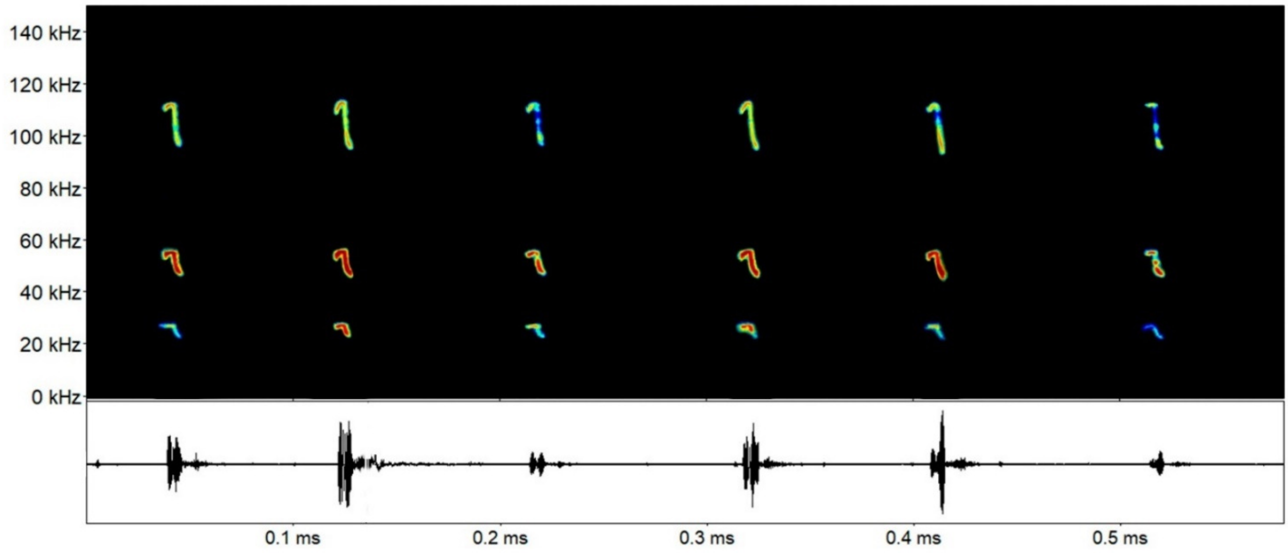

***Pteronotus rubiginosus*** (Wagner, 1843): The striking characteristic of these signals is the long duration CF component (25-37 ms), with multiple harmonics, being the second that with most energy (FME around 55.6 kHz (55.2-56.0 kHz)).

| Parameter | 1st harmonic | 2nd harmonic |
|---|---|---|
| Finitial | 24.33 ±2.38 (21.77-26.49) | 47.80 ±1.61 (45.39-48.74) |
| Ffinal | 29.24 ±0.37 (28.90-29.64) | 57.17 ±0.74 (54.46-56.19) |
| Fmin | 26.17 ±1.40 (24.61-27.34) | 54.67 ±0.66 (53.81-55.37) |
| Fmax | 28.19 ±0.18 (28.06-28.40) | 55.98 ±0.36 (55.59-56.40) |
| FME | 27.61 ±0.33 (27.30-27.96) | 55.63 ±0.36 (55.23-56.09) |
| BW | 2.02 ±1.30 (1.06-3.50) | 1.31 ±0.46 (0.81-1.78) |
| Dur | 0.028 ±0.001 (0.025-0.029) | 0.031 ±0.004 (0.025-0.037) |
| IC | - | 0.032 ±0.008 (0.21-0.041) |

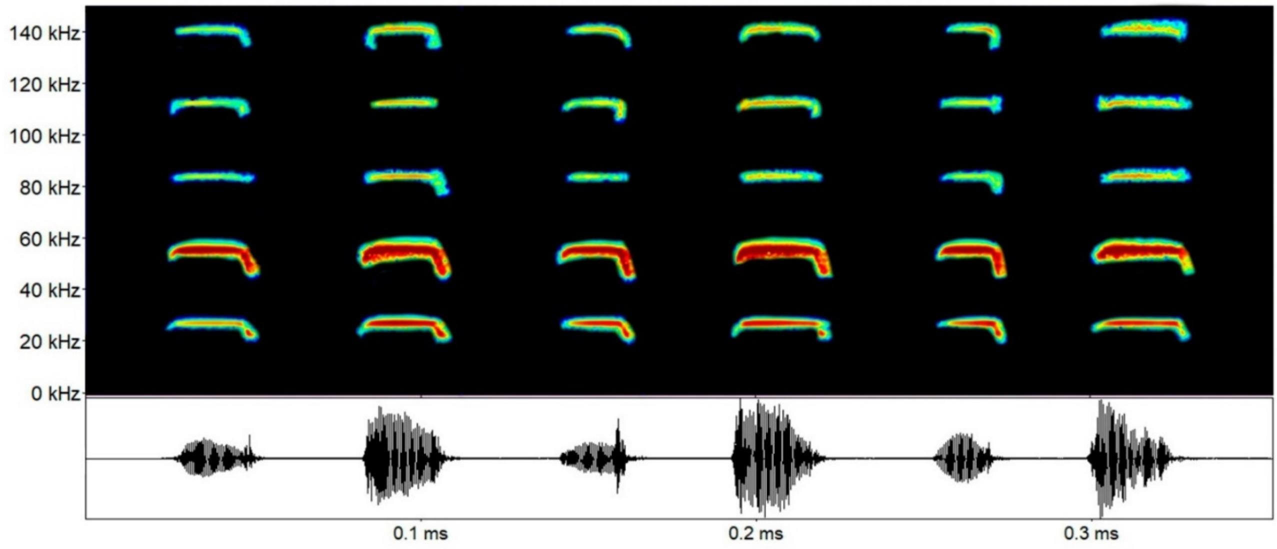

***Pteronotus alitonus*** Pavan, Bobrowiec e Percequillo, 2018: The striking characteristic of these signals is the long duration CF component (15-31 ms), with multiple harmonics, being the second that with most energy (FME around 59.2 kHz (58.9-59.5 kHz)).

| Parameter | 2nd harmonic |
|---|---|
| Finitial | 50.42 ±23.96 (48.75-53.98) |
| Ffinal | 60.34 ±32.53 (59.92-60.61) |
| Fmin | 55.81 ±25. (52.90-59.00) |
| Fmax | 59.51 ±21.16 (59.28-59.78) |
| FME | 59.27 ±26.36 (58.98-59.57) |
| BW | 37.45 ±24.34 (77.67-66.50) |
| Dur | 0.022 ±0.006 (0.015-0.31) |
| IC | 0.028 ±0.012 (0.017-0.043) |

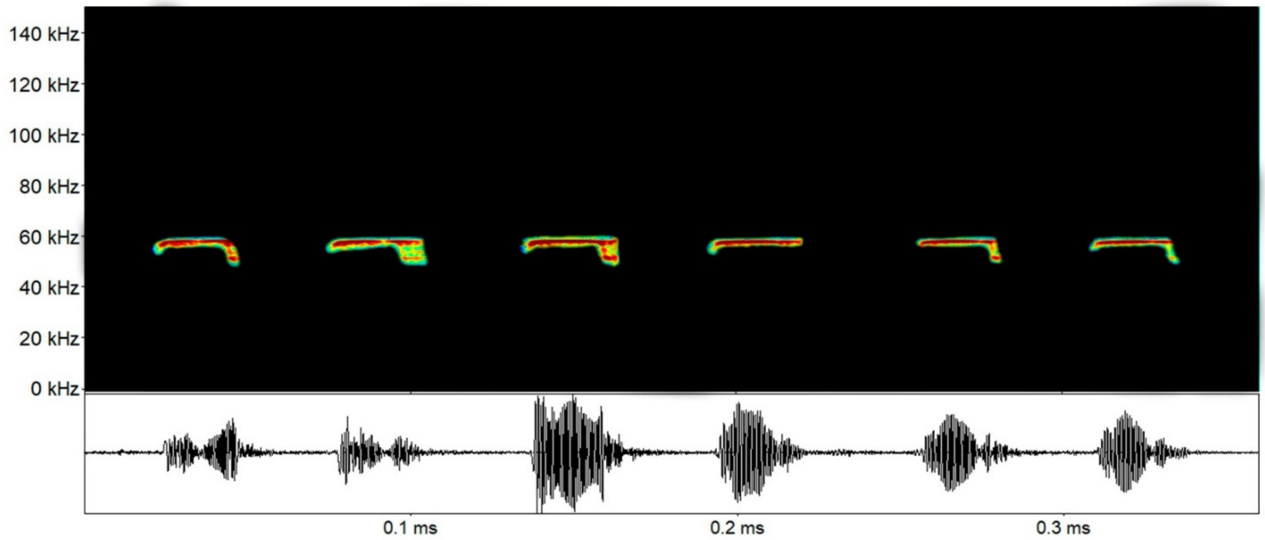

#### Family Noctilionidae

Echolocation calls for *Noctilio albiventris* (Nalb) and *Noctilio leporinus* (Nlep) from the Carajás region, Brazilian Amazonia.

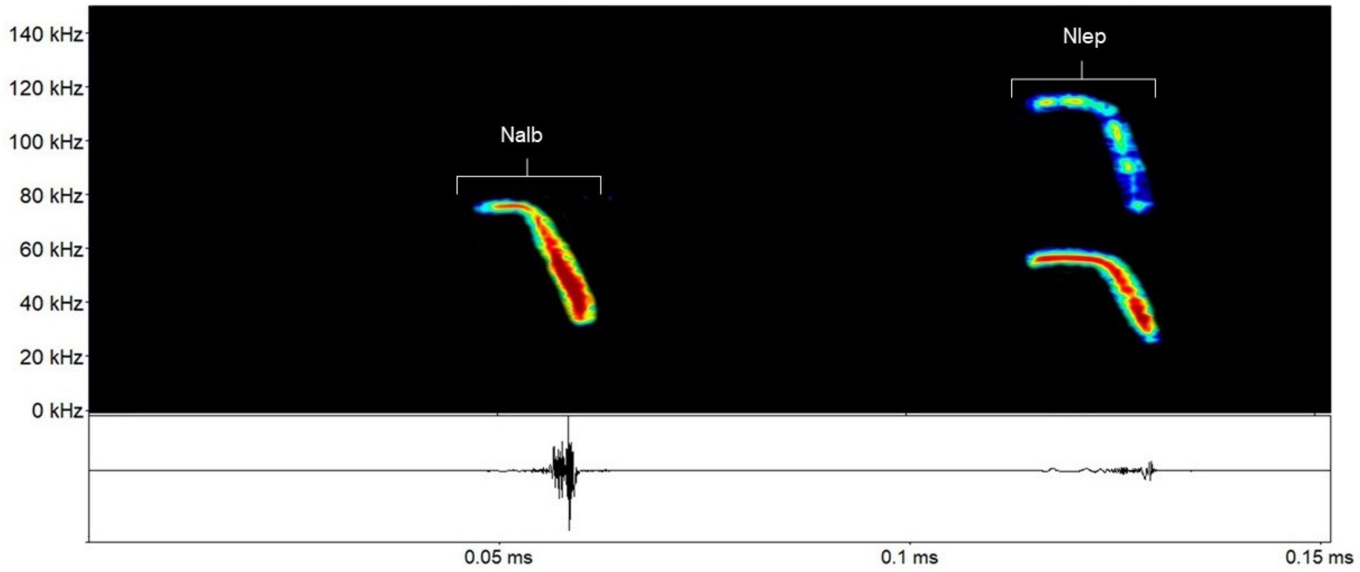

*Noctilio albiventris* Desmarest, 1818: Characterized by calls with a qCF part followed by a steep FM part. We identified call type a, with an evident descendent FM part, and call type b, for those with the FM part not evident. FME for type a was around 44.9 kHz (39.5-48.0 kHz), and for type b around 71.2 kHz (67.6-73.3 kHz. A medida de frequência quase constante (qCF) do sinal tem frequência de máxima de energia em torno de 78-80 kHz.

| Parâmetro | Sinal a | Sinal b |
|---|---|---|
| Finicial | 34.03 ±4.54 (28.87-37.43) | 62.81 ±3.22 (59.41-65.81) |
| Ffinal | 74.22 ±4.06 (69.53-76.58) | 74.05 ±3.67 (69.85-76.64) |
| Fmin | 41.20 ±4.20 (36.37-44.08) | 66.82 ±4.011 (62.73-70.75) |
| Fmax | 53.43 ±5.57 (47.06-57.42) | 72.81 ±4.00 (68.19-75.37) |
| FME | 44.89 ±4.67 (39.52-48.00) | 71.24 ±3.16 (67.60-73.37) |
| BW | 12.23 ±1.37 (10.68-13.33) | 5.98 ±1.68 (4.60-7.87) |
| Dur | 0.014 ±0.0008 (0.013-0.015) | 0.012 ±0.009 (0.011-0.013) |
| IC | 0.063 ±0.021 (0.047-0.087) | |

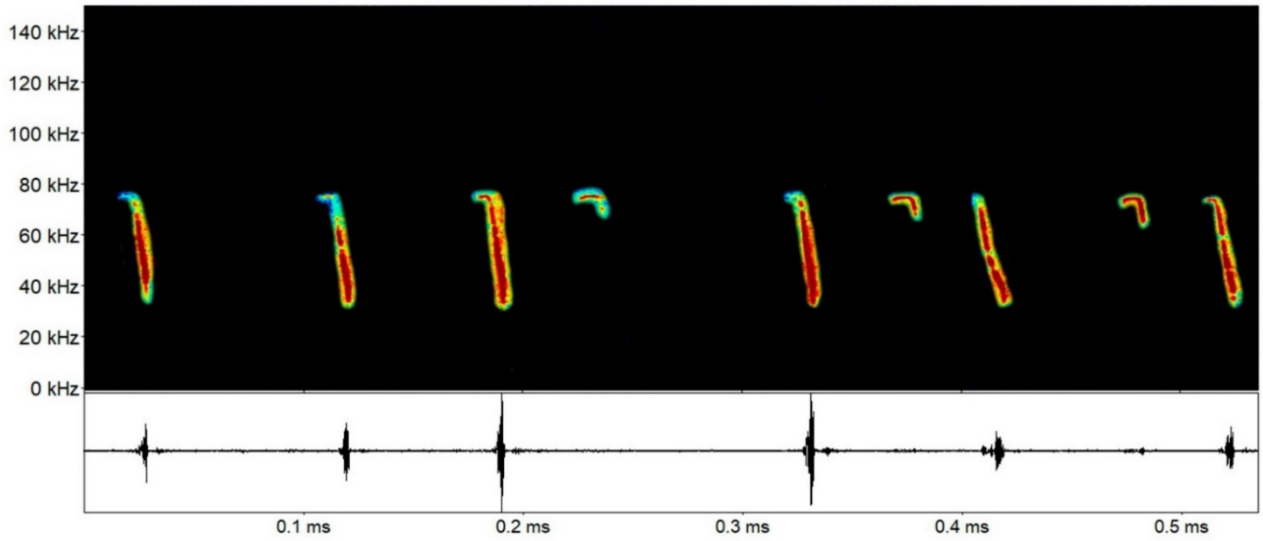

*Noctilio leporinus* (Linnaeus, 1758): É um sinal único, com frequencia quase constante (qCF) e modulação de frequência (FM) (Figura 22). É um sinal que possui harmônicos, sendo o primeiro harmônico mais evidente no sinal. A frequência de máxima energia (FME) do primeiro harmônico completo variou de 33-41 kHz (Tabela 19). A medida de frequência quase constante do sinal tem frequência de máxima de energia em torno de 55 kHz.

| Parâmetro | Sinal a |
|---|---|
| Finicial | 22.36 ±4.97 (16.65-25.73) |
| Ffinal | 56.72 ±1.59 (55.08-58.25) |
| Fmin | 28.63 ±5.00 (23.60-33.60) |
| Fmax | 49.20 ±7.41 (40.67-54.08) |
| FME | 37.19 ±4.11 (33.08-41.32) |
| BW | 20.56 ±4.31 (17.07-25.38) |
| Dur | 0.013 ±0.0009 (0.012-0.014) |
| IC | 0.109 ±0.078 (0.047-0.198) |

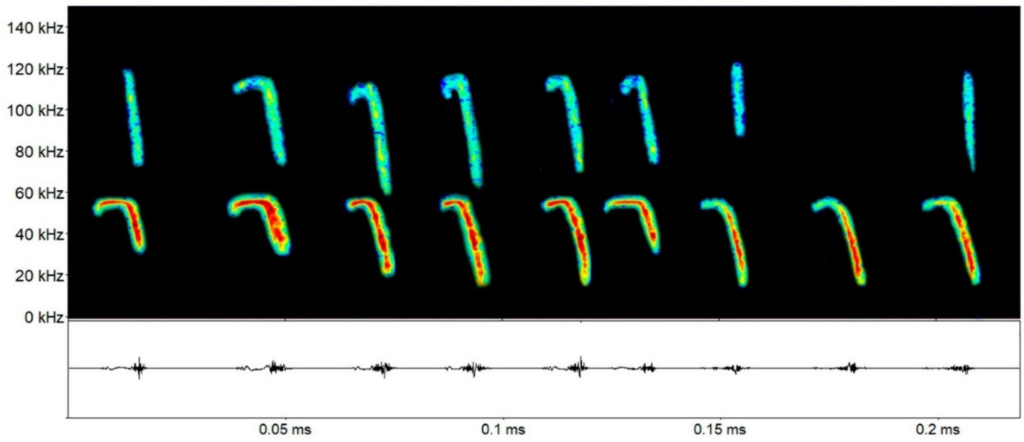

### Family Thyropteridae

Echolocation calls for *Thyroptera* sp. 1 (Tsp1), *Thyroptera* sp. 2 (Tsp2), *Thyroptera tricolor* (Ttri) and *Thyroptera* sp. 3 (Tsp3) from the Carajás region, Brazilian Amazonia.

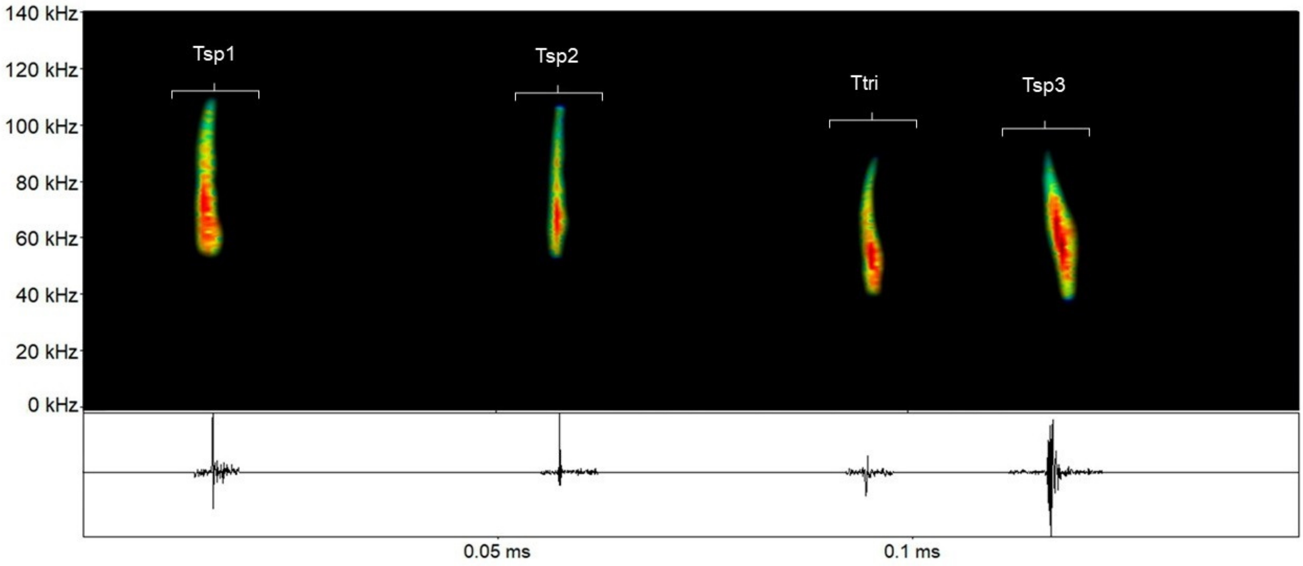

***Thyroptera* sp. 1**: Very short FM calls (1-3 ms), with broad amplitude (51.5 up to 106.3 kHz), usually in sequences with several calls with short interval between them (1-4ms). FME around 67.3 kHz, varying from 60.3 up to 73.8 kHz.

| Parameter | Call |
|---|---|
| Finitial | 54.26 ±2.18 (51.54-59.95) |
| Ffinal | 101.48 ±4.48 (92.76-106.32) |
| Fmin | 58.42 ±1.47 (55.12-61.50) |
| Fmax | 81.11 ±3.24 (76.87-88.50) |
| FME | 67.36 ±4.84 (60.37-73.87) |
| BW | 22.68 ±3.48 (18.37-30.00) |
| Dur | 0.002 ±0.0003 (0.001-0.003) |
| IC | 0.0217 ±0.006 (0.014-0.041) |

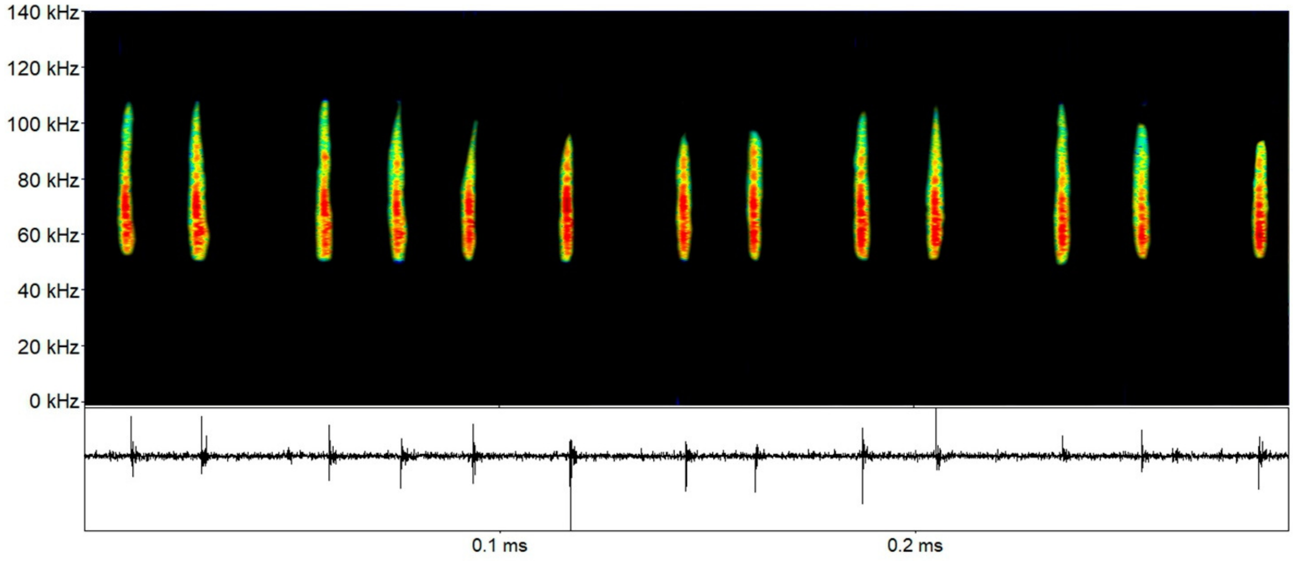

***Thyroptera* sp. 2**: Very short FM calls (around 2 ms) with broad amplitude (58.8 up to 105.4 kHz). FME around 65.7 kHz, varying from 62.6 up to 69.0 kHz.

| Parameter | Call |
|---|---|
| Finitial | 57.48 ±1.59 (55.22-59.44) |
| Ffinal | 97.87±5.55 (86.29-105.46) |
| Fmin | 61.53 ±1.32 (58.87-63.00) |
| Fmax | 79.98 ±4.98 (73.12-89.62) |
| FME | 65.70 ±2.19 (62.62-69.00) |
| BW | 18.45 ±5.15 (10.87-27.00) |
| Dur | 0.002 ±0.0002 (0.002-0.0028) |
| IC | 0.049 ±0.030 (0.022-0.125) |

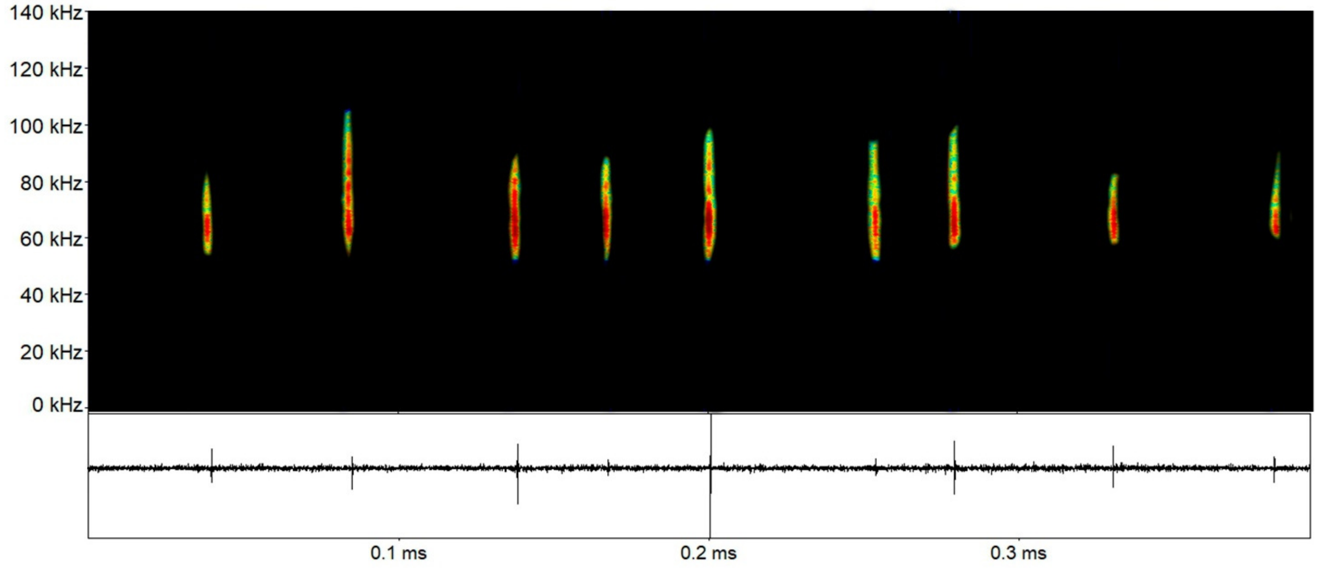

***Thyroptera* sp. 3**: Very short FM calls (around 3-5 ms) with broad amplitude (49.5 up to 90.1 kHz). FME around 61.5 kHz, varying from 57.0 up to 67.8 kHz.

| Parameter | Call |
|---|---|
| Finitial | 46.35 ±2.51 (42.95-49.85) |
| Ffinal | 85.53 ±3.23 (76.70-90.12) |
| Fmin | 53.82 ±1.71 (49.50-55.87) |
| Fmax | 66.52 ±1.26 (64.50-69.37) |
| FME | 61.52 ±3.23 (57.00-67.87) |
| BW | 12.7 ±1.87 (9.75-16.50) |
| Dur | 0.004 ±0.0006 (0.003-0.005) |
| IC | 0.042 ±0.013 (0.023-0.075) |

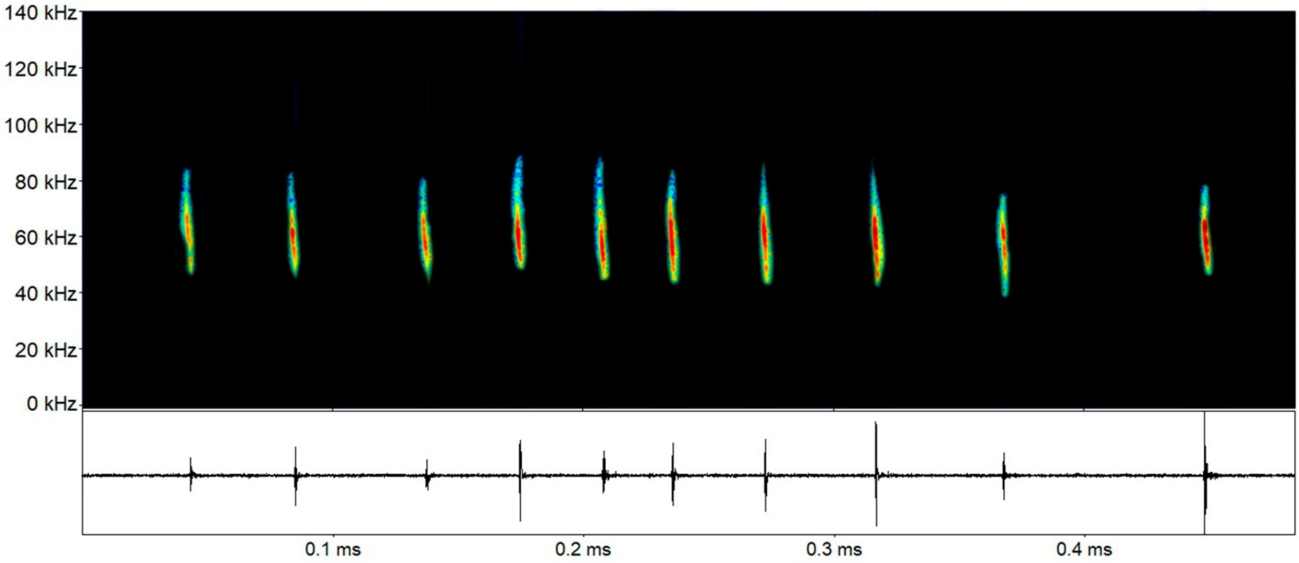

***Thyroptera tricolor*** (Spix, 1823): Short FM calls (around 2-3 ms) with broad amplitude (39.3 up to 78.2 kHz). FME around 55.1 kHz, varying from 50.2 up to 57.3 kHz.

| Parameter | Call |
|---|---|
| Finitial | 42.81 ±3.36 (35.28-49.47) |
| Ffinal | 71.12 ±4.82 (64.43-78.23) |
| Fmin | 47.49 ±3.26 (39.37-52.50) |
| Fmax | 60.65 ±1.72 (58.12-64.87) |
| FME | 55.18 ±2.16 (50.25-57.37) |
| BW | 13.16 ±3.80 (8.25-22.50) |
| Dur | 0.003 ±0.0004 (0.002-0.003) |
| IC | 0.012 ±0.004 (0.009-0.022) |

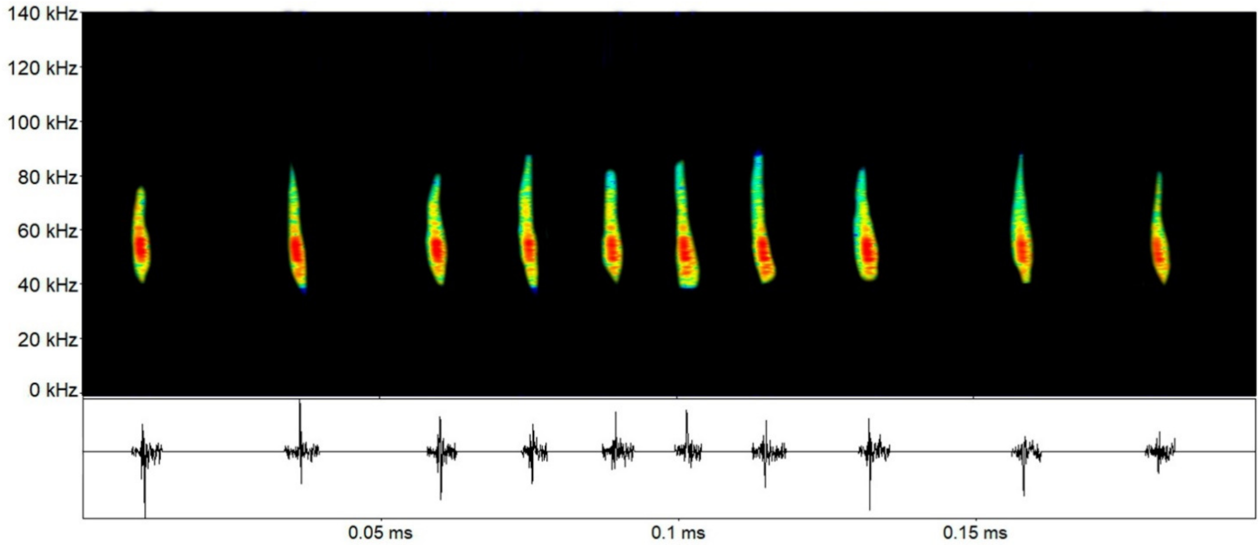

### Family Molossidae

Echolocation calls for *Cynomops* sp. 1 (Csp1), *Cynomops* sp. 2 (Csp2), *Cynomops* cf. *greenhalli* (Cgre), *Cynomops* cf. *planirostris* (Cpla), *Eumops* sp. 1 (Esp1), *Eumops* cf. *dabbenei* (Edab), *Eumops* spp. (Espp), *Molossus* cf. *currentium* (Mcur), *Molossus molossus* (Mmol), *Molossus rufus* (Mruf), *Molossops neglectus* (Mneg), and *Promops centralis* (Pcen) recorded in Carajás region, Brazilian Amazonia.

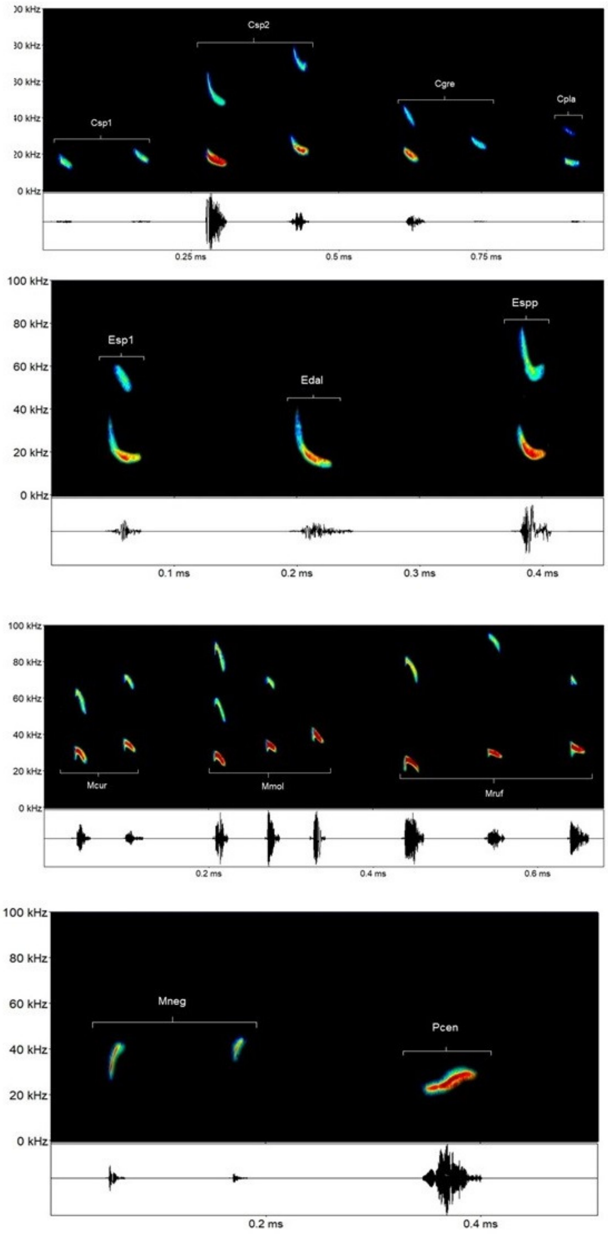

***Cynomops* sp. 1**: Sequencies with two alternating quasi-constant (qCF) descendent calls (call type a lower, and type b higher), sometimes with only one. FME for type a around 16.5 kHz (13.8-18.5 kHz), and around 20.3 kHz (16.6-23.2 kHz) for type b.

| Parameter | Call a | Call b |
|---|---|---|
| Finitial | 18.30 ±1.82 (15.25-20.91) | 22.07 ±3.24 (18.31-25.00) |
| Ffinal | 13.52 ±1.13 (11.78-15.44) | 16.78 ±1.52 (14.74-18.38) |
| Fmin | 15.87 ±1.54 (131.12-17.62) | 18.68 ±2.30 (16.00-21.62) |
| Fmax | 14.67 ±1.34 (12.50-16.56) | 17.78 ±1.73 (15.37-19.50) |
| FME | 16.509 ±1.54 (13.87-18.50) | 20.35 ±2.96 (16.62-23.25) |
| BW | 1.83 ±0.66 (1.12-3.07) | 2.56 ±1.48 (1.25-3.93) |
| Dur | 0.021 ±0.004 (0.017-0.030) | 0.019 ±0.004 (0.015-0.025) |
| IPI | 0.249 ±0.066 (0.158-0.319) | - |
| IC | 0.524 ±0.167 (0.239-0.849) | - |

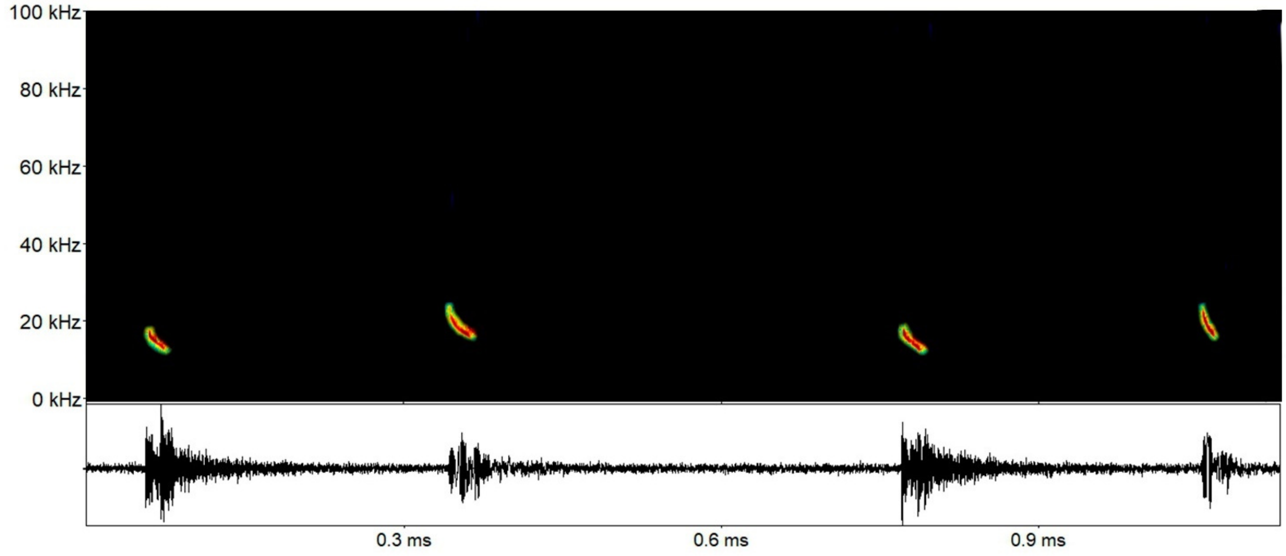

***Cynomops* sp. 2**: Sequencies with two alternating quasi-constant (qCF) descendent calls (call type a lower, and type b higher), sometimes with only one. FME for type a around 19.3 kHz (16.2-20.7 kHz), and around 23.0 kHz (19.4-26.8 kHz) for type b.

| Parameter | Call a | Call b |
|---|---|---|
| Finitial | 21.56 ±1.58 (17.07-24.84) | 25.79 ±2.33 (21.76-31.71) |
| Ffinal | 14.47 ±1.09 (12.17-16.53) | 19.51 ±1.47 (17.09-22.66) |
| Fmin | 17.61 ±1.14 (14.62-19.31) | 21.29 ±1.66 (18.00-24.75) |
| Fmax | 16.11 ±1.01 (13.75-18.00) | 20.45 ±1.58 (17.71-23.75) |
| FME | 19.31 ±1.04 (16.25-20.75) | 23.05 ±1.79 (19.40-26.87) |
| BW | 3.19 ±0.53 (2.00-4.50) | 2.59 ±0.80 (2.00-3.70) |
| Dur | 0.023 ±0.003 (0.012-0.028) | 0.022 ±0.002 (0.016-0.027) |
| IPI | 0.179 ±0.051 (0.096-0.302) | - |
| IC | 0.426 ±0.168 (0.141-0.879) | - |

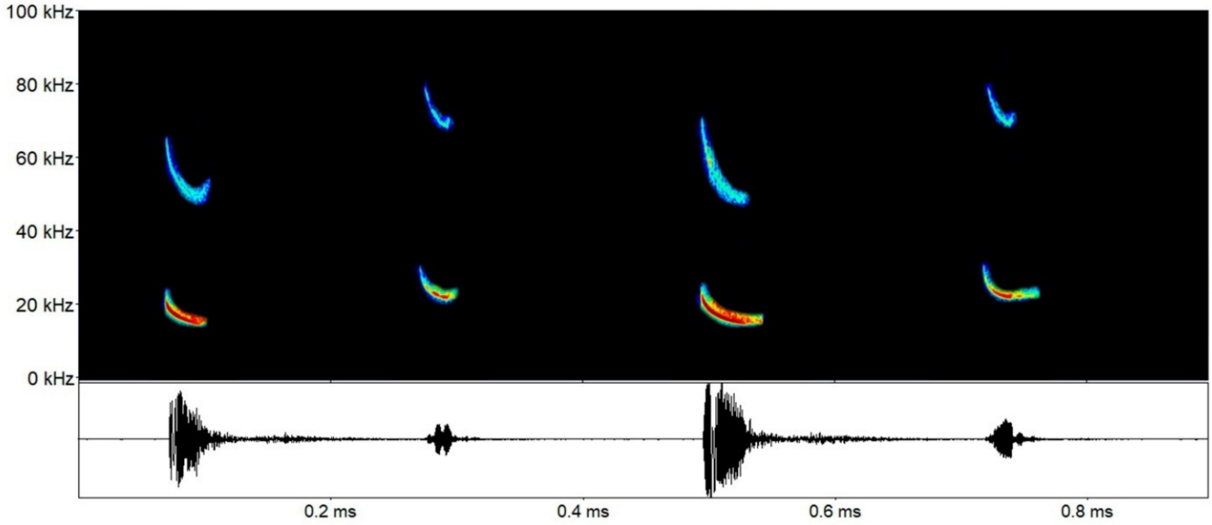

***Cynomops* cf. *greenhalli*** Goodwin, 1985: Sequencies with two alternating shallow descendent FM calls (types a and b), sometimes with only one. Harmonics are not easily distinguishable. FME around 23.1 kHz for type a calls (19.7-26.5 kHz), and around 27.2 kHz for type b (23.5-32.4 kHz).

| Parameter | Call a | Call b |
|---|---|---|
| Finitial | 25.89 ±1.60 (23.56-29.53) | 29.34 ±2.36 (25.82-35.48) |
| Ffinal | 17.82 ±2.69 (11.34-21.85) | 23.11 ±2.15 (19.21-27.82) |
| Fmin | 21.04 ±2.03 (17.12-24.65) | 25.29 ±2.28 (21.56-30.75) |
| Fmax | 19.51 ±2.23 (15.15-23.62) | 24.19 ±2.18 (20.81-29.47) |
| FME | 23.11 ±1.73 (19.79-26.53) | 27.24 ±2.21 (23.56-32.40) |
| BW | 3.60 ±1.04 (2.35-7.40) | 3.05 ±0.47 (2.34-4.03) |
| Dur | 0.019 ±0.003 (0.007-0.026) | 0.019 ±0.003 (0.012-0.025) |
| IPI | 0.115 ±0.021 (0.090-0.181) | - |

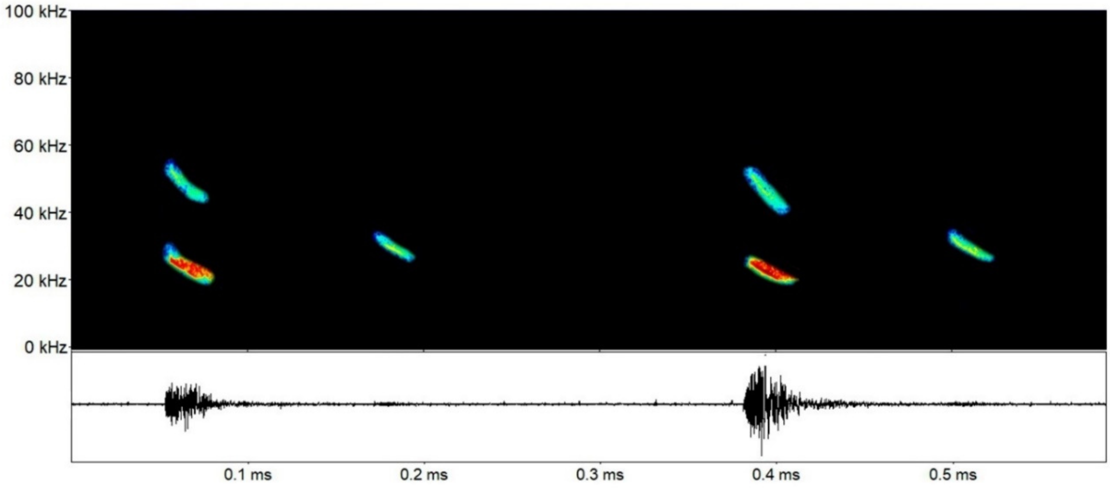

***Cynomops* cf. *planirostris*** (Peters, 1866): Sequencies with a single qCF descendent call, with harmonics, and FME around 28.1 kHz (27.6-28.6 kHz).

| Parameter | Call |
|---|---|
| Finitial | 31.38 ±1.67 (30.20-32.57) |
| Ffinal | 19.28 ±0.32 (19.04-19.51) |
| Fmin | 24.43 ±1.14 (23.62-25.25) |
| Fmax | 21.98 ±0.54 (21.60-22.37) |
| FME | 28.14 ±0.76 (27.60-28.68) |
| BW | 6.15 ±0.22 (6.00-6.31) |
| Dur | 0.010 ±0.001 (0.009-0.011) |
| IC | 0.189 ±0.030 (0.167-0.211) |

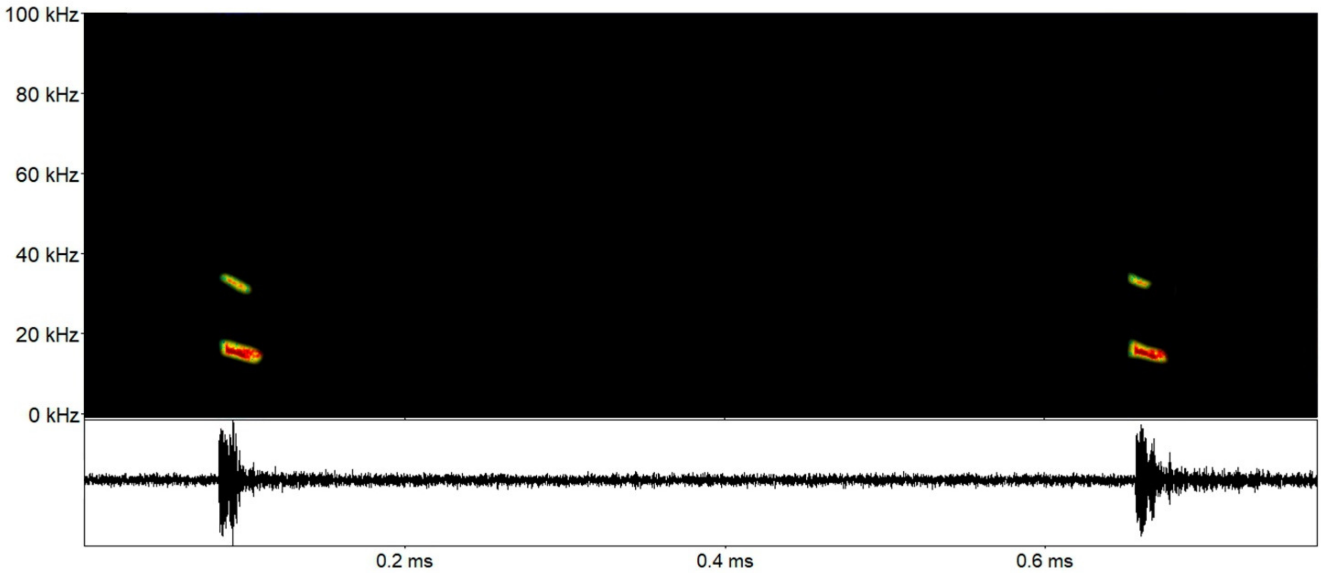

***Eumops* sp. 1**: Sequencies with a FM descendent call and FME around 15.8 kHz (15.3-16.1 kHz).

| Parameter | Call |
|---|---|
| Finitial | 12.52 ±0.37(12.14-13.12) |
| Ffinal | 18.21 ±0.43 (17.82-18.90) |
| Fmin | 15.25 ±0.38 (14.62-15.75) |
| Fmax | 14.18 ±0.28 (13.87-14.62) |
| FME | 15.81 ±0.36 (15.37-16.12) |
| BW | 1.62 ±0.19 (1.50-1.87) |
| Dur | 0.014 ±0.001 (0.012-0.015) |
| IC | 0.664 ±0.016 (0.651-0.692) |

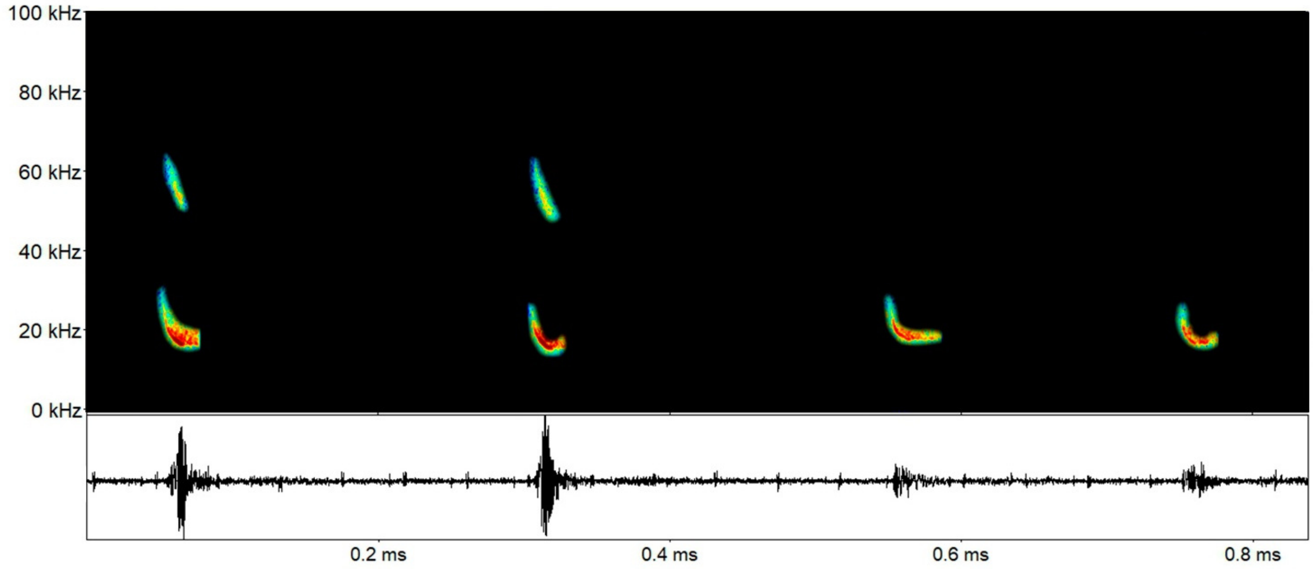

***Eumops* cf. *dabbenei*** Thomas, 1914: Sequencies with two FM descendent calls with slightly different frequencies (type a, lower; type b, higher). FME around 18.8 kHz for type a (18.1-19.5 kHz), and 21.3 kHz for type b (20.1-22.6 kHz).

| Parameter | Call a | Call b |
|---|---|---|
| Finitial | 21.26 ±1.38 (19.54-22.88) | 23.00 ±1.47 (21.96-24.04) |
| Ffinal | 14.69 ±1.19 (13.37-15.82) | 17.52 ±2.10 (16.03-19.01) |
| Fmin | 16.90 ±0.87 (15.67-17.93) | 19.42 ±1.87 (18.09-20.75) |
| Fmax | 15.66 ±0.71 (14.85-16.50) | 18.09 ±1.98 (16.68-19.50) |
| FME | 18.86 ±0.55 (18.18-19.50) | 21.39 ±1.74 (20.15-22.62) |
| BW | 3.20 ±0.57(2.345-3.75) | 3.29 ±0.24 (3.12-3.46) |
| Dur | 0.021 ±0.004 (0.014-0.025) | 0.017 ±0.003 (0.015-0.019) |
| IPI | 0.124 ±0.063 (0.079-0.169) | - |
| IC | 0.552 ±0.488 (0.098-1.380) | - |

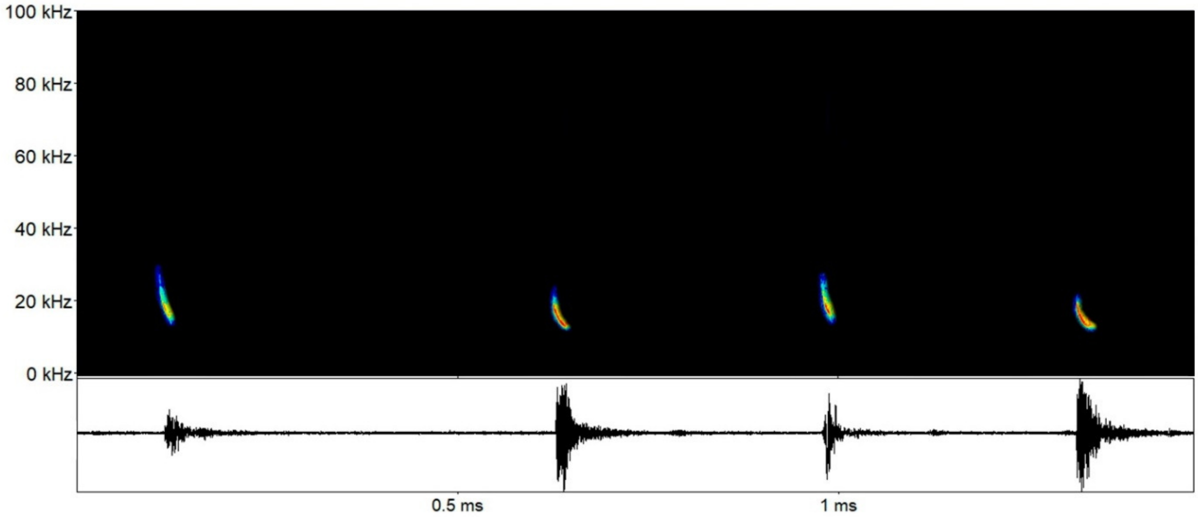

***Eumops* spp. (*Eumops hansae* or *Eumops maurus*)**: Sequencies with two FM descendent calls (type a, lower; type b, higher), with harmonics. FME around 21.8 kHz (20.8-23.3 kHz) for type a, and around 23.9 kHz (22.5-25.5 kHz) for type b.

| Parameter | Call a | Call b |
|---|---|---|
| Finitial | 27.00 ±3.60 (23.59-34.34) | 27.07 ±3.69 (24.46-29.68) |
| Ffinal | 16.72 ±1.14 (15.32-18.94) | 20.53 ±2.52 (18.75-22.32) |
| Fmin | 19.64 ±1.09 (18.52-21.3) | 22.12 ±2.12 (20.62-23.62) |
| Fmax | 18.36 ±1.46 (16.46-20.47) | 21.31 ±2.56 (19.50-23.12) |
| FME | 21.83 ±0.84 (20.81-23.32) | 23.93 ±2.03 (22.50-25.37) |
| BW | 3.46 ±1.08 (2.15-5.66) | 2.62 ±0.53 (2.25-3.00) |
| Dur | 0.021 ±0.005 (0.012-0.029) | 0.019 ±0.0004 (0.0190-0.0197) |
| IPI | 0.128 ±0.011 (0.120-0.135) | - |
| IC | 0.370 ±0.072 (0.307-0.474) | - |

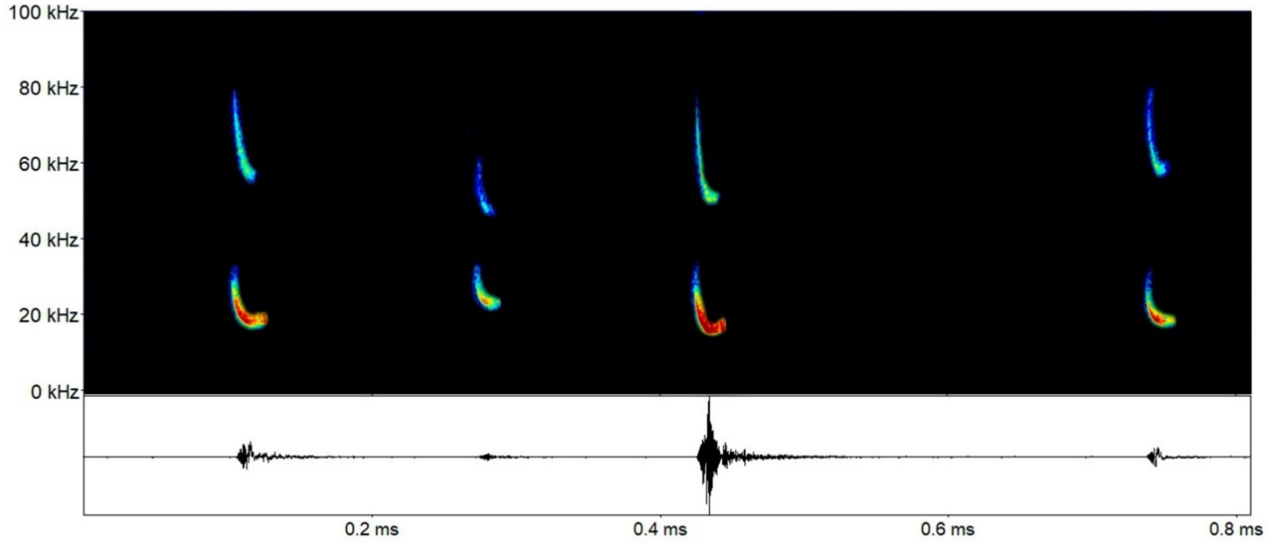

***Molossops neglectus*** Williams e Genoways, 1980: Sequencies composed by two alternating FM ascendent calls, being type a calls with a broader frequency variation. FME for type a calls around 34.5 kHz (25.5-39.3 kHz), and around 40.4 kHz (39.3-41.2 kHz) for type b.

| Parameter | Call a | Call b |
|---|---|---|
| Finitial | 26.80 ±1.73 (25.22-29.83) | 34.98 ±1.41 (32.92-36.18) |
| Ffinal | 42.84 ±6.46 (28.20-45.55) | 45.86 ±0.54 (45.23-46.64) |
| Fmin | 31.55 ±2.80 (25.50-33.37) | 38.77 ±0.94 (37.12-39.37) |
| Fmax | 38.94 ±4.91 (28.12-42.00) | 44.25 ±0.79 (43.12-45.00) |
| FME | 34.55 ±4.59 (25.50-39.37) | 40.42 ±0.72 (39.37-41.25) |
| BW | 7.39 ±2.18 (2.62-9.00) | 5.47 ±1.44 (4.12-7.87) |
| Dur | 0.013 ±0.004 (0.002-0.016) | 0.011 ±0.001 (0.01-0.014) |
| IPI | 0.103 ±0.011 (0.091-0.121) | - |
| IC | 0.111 ±0.014 (0.095-0.132) | - |

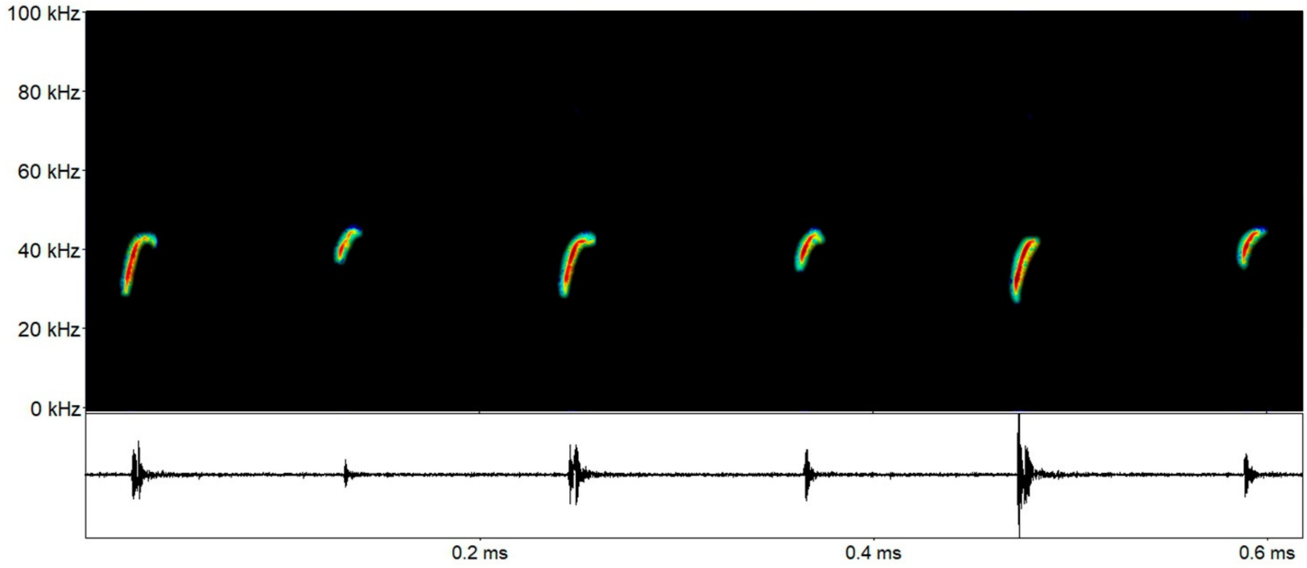

***Molossus* cf. *currentium*** Thomas, 1901: Sequencies composed by two alternating qCF descendent calls, being type a calls with lower average frequency, and higher in type b calls. FME for type a calls around 29.6 kHz (29.2-30.1 kHz), and around 34.5 kHz for type b.

| Parameter | Call a | Call b |
|---|---|---|
| Finitial | 24.28 ±0.11 (24.20-24.36) | 30.83 ±1.17 (30.00-31.66) |
| Ffinal | 32.13 ±1.18 (31.29-32.96) | 37.35 ±0.68 (36.86-37.83) |
| Fmin | 26.45 ±0.84 (25.85-27.05) | 32.87 ±0.09 (32.81-32.94) |
| Fmax | 30.51 ±0.78 (29.95-31.07) | 35.68 ±0.006 (35.67-35.68) |
| FME | 29.69 ±0.62 (29.25-30.14) | 34.52 ±0.03 (34.50-34.55) |
| BW | 4.06 ±0.06 (4.01-4.10) | 2.80 ±0.10(2.73-2.87) |
| Dur | 0.018 ±0.003 (0.015-0.020) | 0.018 ±0.002 (0.016-0.019) |
| IPI | 0.08 ±0.006 (0.075-0.084) | - |
| IC | 0.174 ±0.024 (0.156-0.191) | - |

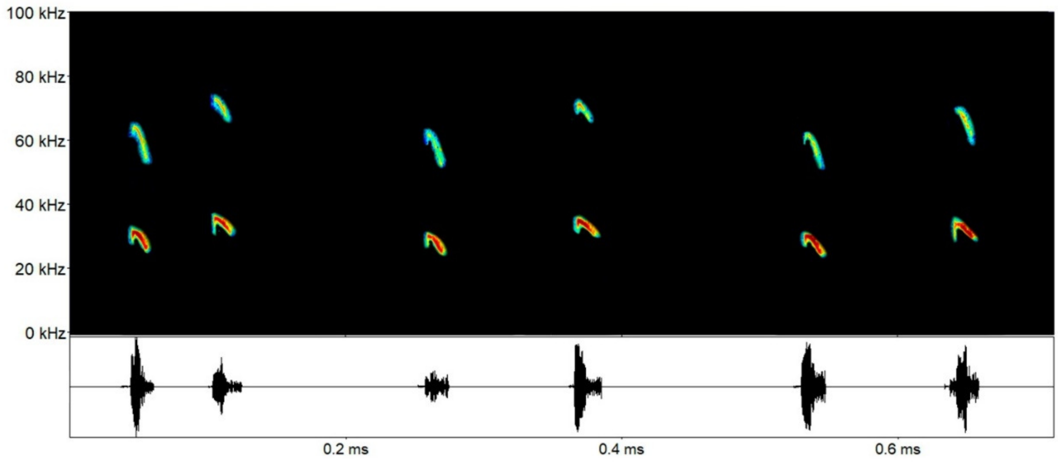

***Molossus molossus*** Pallas, 1766: Sequencies composed by two – sometimes three – alternating qCF descendent calls. FME around 33.9 kHz for type a calls (28.8-35.8 kHz), 38.9 kHz for type b calls (35.2-42.0 kHz), and 43.3 kHz for type c calls (38.6-48.0 kHz).

| Parameter | Call a | Call b | Call c |
|---|---|---|---|
| Finitial | 29.98 ±3.05 (24.61-31.96) | 35.16 ±2.60 (30.97-37.31) | 38.67 ±3.85 (35.95-41.40) |
| Ffinal | 36.48 ±1.97 (33.13-38.42) | 41.60 ±2.94 (38.09-46.10) | 47.05 ±7.80 (41.53-52.57) |
| Fmin | 32.47 ±3.11 (26.93-34.38) | 37.82 ±2.48 (33.82-40.37) | 41.34 ±5.43 (37.50-45.18) |
| Fmax | 34.73 ±2.39 (30.56-36.67) | 39.69 ±2.78 (35.85-43.41) | 44.95 ±7.88 (39.37-50.53) |
| FME | 33.92 ±2.85 (28.87-35.88) | 38.99 ±2.55(35.25-42.08) | 43.35 ±6.69 (38.62-48.09) |
| BW | 2.26 ±0.79 (1.60-3.62) | 1.875 ±0.72 (1.12-3.04) | 3.60 ±2.45 (1.87-5.34) |
| Dur | 0.015 ±0.005 (0.008-0.024) | 0.012 ±0.002 (0.008-0.014) | 0.009 ±0.002 (0.007-0.011) |
| IPI | 0.065 ±0.019 (0.044-0.091) | - | - |
| IC | 0.111 ±0.027 (0.071-0.143) | - | - |

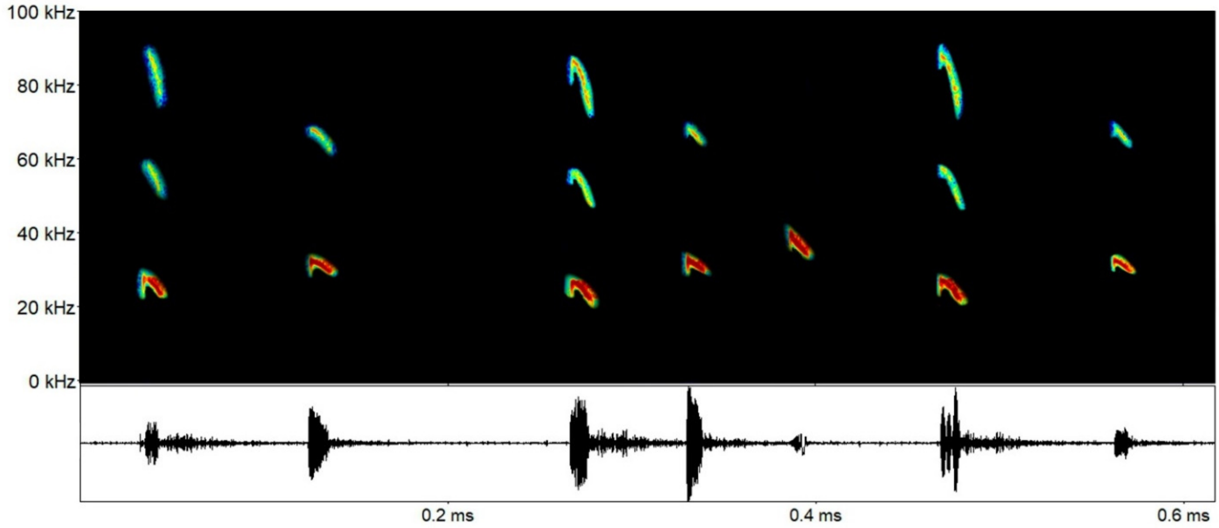

***Molossus rufus*** É. Geoffroy, 1805: Sequencies composed by two or three low qCF descendent calls, with the first being the stronger one. FME around 25.9 kHz for type a calls (24.7-27.4 kHz), 29.5 kHz for type b calls (28.5-30.6 kHz), and 31.8 kHz for type c (30.7-34.1 kHz).

| Parameter | Call a | Call b | Call c |
|---|---|---|---|
| Finitial | 21.14 ±1.71 (19.49-22.91) | 26.49 ±1.77 (24.57-28.08) | 29.27 ±0.84 (28.53-30.51) |
| Ffinal | 28.65 ±0.77 (28.06-29.53) | 31.63 ±0.68 (31.15-32.41) | 33.46 ±1.75 (31.97-36.80) |
| Fmin | 24.45 ±0.95 (23.71-25.53) | 28.42 ±1.09 (27.32-29.51) | 30.81 ±0.90 (30.00-32.25) |
| Fmax | 27.21 ±1.04 (26.55-28.42) | 30.46 ±0.99 (29.57-31.53) | 32.18 ±1.28 (30.75-34.50 |
| FME | 25.91 ±1.35 (24.79-27.41) | 29.51 ±1.03 (28.59-30.63) | 31.81 ±1.24 (30.75-34.12) |
| BW | 2.75 ±0.28 (2.43-2.95) | 2.04 ±0.20 (1.85-2.25) | 1.37 ±0.51 (0.75-2.25) |
| Dur | 0.016 ±0.003 (0.012-0.019) | 0.015 ±0.002 (0.012-0.016) | 0.013 ±0.002 (0.01-0.018) |
| IPI | 0.076 ±0.025 (0.047-0.095) | | |
| IC | 0.175 ±0.026 (0.145-0.191) | | |

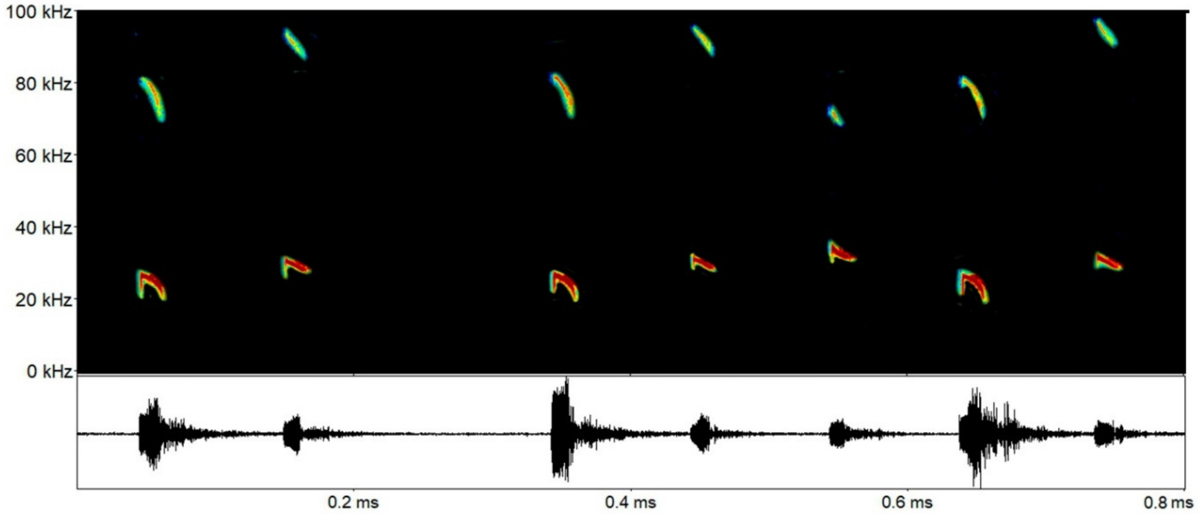

***Promops centralis*** Thomas, 1915: Sequencies may present very distinctive and different calls (Arias-Aguilar et al. 2018; Hintze et al. 2019). In our analysis we detected only the lazy s-shape calls, with FME around 26.8 kHz (23.2-29.6 kHz).

| Parameter | Call |
|---|---|
| Finitial | 22.82 ±1.14 (20.70-24.73) |
| Ffinal | 31.54 ±0.63 (30.38-32.79) |
| Fmin | 24.79 ±1.05 (22.50-25.87) |
| Fmax | 29.70 ±1.13 (26.62-31.12) |
| FME | 26.89 ±1.89 (23.25-29.62) |
| BW | 4.91 ±0.86 (3.75-6.37) |
| Dur | 0.06 ±0.025 (0.024-0.106) |
| IC | 0.14 ±0.064 (0.002-0.287) |

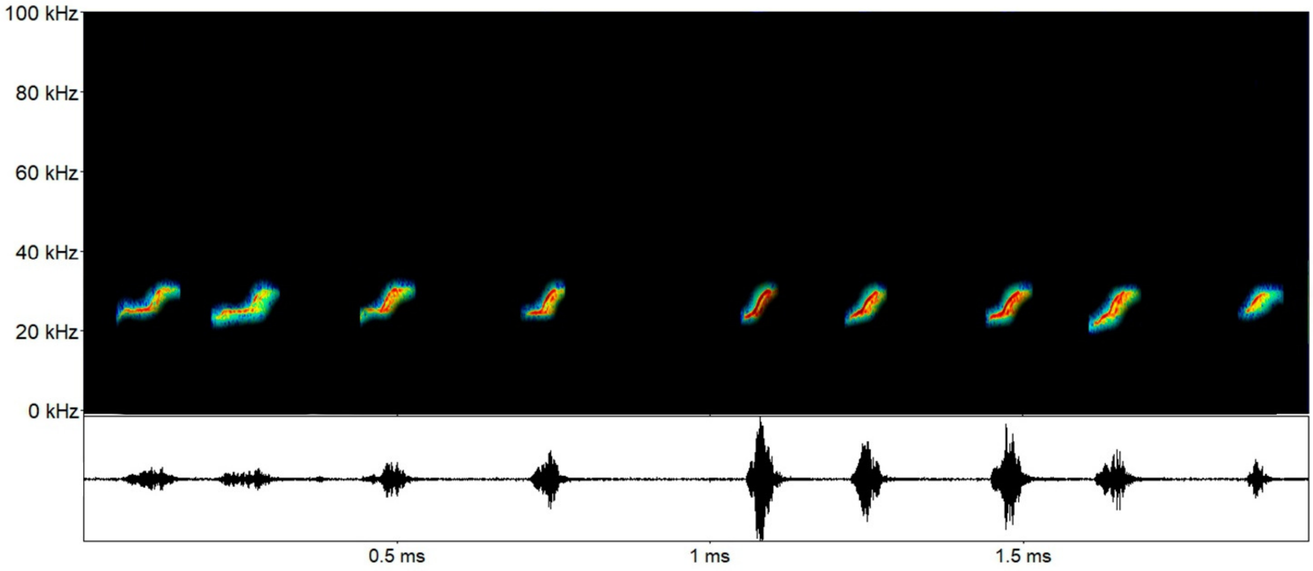

### Family Vespertilionidae

Echolocation calls for *Eptesicus*/*Lasiurus* sp. 1 (EpLasp1), *Eptesicus*/*Lasiurus* sp. 2 (EpLasp2), *Eptesicus*/*Lasiurus* sp. 3 (EpLasp3), *Eptesicus*/*Lasiurus* sp. 4 (EpLasp4), *Eptesicus*/*Lasiurus* sp. 5 (EpLasp5), *Myotis* sp. 1 (Mysp1), *Myotis* sp. 2 (Mysp2), *Myotis* sp. 3 (Mysp3), and *Myotis* sp. 4 (Mysp4) in the Carajás region, Brazilian Amazonia.

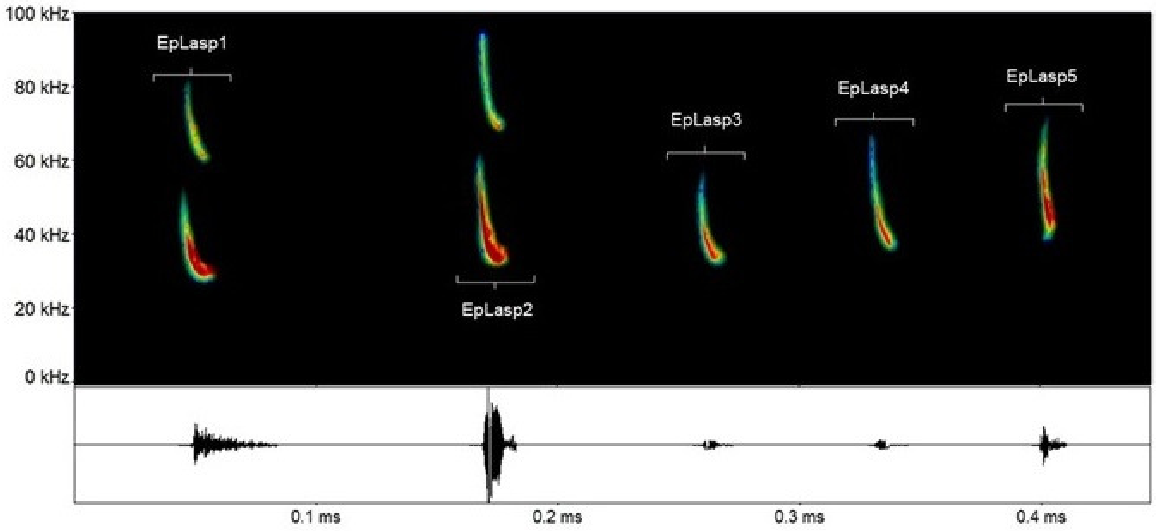


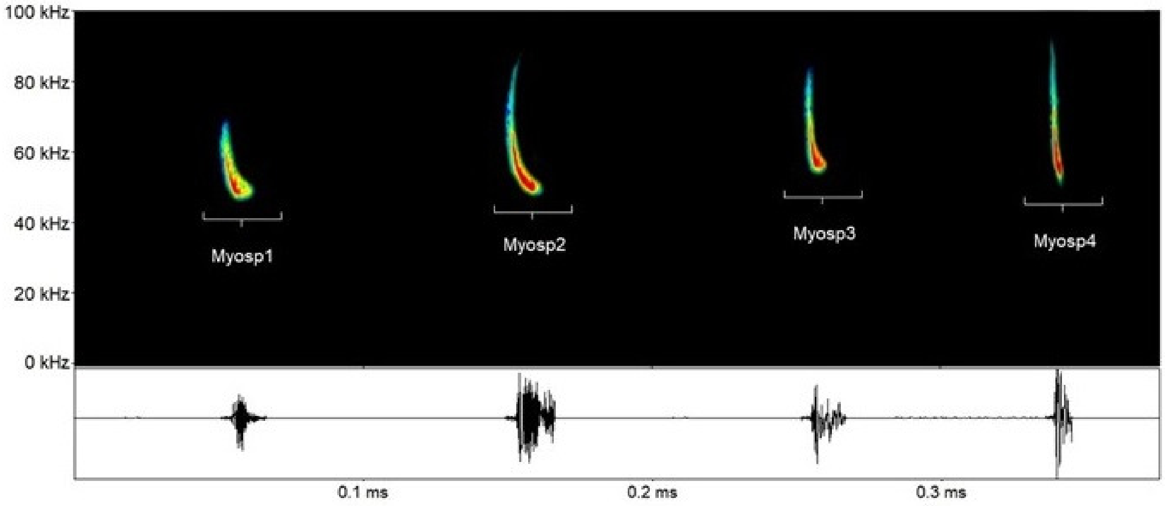

***Eptesicus/Lasiurus* sp. 1**: Sequencies with broad descendent FM calls, with a rounded inflection point and harmonics, without frequency alternance. FME around 34.4 kHz (32.4-36.9 kHz) and lower frequencies for the first harmonic – which are more useful for sonotype/species differentiation among vespertilionids – around 29.0 kHz (27.1-29.9 kHz).

| Parameter | Call |
|---|---|
| Finitial | 26.92 ±1.43 (25.14-28.40) |
| Ffinal | 43.60 ±6.25 (39.38-54.46) |
| Fmin | 29.07 ±1.15 (27.14-29.96) |
| Fmax | 28.07 ±1.28 (26.43-29.28) |
| FME | 34.46±1.78 (32.48-36.97) |
| BW | 6.39 ±2.03(4.46-8.57) |
| Dur | 0.018 ±0.003 (0.013-0.021) |
| IC | 0.214 ±0.055 (0.123-0.256) |

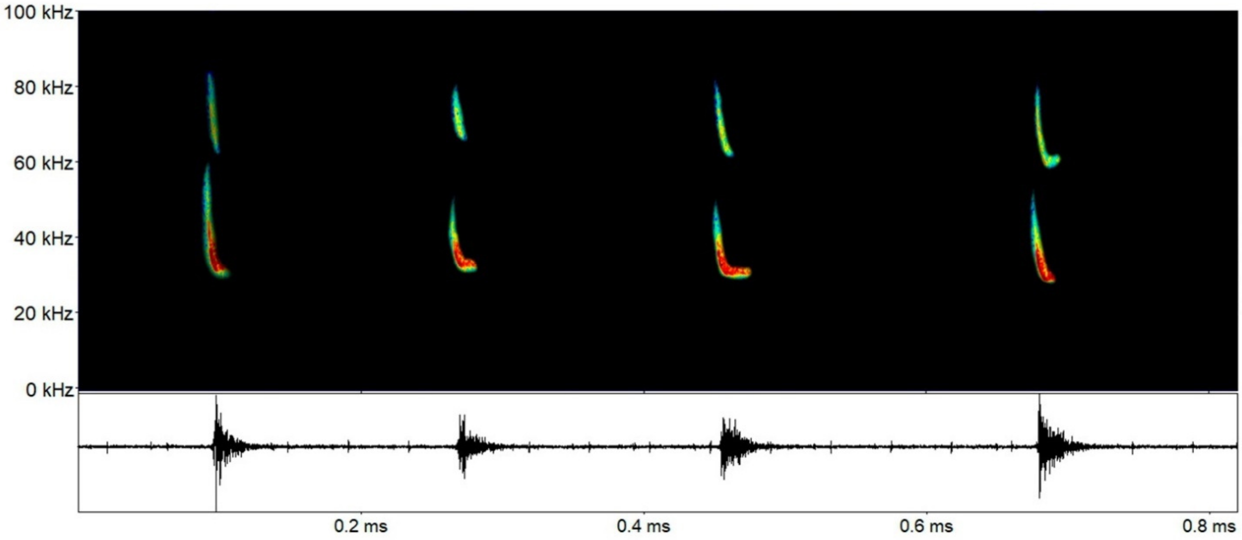

***Eptesicus/Lasiurus* sp. 2**: Sequencies with broad descendent FM calls, with a rounded inflection point and harmonics, without frequency alternance. FME around 39.4 kHz (34.2-51.1 kHz) and lower frequencies for the first harmonic – which are more useful for sonotype/species differentiation among vespertilionids – around 32.5 kHz (30.56-34.95 kHz).

| Parameter | Call |
|---|---|
| Finitial | 30.50 ±1.44 (28.30-32.18) |
| Ffinal | 48.40 ±6.96 (39.31-57.39) |
| Fmin | 32.57 ±1.65 (30.56-34.95) |
| Fmax | 31.66 ±1.50 (29.77-33.75) |
| FME | 39.43 ±5.64 (34.27-51.15) |
| BW | 7.76 ±4.56(3.82-8.06) |
| Dur | 0.020 ±0.015 (0.007-0.052) |
| IC | 0.165 ±0.056 (0.084-0.246) |

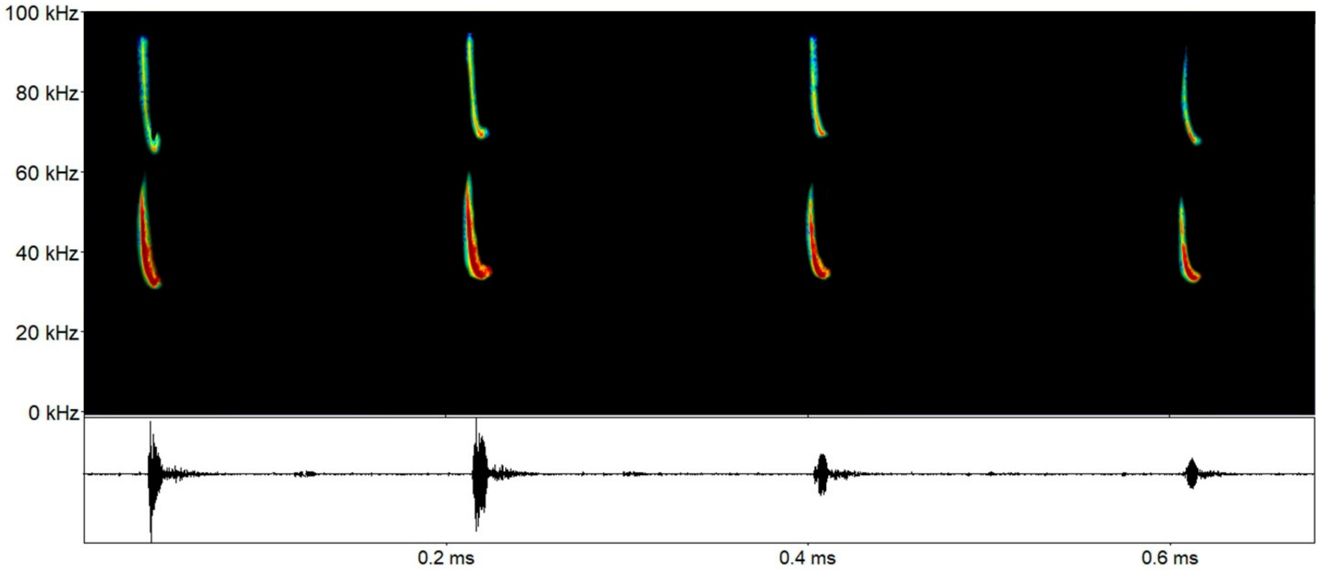

***Eptesicus/Lasiurus* sp. 3**: Sequencies with broad descendent FM calls, with a rounded inflection point and harmonics, without frequency alternance. FME around 42.1 kHz (39.7-46.9 kHz) and lower frequencies for the first harmonic – which are more useful for sonotype/species differentiation among vespertilionids – around 36.7 kHz (35.5-38.3 kHz).

| Parameter | Call |
|---|---|
| Finitial | 33.97 ±0.82 (32.79-35.07) |
| Ffinal | 51.46 ±5.60 (44.66-61.82) |
| Fmin | 36.77 ±1.06 (35.56-38.37) |
| Fmax | 35.38 ±0.65 (34.56-36.37) |
| FME | 42.10 ±2.70 (39.71-46.91) |
| BW | 6.71 ±2.60 (3.90-7.65) |
| Dur | 0.016 ±0.009 (0.005-0.031) |
| IC | 0.163 ±0.053 (0.094-0.232) |

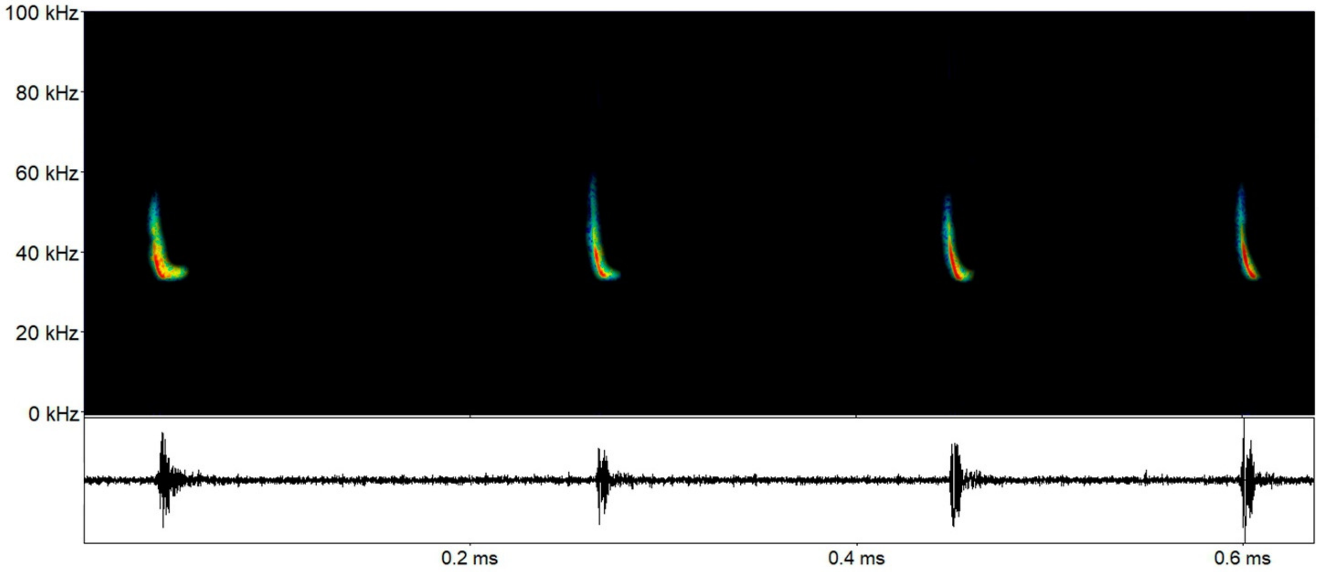

***Eptesicus/Lasiurus* sp. 4**: Sequencies with broad descendent FM calls, but without the rounded inflection point and harmonics like in the previous sonotypes, and no frequency alternance. FME around 45.3 kHz (33.7-54.1 kHz) and lower frequencies for the first harmonic – which are more useful for sonotype/species differentiation among vespertilionids – around 38.3 kHz (38.0-40.6 kHz).

| Parameter | Call |
|---|---|
| Finitial | 35.37 ±2.76 (34.08-37.01) |
| Ffinal | 59.79 ±9.46 (50.99-71.69) |
| Fmin | 38.30 ±3.23 (38.08-40.66) |
| Fmax | 36.88 ±2.92 (37.01-38.85) |
| FME | 45.38 ±5.54 (33.75-54.10) |
| BW | 8.49 ±3.66(5.14-10.53) |
| Dur | 0.012 ±0.004 (0.007-0.018) |
| IC | 0.113 ±0.026 (0.089-0.176) |

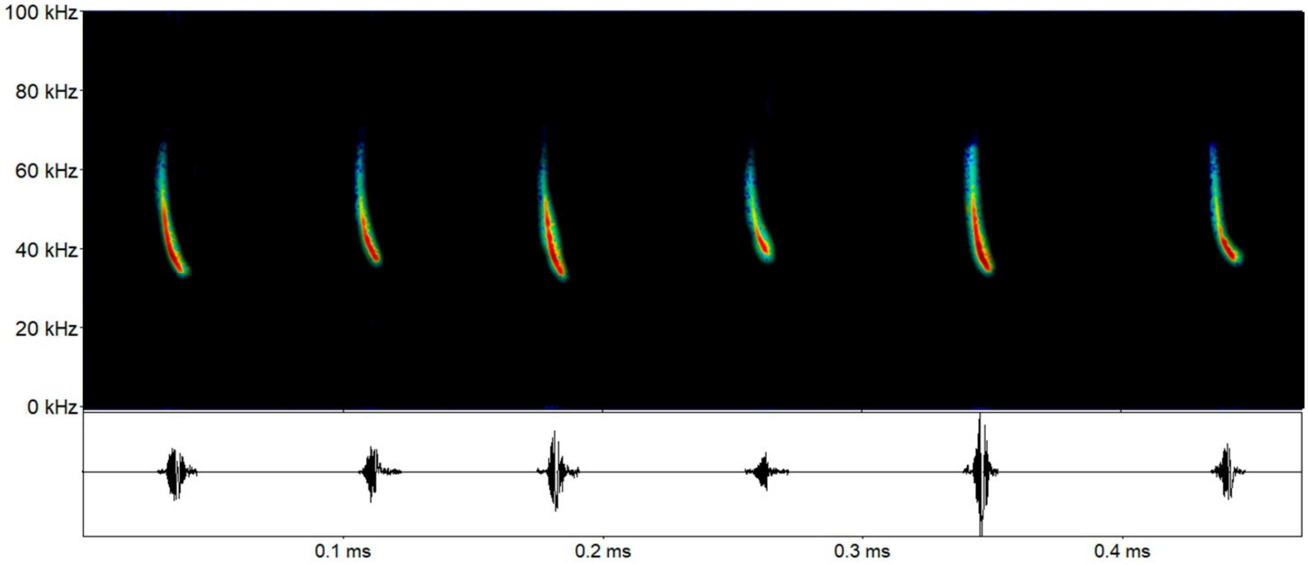

***Eptesicus/ Lasiurus* sp. 5**: Sequencies with broad descendent FM calls, but without the rounded inflection point and harmonics like in the previous sonotypes, and no frequency alternance. FME around 52.5 kHz (47.5-55.7 kHz) and lower frequencies for the first harmonic – which are more useful for sonotype/species differentiation among vespertilionids – around 44.7 kHz (43.2-46.6 kHz).

| Parameter | Call |
|---|---|
| Finitial | 40.97 ±1.33 (39.39-42.34) |
| Ffinal | 62.88 ±11.24 (52.44-67.68) |
| Fmin | 44.73 ±1.60 (43.28-46.46) |
| Fmax | 42.98 ±0.68 (42.26-43.95) |
| FME | 52.53 ±4.01 (47.55-55.76) |
| BW | 9.55 ±3.92(4.20-12.75) |
| Dur | 0.01 ±0.003 (0.007-0.014) |
| IC | 0.122 ±0.058 (0.069-0.213) |

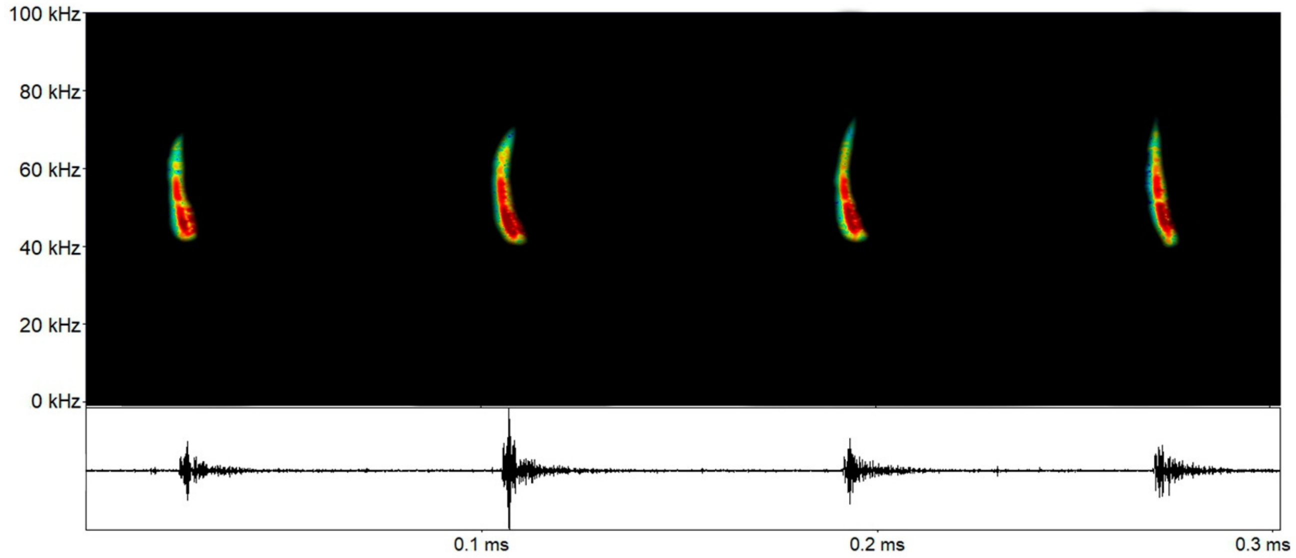

***Myotis* sp. 1**: Sequencies with broad descendent FM calls, with a rounded inflexion point, but no visible harmonics. FME around 54.4 kHz (52.7-55.9 kHz), and lower frequencies around 49.3 kHz (48.4-50.3 kHz).

| Parameter | Call |
|---|---|
| Finitial | 47.06 ±1.00 (46.02-48.57) |
| Ffinal | 64.73 ±3.71 (64.97-68.33) |
| Fmin | 49.35 ±0.74 (48.41-50.30) |
| Fmax | 48.40 ±0.84 (47.36-49.71) |
| FME | 54.41 ±1.56 (52.73-55.96) |
| BW | 6.00 ±1.41 (4.45-7.68) |
| Dur | 0.011 ±0.001 (0.009-0.013) |
| IC | 0.116 ± 0.014 (0.101-0.134) |

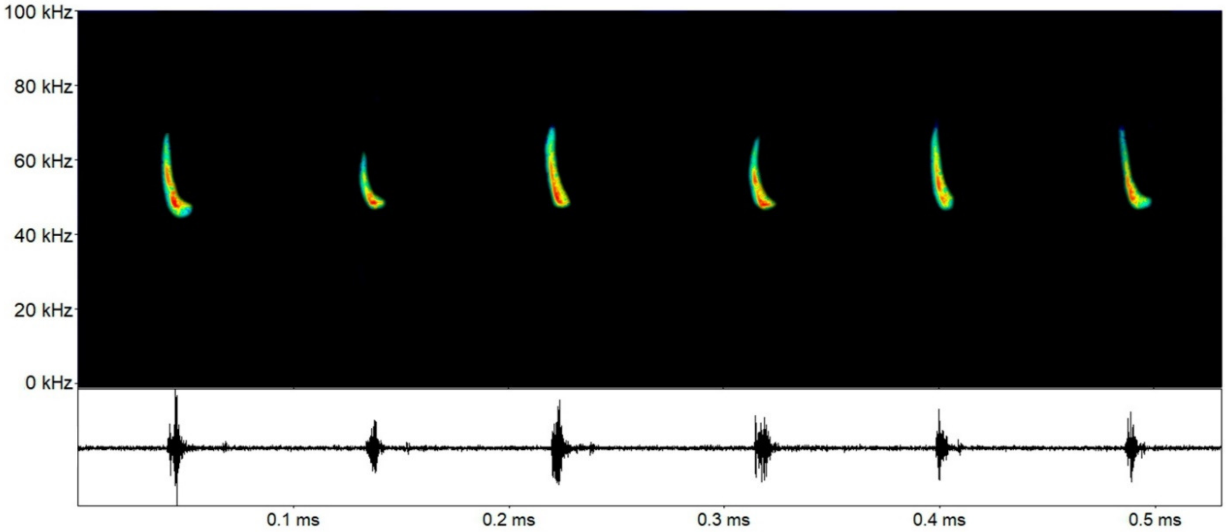

***Myotis* sp. 2**: Sequencies with broad descendent FM calls, with a not so evidente rounded inflexion point. FME around 60.0 kHz (55.6-63.3 kHz), and lower frequencies around 53.3 kHz (51.0-54.2 kHz).

| Parameter | Call |
|---|---|
| Finitial | 50.16 ±1.11 (48.62-51.63) |
| Ffinal | 70.81 ±5.54 (64.25-80.42) |
| Fmin | 53.33 ±1.07 (51.07-54.28) |
| Fmax | 51.87 ±0.85 (50.55-52.87) |
| FME | 60.07 ±2.59 (55.61-63.30) |
| BW | 8.19 ±1.91 (5.06-10.59) |
| Dur | 0.009 ±0.002 (0.005-0.012) |
| IC | 0.128 ±0.025 (0.086-0.157) |

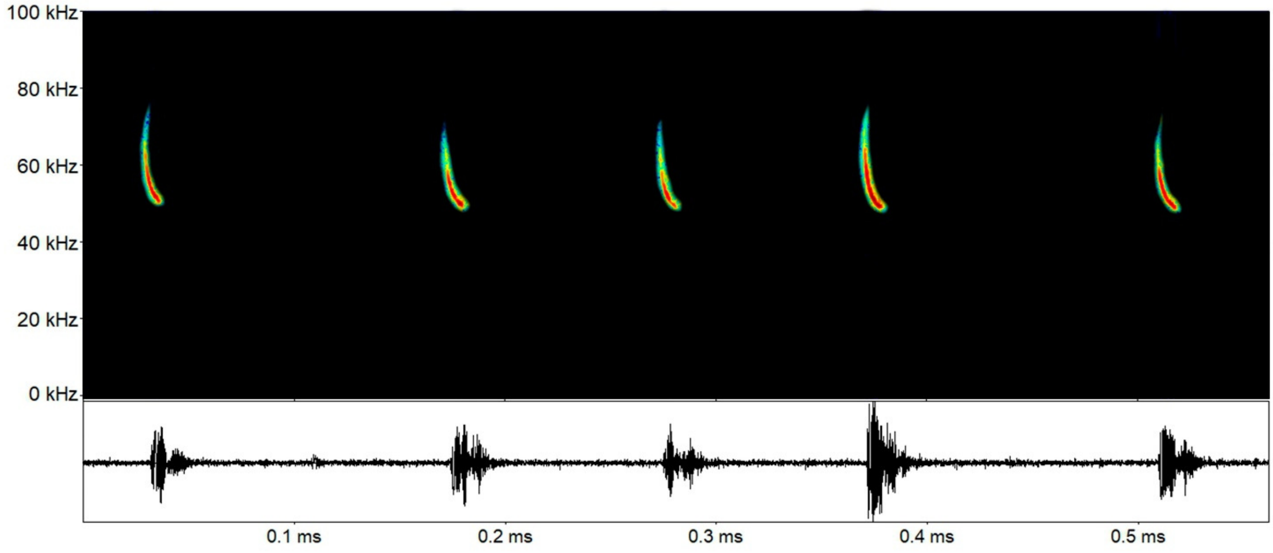

***Myotis* sp. 3**: Sequencies with broad descendent FM calls, with an evidente rounded inflexion point. FME around 62.1 kHz (59.8-64.7 kHz), and lower frequencies around 55.7 kHz (53.6-57.2 kHz).

| Parameter | Call |
|---|---|
| Finitial | 53.27 ±0.84 (52.08-54.57) |
| Ffinal | 72.21 ±3.24 (69.36-76.83) |
| Fmin | 55.70 ±1.18 (53.62-57.26) |
| Fmax | 54.72 ±1.06 (53.34-55.98) |
| FME | 62.16 ±1.55 (59.81-64.75) |
| BW | 7.44 ±1.54 (4.12-8.62) |
| Dur | 0.010 ±0.003 (0.005-0.016) |
| IC | 0.098 ±0.057 (0.044-0.194) |

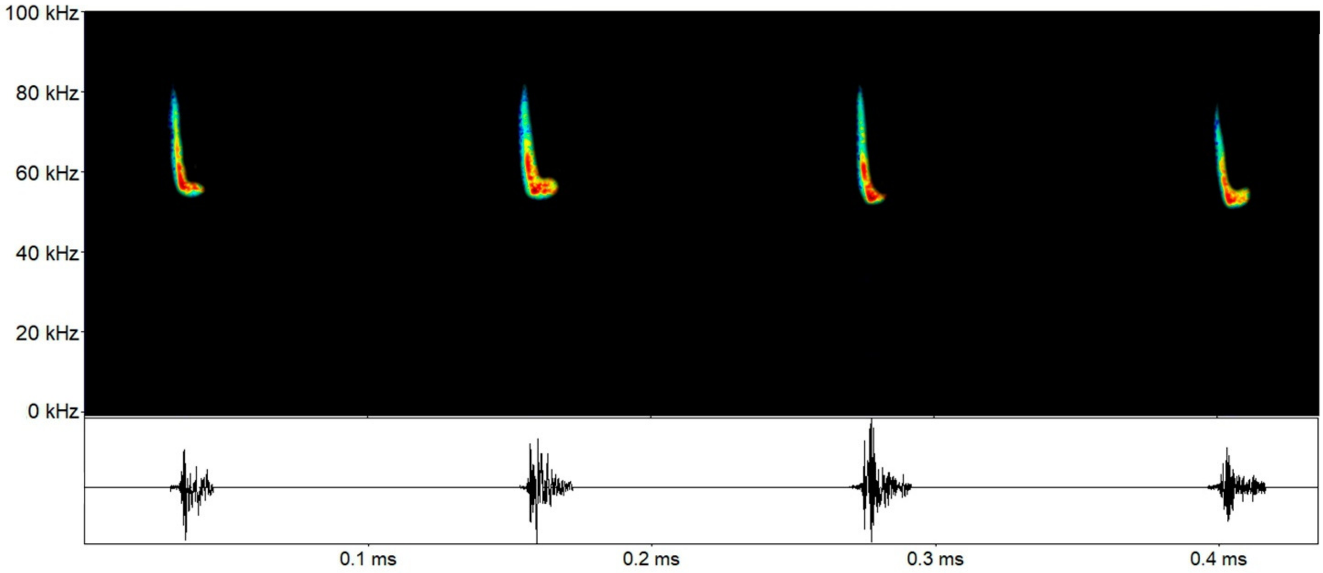

***Myotis* sp. 4**: Sequencies with broad descendent FM calls, with no rounded inflexion point. FME around 66.1 kHz (63.8-68.6 kHz), and lower frequencies around 59.7 kHz (58.5-61.5 kHz).

| Parameter | Call |
|---|---|
| Finitial | 56.87 ±1.69 (55.06-59.14) |
| Ffinal | 81.54 ±5.96 (74.77-89.17) |
| Fmin | 59.78 ±1.35 (58.50-61.59) |
| Fmax | 58.60 ±1.29 (57.37-60.37) |
| FME | 66.13 ±1.97 (63.80-68.62) |
| BW | 7.52 ±0.87 (6.42-8.25) |
| Dur | 0.008 ±0.002 (0.005-0.011) |
| IC | 0.098 ±0.046 (0.043-0.144) |

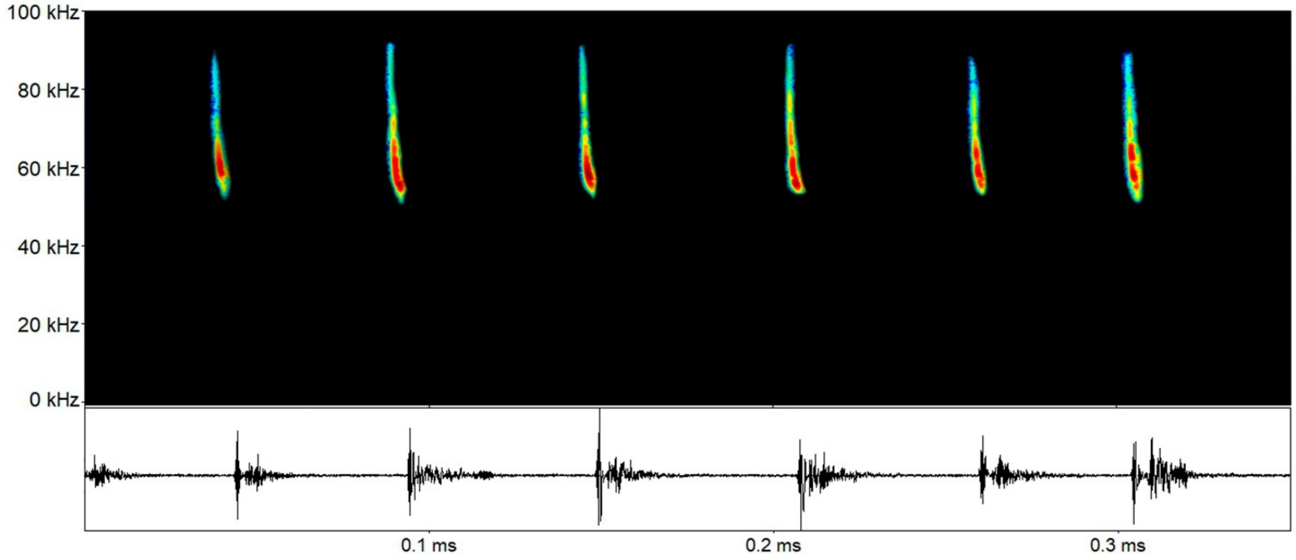
